# Neural speech tracking in newborns: prenatal learning and contributing factors

**DOI:** 10.1101/2024.03.18.585222

**Authors:** Cristina Florea, Michaela Reimann, Fabian Schmidt, Jasmin Preiß, Eva Reisenberger, Monika Angerer, Mohamed Ameen, Dominik Heib, Dietmar Roehm, Manuel Schabus

## Abstract

**Introduction:** Early language development in infants is being increasingly studied, though only recently with direct measurements of brain activity rather than with behavioral or physiological measurements. In the current study, we use electroencephalographic (EEG) recordings of 2-week-old infants to look for signs of prenatal learning and to investigate newborns’ abilities to process language. We also look at the influence of prenatal stress factors and at the predictive value of the newborns’ language processing abilities for later language development.

**Methods:** Sixty pregnant women played a rhyme to their abdomen twice a day from the 34^th^ week of pregnancy until birth, to familiarize the fetus with the rhyme. At around 2 weeks after delivery (mean age 16 days), the newborns were exposed to the familiar rhyme as well as to an unfamiliar one while their EEG was recorded. Additionally, three manipulations of the familiar rhyme were played: (1) low-pass filtered, (2) with changed rhythm, and (3) inverted and played backwards. The data was analyzed to see how well the infant brain signal followed the speech envelope in each condition.

Accounting for the heterogenous approach used for neural speech tracking in the literature, we used four methods, namely: (1) coherence, (2) Hilbert coherence, (3) temporal response functions (TRF), and (4) mutual information (MI). The maternal prenatal depression was evaluated with Edinburgh Prenatal Depression Score and the chronic fetal stress was measured from the hair cortisol levels of the 2 week-olds. The language development at 6 months of age was evaluated with the Bayley Scales.

**Results and discussion:** Overall, the results indicate the presence of prenatal learning, with the unfamiliar rhyme eliciting stronger cortical tracking (higher coherence and MI) than the familiar rhyme, which suggests stronger brain-to-speech coupling for the unfamiliar rhyme, perhaps deriving from more effort to process the unexpected stimulus. However, the original version of the familiar rhyme proved to be the easiest to track compared to the language- and rhythm-manipulations, (higher MI for the original rhyme than the language manipulation and higher coherence and mTRF correlation coefficients for the original rhyme than the rhythm manipulation). This indicates language discrimination and a prosodic-based learning of the familiar rhyme. Furthermore, there is an indication of phonotactic sensitivity at this young age, with less tracking (lower Hilbert coherence and lower mTRF correlation coefficients) of the low-pass filtered rhyme than the original version, indicating that the phonological cues erased by the filtering were important for the newborn’s ability to follow the rhyme.

Furthermore, the mothers’ depression scores positively correlated with the infant’s tracking ability for the familiar rhyme. This suggests that a slightly lower mood was more stimulative for the fetal language development. The chronic fetal stress levels, however, were negatively correlated with the cortical tracking abilities. Importantly, the newborn’s cortical tracking was positively correlated with the infant’s language development at 6 months of age, underlining the predictive value of the early assessment of language processing.

**Conclusion:** Prenatal learning is well established, but evidence including (healthy) brain data in the first weeks of life is scarce. The current study shows that newborns can discriminate between a familiar and unfamiliar rhyme, while also highlighting the role of prosody in early language processing, and bringing new evidence of their sensitivity to phonotactic cues in auditory stimuli. Furthermore, the newborn’s ability to track a rhyme is correlated with their language development at 6 months. The newborn’s cortical tracking of the familiar rhyme is further increased by moderately low maternal mood, but decreased by fetal stress. Future studies with similar fine-grained linguistic designs but of older infants should teach us the timeline of what exactly is learned prenatally and at very early age in respect to language.

## Introduction

The development of perceptive language skills and the familiarization with the mother tongue starts before birth (Dehaene-Lambertz & Spelke, 2015) and it strongly depends on environmental factors, i.e., auditory input (Webb et al., 2015). The sounds to which a fetus is exposed during gestation shape their ability to discriminate the mother’s voice from a stranger’s (Beauchemin et al., 2011; DeCasper & Fifer, 1980; Kisilevsky et al., 2003, 2009; Picciolini et al., 2014; Rand & Lahav, 2014), the maternal language from a foreign one (Abboub et al., 2016; Kisilevsky et al., 2009; Minai et al., 2017), and even influences the prosody of their cries after birth (Mampe et al., 2009).

### Newborns can track continuous speech

On fetuses and newborns, most language processing studies measure indirect behavioral or physiological indicators like fetal heart rate or movement (Kisilevsky et al., 2009; Kisilevsky & Hains, 2011), or, for infants, head turn (Thiessen et al., 2005; Trainor & Desjardins, 2002) or suckling frequency (DeCasper & Spence, 1986; Moon et al., 2013). More recently, studies started to acquire direct brain data, using either electroencephalography (EEG) or near-infrared spectroscopy (NIRS), but most often the auditory stimuli used are short, designed to evoke an event-related potential (ERP, in the EEG method: Partanen, Kujala, Näätänen, et al., 2013a), or rapid changes in the concentrations of oxygenated and deoxygenated hemoglobin (in the NIRS method: Abboub et al., 2016; Gervain et al., 2008). Even so, reports with direct brain data within the first month of life in at-term infants are scarce (Abboub et al., 2016; Gervain et al., 2008; Ortiz Barajas et al., 2021; Partanen, Kujala, Näätänen, et al., 2013a), and very few investigate the tracking of longer, continuous language stimuli (Ortiz Barajas et al., 2021). However, even if not in newborns, in older infants (7-14 months) continuous speech tracking has been investigated by some studies (Kalashnikova et al., 2018; Menn, Michel, et al., 2022; Menn, Ward, et al., 2022). Continuous speech stimuli allow the analysis of infants’ language processing abilities in a more naturalistic situation, as speech is a complex stimulus with many variable attributes (rhythm, pitch, volume, etc.) that short auditory stimuli cannot always reproduce. Following from these studies, we expected that cortical tracking of continuous speech is also detectable in newborns.

### Cortical tracking of the familiar language is better than that of unintelligible language

Discrimination of the maternal language from a foreign one has been shown previously, e.g. by Kisilevsky et al., 2009, who found that the fetal heart rate increases when a female stranger changed the language from familiar (English) to unfamiliar (Mandarin). Another study on fetuses (Minai et al., 2017), using a bilingual English-Japanese unfamiliar female speaker, showed that fetuses (with English as their mother-tongue) specifically discriminate the two languages, having a novelty reaction with faster heart rate to the unfamiliar language. However, language discrimination is only possible if the two languages have different rhythms (Nazzi et al., 1998). Evidence from brain data indicates that already at birth children show cortical tracking for speech in both familiar and unfamiliar language (Ortiz Barajas et al., 2021). They can even discriminate unfamiliar language-related patterns (Abboub et al., 2016), even though the process of perceptual narrowing has not begun yet. This process takes place between 6 and 12 months of age, during which infants improve their phoneme discrimination abilities for native phoneme contrasts, but worsen/lose this ability for unfamiliar, socially irrelevant phoneme contrasts (Jansson-Verkasalo et al., 2010; Jusczyk et al., 1993; Shafer et al., 2011). In language-specific analyses, backward speech is often used as a control condition for the familiar language, since it matches forward speech in pitch, intensity and duration. Even if backward speech violates several phonological properties of human speech, as some backward sequences cannot be produced by a human speaker (Cowan et al., 1985), it still sounds like an articulated language (Kimura & Folb, 1968). However, further studies have shown that these phonological properties are important to newborns, as they help them discriminate between languages, which they can do in forward speech but not in backward speech (Ramus et al., 2000). As forward speech seems to activate several brain regions stronger than backward speech (Dehaene-Lambertz et al., 2002), we expected that the cortical tracking of the unintelligible, backward language might be worse or similar to that of the natural, familiar language.

### Cortical tracking of prosodic manipulated speech is weaker than that of natural speech

Studies investigating the more detailed underlying factors of speech processing have shown that infants are sensitive to the rhythmical regularities in the auditory input (François et al., 2017), and can segment speech based on statistical information (Teinonen et al., 2009). Stress is a salient prosodic cue for newborns in speech perception (Sansavini et al., 1997). At this age, prosody is a more important cue than word order to indicate linguistic units (Benavides-Varela & Gervain, 2017). This was also shown in a study (Fló, Benjamin, et al., 2022) where sleeping neonates could track transitional probabilities in a speech stream, and that these, together with prosodic cues, allowed them to segment the speech into word-like units. These studies point towards prosody as a strong, important linguistic cue, that newborns are specifically attuned to, and led us to expect that a change in prosody would make the familiar rhyme more difficult to track.

### Prenatal learning

Of further interest in the process of language development is the fetus’ ability to hear and build a lasting memory of an auditory stimulus. Prenatal learning is the process of recognition and discrimination of a familiar stimulus by the newborn, after repeated exposure to that stimulus during fetal life. Prenatal learning has been studied with music, where newborns and infants who had been exposed to a melody during pregnancy had a higher ERP amplitude to the familiar notes of the same melody than the infants that had not been prenatally exposed (Partanen, Kujala, Tervaniemi, et al., 2013) or where 1-month old infants have a heart rate deceleration in response to a piano melody that had been played to them during pregnancy (Granier-Deferre et al., 2011). Already 34-38 gestational week fetuses show heart rate decelerations in response to a familiar rhyme (DeCasper et al., 1994; Krueger & Garvan, 2014), and newborns preferentially react (sucking) to a rhyme presented during pregnancy even if not recited by their mother (DeCasper & Spence, 1986). A heart rate reaction study (Krueger et al., 2004) showed that learning in a fetus is likely affected by both prior experience (amount of recitation of a nursery rhyme) and the timing of that experience in relation to fetal development (maturation of the autonomic nervous system). Another study of prenatal learning has used specific linguistic stimuli (Partanen, Kujala, Näätänen, et al., 2013a), exposing fetuses to a pitch change in the middle of a trisyllabic word (atypical for the native language of the participants). Newborns exposed during pregnancy reacted with a stronger mismatch response (MMR) than the control group to middle-of-the-word pitch changes, while the MMRs to other deviants (vowel identity change, normal in native language) did not differ between groups. This suggests that prenatal exposure to specific linguistic stimuli shapes the way newborns react to and discriminate these language features. Given this literature, we hypothesized that cortical tracking of familiar rhymes would be higher than cortical tracking of unfamiliar rhymes in newborns.

### Prenatal stress factors and predictive value of cortical tracking

As the fetal nervous system slowly develops in the womb, it is subject to many influencing factors, from nutrients to hormones and external input (De Haan & Johnson, 2005). Studies of prenatal maternal depression indicated a negative impact on infant’s cognitive development (Barker et al., 2013; Deave et al., 2008; Field, 2011), which led us to hypothesize that infants of depressed mothers would have lower brain-to-speech coupling than infants of non-depressed mothers. Furthermore, fetal stress has also been independently associated with negative cognitive outcomes (Bergman et al., 2010), and as hair cortisol can be used as a marker of chronic stress (Russell et al., 2012) in a non-invasive way, we examined the correlation between newborns’ hair cortisol and their ability to track the familiar rhyme. Another important outlook of the current study was to assess the predictive value of the cortical tracking abilities of newborns. As other studies reported that infant cortical tracking positively correlated with language development (Menn, Ward, et al., 2022; Ní Choisdealbha et al., 2023), we also measured the infants’ language development at 6 months of age using the Bayley scales (Macha & Petermann, 2015) and investigated the correlation with newborns’ cortical tracking ability.

### Study purpose

In this expanding field of early language development, an increasing number of studies are being made with preverbal infants or even fetuses. While earlier studies relied mainly on behavioral (gaze, suckling, fetal movement) or physiological methods (heart rate), more recent studies started to use brain activity measurements, like EEG (Partanen, Kujala, Näätänen, et al., 2013a) or NIRS (Abboub et al., 2016; Gervain et al., 2008). However, even when the same measurement is used, data is often analyzed in very different ways. Given the high variability used in data analysis, studies are difficult to compare and rarely replicated. Therefore, a side purpose of the present study was to compare some of the most frequently used methods by applying them to the same dataset, the research question being to find evidence of cortical tracking, language discrimination and prenatal learning in 2-week-old infants. Using a fine-grained linguistic paradigm, we additionally aimed to investigate language structures on different levels (rhythm, phonetic cues) and better understand which parts of language are actually shaped before birth. Further, we investigated the influence that prenatal stress factors might have on the newborn’s cortical tracking and how this reflects on later language development.

## Methods

The current report is part of a longitudinal study funded by the Austrian Science Fund (FWF), in which infants and their parents were followed up from the 34^th^ week of gestation to their first birthday. During the pregnancy, the mothers filled out the Edinburgh Postnatal Depression Score (Cox et al., 1987) for screening of depression symptoms (Fig. 1). Infant EEG data was recorded at the 2-week appointment, and developmental diagnostic data was acquired at 6 months with the Bayley Scales for cognition and language (Macha & Petermann, 2015).

**Fig. 1:**
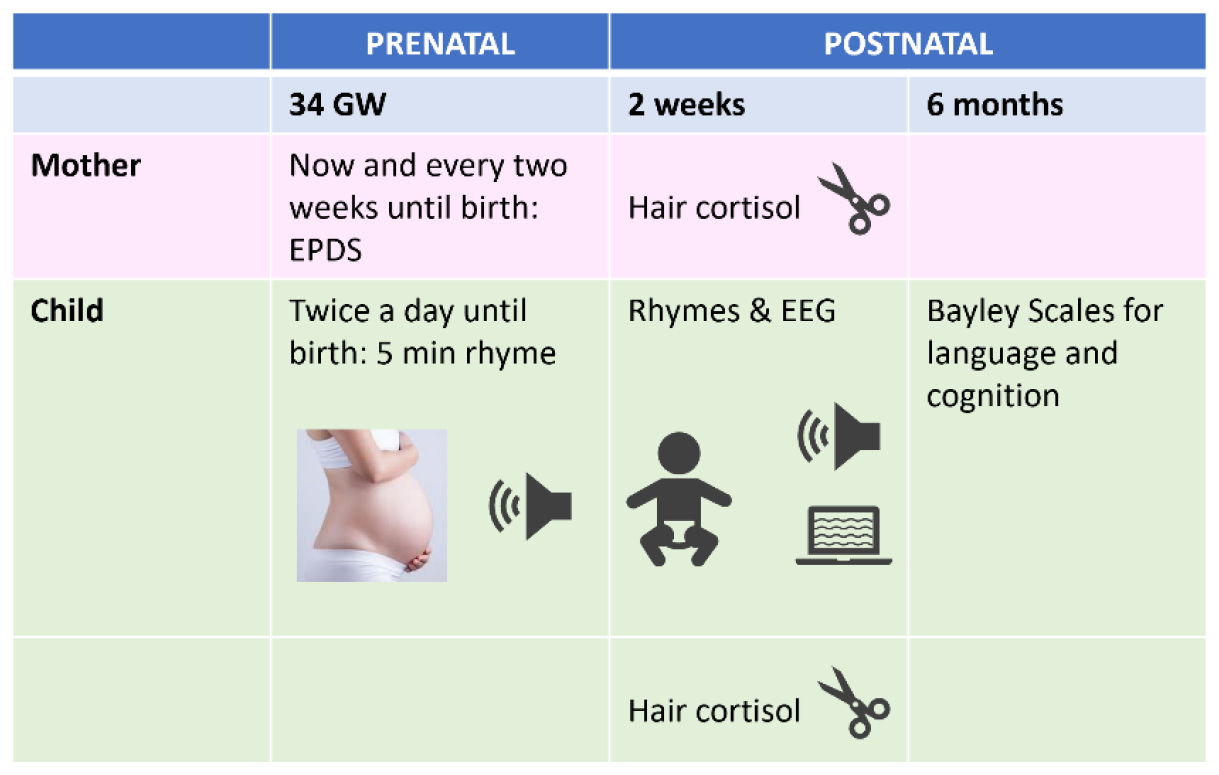
Study design. From the 34^th^ gestational week until birth, the mother played the rhyme twice a day and completed questionnaires every two weeks. At two weeks after birth, infant EEG was recorded and infant and mother hair samples were acquired. At 6 months of age, the developmental diagnostic of cognition and language was performed. The figure photo is just symbolic, as the children were just 2 week old and did not wear headphones, but were stimulated via loudspeakers. EPDS = Edinburgh Postnatal Depression Scale.

### Participants

We recruited 66 participants, and by the time of the analysis 65 had been born. Due to three dropouts before the first recording, only 62 newborns were recorded. Another two participants were excluded because of very noisy data or technical problems during the recording, leaving 60 participants (29 female, 31 male) for the analysis. The mean age at recording was 16.48 days, SD = 3.64 days. The mean gestational age at birth was 278 days (39 weeks and 6 days), SD = 7.9 days. Four participants were born before term (gestational age at birth < 38 weeks), with the smallest gestational age being 36 weeks and 3 days. The mean APGAR score at birth was 9.38, SD = 0.997 (averaged values from 1 min, 5 min, and 10 min). Only two children had an average APGAR lower than 7, and the lowest 10 min APGAR score was 6.

### Experimental design

The pregnant women were requested to play a specific rhyme (audio file) to their abdomen for 5min twice daily from the 34^th^ week of gestation until childbirth, using standardized conditions (sitting or lying down, with 73dB stimulation volume). The women were randomly assigned to one of two groups: Group 1 (N = 32) played Rhyme 1 (*Es tanzt ein Biba Butzeman*) and Group 2 (N = 34) played Rhyme 2 (*Es war eine Mutter*). At two weeks after delivery, the infants were exposed to both rhymes and their EEG activity was measured. One of the rhymes was the familiar one they had been exposed to before birth, while the other one was unfamiliar (the rhyme used for the other group). Additionally, three manipulated versions of the familiar rhyme were played during the EEG recordings: a low-pass filtered version, a rhythm-modified version, and a time-reversed version (backwards speech, also named here the “unintelligible language” condition). The different rhyme manipulations aimed to reveal the language structures present at birth. The (1) low-pass filtering mainly removed phonological cues, so it was named the “phonological paradigm”. It was based on the hypothesis that it would grossly resemble how the fetuses heard the rhyme in the womb, because the abdominal wall, uterine wall, and amniotic liquid would act as a low-pass filter. This filtering also removed part of the phonological elements of the nursery rhyme but would leave most prosodic cues intact. In this way, we could investigate the importance of phonological cues for infants. The (2) rhythm manipulation was based on the hypothesis that infants can follow the rhythmic structure of the stressed and non-stressed syllables in a rhyme, and that changing the rhythm of the familiar rhyme makes it harder to recognize. The (3) backwards speech manipulation was meant to investigate if speech processing is language-specific at birth, and was based on the hypothesis that a newborn can already recognize the maternal language and discriminate it from unintelligible language.

These five rhymes (conditions) were played in random order for about 3min each (an approximately 1min rhyme looped for 3 times), with 2min of silent intervals between them. The beginning of the recording also included a 3min silent baseline and 1.5min of beeps (1000Hz pitch, 100ms length, at 1.5s interstimulus intervals), so that the total recording length was 29.5min. The stimulation volume was 60dB, measured at the infant’s head, from loudspeakers placed at 1m distance, while the mother wore noise-cancelling headphones with a white noise playing, so that she could not tell what rhyme was playing in the room.

### Preparing auditory stimuli

The two rhymes were chosen to have different rhythms. Rhyme 1 (R1), *“(Es) tanzt ein Biba-Butzemann”*, had a 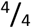 meter with a trochee rhythm, emphasizing beats 1 and 3. Rhyme 2 (R2), *“(Es) war eine Mutter”*, had a ¾ meter with dactyl rhythm, accentuating beat 1 (Figure 2). For Rhyme 1, the average syllable rate was 3.12 Hz (SD = 8.88 Hz), while for Rhyme 2, it was slightly higher at 3.52 Hz (SD = 7.27 Hz). The prosodic stress rate, referring to the interval between stressed syllables, was 1.40 Hz (SD = 5.76 Hz) for Rhyme 1 and 1.07 Hz (SD = 4.74 Hz) for Rhyme 2. The rhymes were recorded in a sound-proof studio as recited (not sung) by a professional male speaker. The rhymes were recorded with the lingWAVES Sound Pressure Level meter microphone and the lingWAVES software, version 3.1 (Copyright 2016 WEVOSYS). Afterwards, the manipulated conditions were obtained by altering different parameters in the software Audacity version 3.0.2.0, and Praat version 6.0.49, as further described. For the backward speech manipulation, the rhymes were inverted and played backward. This approach matches the characteristics of intensity and pitch to the unmanipulated version, while simultaneously creating the impression of an unfamiliar, unintelligible language. For the phonological manipulation, a low pass filter of 1000Hz was applied, with 6dB decrease per octave, removing phonological aspects and primarily retaining prosodic information. In addition, the “WahWah” function was applied to especially accentuate low frequencies, modulated by a moving band-pass filter (LFO = 1,5 Hz, depth = 70%, resonance = 2,5, Wah Frequency Offset = 30%, Output gain -6.0 dB). As a final step, the loudness was adapted to match the unmanipulated version. In the prosodic manipulation, we modified the pitch (± 25%) and the length (± 20%) of the syllables with the aim of creating an arrhythmic accent pattern. For Rhyme 1, in the first bar, the first, second, and fourth beats were heightened, while the third beat was lowered. In the second bar of Rhyme 1, only the third beat was heightened, and the others were lowered. In Rhyme 2, in the first bar, the first and second beats were heightened, and the third beat was lowered. In the second bar, only the first beat was heightened, with the second and third beats lowered. These patterns were consistently applied throughout the rhymes.

**Fig. 2:**
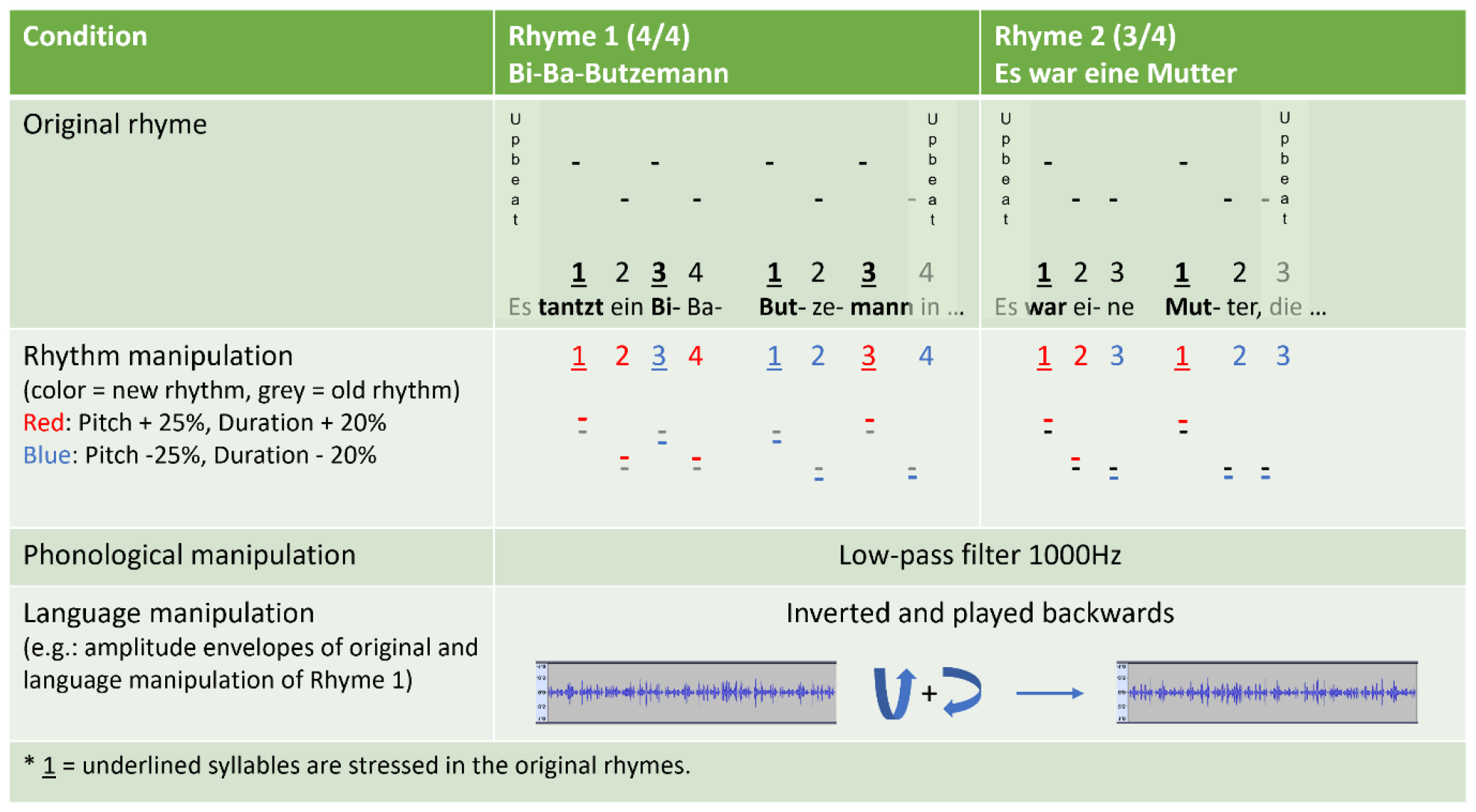
Experimental design of auditory stimulation. In Rhyme 1, the syllables in each measure are numbered from 1 to 4, and in Rhyme 2 they are numbered from 1 to 3. The first syllable (*Es*) is not stressed and is therefore counted as upbeat. The stressed syllables in the original rhymes are underlined. The amplification or reduction of stress of each syllable is marked with red and blue, respectively. For the phonological manipulation, just a low-pass filter was applied. For the language manipulation, the rhyme was amplitude-inverted and time-reversed.

### Processing of the audio data (speech envelopes)

For the neural speech tracking analysis, we extracted the speech envelopes from the audio files with a custom-made Matlab script using the Chimera toolbox (Smith et al., 2002). First, nine cochlear frequency bands at equal intervals were defined (using the function equal_xbm_bands from the Chimera toolbox). For each of these frequency bands, the data was band-pass filtered in the corresponding frequency intervals, the Hilbert transform was applied, and the amplitude (absolute value) was extracted. Then the mean over frequency bands was computed and the data was transformed in percentage (the data was divided by the maximum value). The files were resampled to 500Hz to reduce computation times but preserve a high information level in the data. As previously described, the audio files were then added as another channel to the EEG files.

### Equipment

The EEG data was recorded at a 1000Hz sampling rate, using the EGI system (Electrical Geodesics Inc., US, recently acquired by Magstim, Canada) with hdEEG caps (HydroCel Geodesic Sensor Nets), with 124 channels, and the NetStation acquisition software (version 5.4.2, release r29917). The recordings were performed either in the university lab or (most often) at the parents’ home. The total recording time was 29.5min, and during the recording the babies would usually lie in their parents’ arms, or sometimes on the bed. The rhymes were played via a separate laptop than the recording one, using a custom Matlab (The MathWorks Inc. 2022, MATLAB version: 9.13.0, R2022b, Natick, Massachusetts: The MathWorks Inc. https://www.mathworks.com) script, that implemented the Psychophysics Toolbox extensions (Brainard, 1997; Kleiner, 2007; Pelli, 1997). In previous pilot studies, there was an evident delay in the markers that would accumulate over the 1min loop of the continuous rhyme and would reach significant lags of tens to hundreds of milliseconds from the real audio stimulus. As this was relevant only for long, continuous auditory files, while most literature uses short stimuli, the synchronization problem had not been previously addressed, which led us to a custom-made solution (see Supplementary Material).

### Data preprocessing

The data was preprocessed using Matlab version R2021a, the EEGLab toolbox (Delorme & Makeig, 2004) and the Fieldtrip toolbox (Oostenveld et al., 2011). The raw EEG files were resampled to 500Hz, and the EEG markers were corrected to be synchronized with the audio signal. Next, the data was filtered with the APICE pipeline for preprocessing of continuous data (Fló, Gennari, et al., 2022), which involved multiple steps described in the supplemental material. This resulted in continuous data and a matrix of “artifacts” storing the times, channels and epochs marked as “bad”, i.e., those that could not be interpolated or corrected through filters. The two outermost electrode rows, as well as the occipital electrodes, were excluded because of high artifact prevalence, leaving 82 channels for analysis. The EEG was then segmented into epochs corresponding to the different rhymes, and the speech envelopes were then attached to their corresponding EEG data.

### Computation of classical coherence

Coherence is a method used to compare two electrical signals, often applied in the analysis of brain connectivity between different regions (Nunez et al., 1997; Rosenberg et al., 1989). Coherence is expressed as a function of frequency, and it is computed for each frequency value using the cross spectral density of the two channels compared, normalized by the auto-spectral density (or “power spectrum”) of each channel for that frequency value. The formula for coherence between channels *i* and *j* at frequency *f*, estimated over *N* trials, is (Nunez et al., 1997):

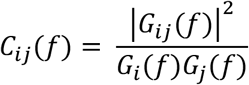

Where *G*_*ij*_(*f*) is the cross-spectral density function for channels *i* and *j* over *N* trials, computed by using the formula (Nunez et al., 1997):

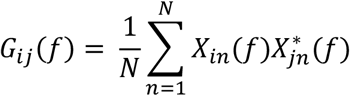

Where *X*_*in*_(*f*) is the complex Fourier transform of each trial (*n*) of channel *i* and 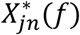 is the complex conjugate (*) of the complex Fourier transform at epoch *n* for channel *j*.

The two channels compared can be a “stimulus” (the speech envelope) and a “response” signal (an EEG channel), and the method has been widely used for the study of brain activation during speech (Bourguignon et al., 2013; Gross et al., 2014; Kolozsvári et al., 2021). For the coherence analysis, the EEG epochs of the current study were segmented in 3s trials, overlapping by 50%, and the trials that had been marked as bad in the APICE pipeline were removed. The 3s length of the trials was chosen to include three cycles of the smallest frequency that we wanted to analyze, that being the 1 Hz frequency. Datasets with less than 25 trials (per participant and condition) were excluded, so that some participants resulted in having data only for certain conditions.

Using filters optimized with the Parks McClellan algorithm, the data was filtered in two frequency bands, one at the syllable rate (the frequency at which the syllables alternate, 2.62-3.67Hz in Rhyme 1 and 3.00-4.02Hz in Rhyme 2), and one at the stress (or prosody) rate (the frequency at which the stressed syllables alternate, 1-2Hz in Rhyme 1 and 1-1.67Hz in Rhyme 2). Coherence was then computed for each of these bands based on the cross-spectral density using the Fieldtrip toolbox (Rosenberg et al., 1989), which resulted in a series of 3 to 4 coherence values, 0.33Hz apart, for each frequency point in the given interval. The coherence values were then averaged over trials and channels. For further statistical analysis, three coherence values were used: the average coherence over the frequency band of interest, the maximum coherence in that band, and the average of the two maximal coherence values in that frequency band. All three methods were compared with the aim of avoiding frequency smoothing but also to keep as much information as possible from the entire frequency band.

### Computation of Hilbert coherence

The Hilbert coherence is another measure of correlation between two signals but in contrast to the classical coherence, it is computed as a function of time. In the Hilbert coherence, each sample (value at each timepoint) is weighted by its amplitude, while for the classical coherence, which is a function of frequency, not of time, the samples at a given frequency are weighted by their joint amplitude together (Bruña et al., 2018). In this way, while the classical coherence estimates the synchronization between tow signals for just one frequency value, the Hilbert coherence estimates the synchronization over the whole frequency band in which the data has been filtered. In our analysis, this means that, for example in the prosodic interval (1-2 Hz), the classical coherence was computed at fixed frequency values (as reported above: 1Hz, 1.33Hz, 1.67Hz and 2Hz). This leads to a frequency smoothing, as coupled frequencies are averaged together with uncoupled ones over each of the 0.33Hz bandwidths. Further smoothing occurs when the average coherence over the whole prosodic or syllable frequency band is computed. This might lead to an underestimation of coupling. The Hilbert coherence, on the other hand, uses not the Fourier or wavelet phases, but the temporal, Hilbert phase of the signal. Therefore, at each timepoint it calculates the instantaneous phase over the whole band, and not from an average of frequencies (Bruña et al., 2018). Thus, Hilbert coherence might identify a brain synchronization with an oscillator that changes frequencies (e.g., the prosodic and syllable frequencies in our rhymes were also not perfectly constant). This might reveal synchronizations that are otherwise underestimated by the classical coherence. For the Hilbert coherence, data was segmented, cleaned, and filtered in two frequency bands just like for the classical coherence. The data was then Hilbert-transformed, and the Hilbert coherence was computed for each trial and each channel, based on the formula (Bruña et al., 2018):

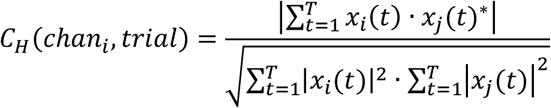

Where *C*_*H*_(*chan*_*i*_, *trial*) is the Hilbert coherence computed for each channel and each trial, between the channel *i* and the audio channel *j, t* is the timepoint, with T being the last (highest) timepoint of each trial, *x* is the band-pass filtered and Hilbert-transformed data, * is the conjugate of a complex number, and | | is the absolute value of a complex number. The Matlab code for the Hilbert coherence is described in the supplemental material. The resulting coherence values (one for each frequency band) were then averaged over trials and channels.

### Computation of surrogate data for coherence significance

To check the statistical significance of the coherence values, they were compared to a distribution of 1000 “surrogate” coherences computed between the EEG data and 1000 circular shifts (Keitel et al., 2017) of audio data from a different condition. The percentage of surrogate coherence values higher than the measured coherence gave the *p* value, i.e., the probability that the measured coherence is significant.

### Statistical analysis for the classic and Hilbert coherence

For the statistical analysis, the average classical coherence over the frequency points in each frequency range (prosodic or syllable) was used. Additionally, considering that perhaps each child was following best at a different frequency, the statistical analysis was performed on (1) just the maximum value and (2) the average of the highest two values.

Both the classical and the Hilbert coherence analyses were performed first on the whole dataset and second on a selection of values that only included the significant coherences based on the surrogate distribution. The results of the analysis of only the significant coherences are presented in the Supplemental material.

### Computation of multivariate temporal response functions (mTRF)

The mTRF toolbox (Crosse et al., 2016) was used to compute multivariate temporal response functions (mTRF). mTRF is an encoding model that describes the association between an auditory stimulus and a brain signal (Crosse et al., 2016). In brief, a TRF is a time-lagged regression model that describes a neural activity as a time lagged response to a stimulus (e.g., the amplitude in the speech envelope), where the interpretation of a TRF follows that of an ERP in response to a continuous stimulus (Lalor et al., 2006). In the case of newborns, where ERP components are stretched over a longer period of time due to brain immaturity, we chose time lags up to 1000ms after the stimulus for our encoding model. The resulting value of the TRF at a certain time lag (e.g., 450ms) indicates “how a unit change in the amplitude of the speech envelope would affect the EEG” 450ms later (Crosse et al., 2016). To account for the multiple frequency bands contained in the speech stimulus, the multivariate analysis computes the TRF for each frequency band defined (similarly to Attaheri et al., 2022). However, in our case, the bands of interest being so narrow, we computed the TRF on the filtered data and speech for three different frequency intervals of interest: on data band-pass filtered for 0.5-2Hz, covering the prosodic stress rate, for 0.5-4Hz, also covering the syllable rate, and for 0.5-10Hz, which additionally covered shorter linguistic structures, such as phonemes. That means that we computed de facto a univariate TRF for each of these intervals.

The continuous preprocessed data was down-sampled to 100Hz to increase the computation speed. As each 1min long rhyme was played three times in a loop, we obtained six segments (folds) of 30s each. Only the folds with less than 30% bad times or bad channels were kept. The last fold was used as test data, and the others as training data. The data was z-scored and split into a stimulus channel (i.e., the speech envelope or the markers of the stressed syllables) and the response channels (the EEG channels). For each subject and model, an optimal regularization parameter (lambda) was chosen via cross-validation from the possible values of 10^-3^ to 10^10^. A forward (encoding) regression model was then trained on windows of 1200ms length (-100 to 1100ms) to predict the EEG signal from the speech envelope and used to predict neural activity in the test set. Afterwards, the Spearman correlation coefficient *r* between the predicted and the real EEG signal was computed.

### Statistical analysis of mTRF models

The mTRF models were analyzed in two ways, (1) using the correlation strength (*r*) between the predicted and measured signal (on the test set), and (2) by applying a cluster-based permutation test on the weights of the models from each condition and participant. For the first analysis, Wilcoxon signed rank tests were performed in RStudio, while for the second analysis we used Matlab and the Fieldtrip toolbox.

In addition to the correlation coefficient, the statistical analysis was performed also on *r*^2^ and |*r*| (absolute value), as the negative correlations might reflect that the model was in counter-phase, but not ill-fitting. The cluster-based analysis of the model functions used a *t*-test for repeated measures as a statistical test between conditions.

### Computation and analysis of mutual information

The mutual information (MI) computes how much of the entropy in one signal (response = EEG) can be reduced by the information contained in the other signal (stimulus = speech envelope), without assuming a linear relationship. In the current study, we computed it between the same frequency bands in the speech envelope and the brain signal, but analyses of combinations between different frequency bands in the “stimulus” and the “response” can also be done (Cogan & Poeppel, 2011). The advantage of MI analysis in the current study is that it compares not just the amplitudes of the signals, but also the phases, in all possible combinations. For example, if the stimulus amplitude changes the phase (but not the amplitude) of the response, then that would be caught by the MI of phase-amp (EEG phase vs. speech envelope amplitude). Sometimes, auditory stimuli lead to such phase resets in the brain signal (Canavier, 2015). Inversely, groups of neurons might fire simultaneously at a phase change in speech, leading to a larger amplitude in the EEG, in which case MI for amp-phase would increase. These changes are not easily uncovered by the other methods (coherence, TRF), which is why the MI might reveal effects that are otherwise masked.

For the computation of mutual information (MI), the continuous preprocessed data was segmented into 3s segments, and the noisy segments were removed. The data was band-pass filtered with an optimized algorithm in 1-Hz-wide frequency bands between 0.5-10Hz, to cover the same interval as the mTRF. The phase and amplitude were extracted with a Hilbert transform and the MI was computed between all combinations of phase and amplitude between EEG and the speech envelope. Considering the immaturity of the newborn brain, interindividual variations in response lag had to be accounted for, which is why we computed MI with possible lags from 0 to 800ms between the two signals, choosing the highest MI from all the lags. The MI was then averaged over electrodes and trials. The four resulting MI tables (phase-phase, phase-amp, amp-amp, amp-phase) were then analyzed in RStudio. The average MI over each frequency interval (similar to mTRF models: 0.5-2Hz, 0.5-4Hz, and 0.5-10Hz) was compared between conditions using Wilcoxon signed rank tests.

### Maternal prenatal distress, fetal stress, and infant language development

The maternal prenatal depression score was evaluated with the Edinburgh Prenatal Depression Scale (EPDS) (Cox et al., 1987) at 34 weeks of gestation and then every 2 weeks until birth, via an online survey. The average score across the pregnancy was computed and used for further correlation analyses. Hair samples from the mother and child were acquired at 2 weeks after birth from the occipital area and used for the analysis of hair cortisol, to measure chronic fetal stress and chronic maternal prenatal stress. The infant language development was assessed by trained testers at 6 months of age using the Bayley Scales (3^rd^ edition) for expressive and receptive language development (Macha & Petermann, 2015). The correlations were computed in RStudio using Spearman’s correlation.

### Statistical analysis

Due to assumptions violation for normal distribution and/or homogeneity of variance, nonparametric tests were performed, namely Wilcoxon signed rank tests for dependent samples comparisons (comparisons between conditions), Wilcoxon rank sum test for independent samples (comparisons between groups) and Spearman’s correlation for the relationship between neural speech tracking and prenatal factors or language development. The statistical analyses were made with RStudio, version 2022.02.3 (RStudio Team, 2020). Where there were outliers that strongly distorted the data, they would be cut off at 3 SD over the average. The minimum sample size for performing the Wilcoxon signed rank test was N = 6 (total sample size), with 2 groups of size within groups N = 3 (Dwivedi et al., 2017).

## Results

In the following, we present the main results for the (1) prenatal learning, (2) language manipulation, (3) rhythm manipulation, and (4) phonological manipulation across methods. Further we present the (5) comparison of measured coherence to surrogate coherence and lastly the (6) correlation of neural tracking with prenatal factors and later language development. Details on the agreement between methods, as well as statistics of non-significant results and figures of trending effects can be found in Supplemental materials.

### 1. Prenatal learning

In the analysis of classical coherence and of mutual information we found a familiarity effect, with higher coherence/MI for the unfamiliar rhyme than for the familiar one (Figure 3). In the Hilbert coherence and mTRF analyses there was no effect of familiarity. In the classical coherence analysis, the effect was significant (*V* = 432, *p* = 0.048) at the prosodic rate for the average coherence and trending (*V* = 446, *p* = 0.065) at the syllable rate for the average of two maximums. In the MI analysis, the effect was significant in the amplitude-amplitude comparison over all frequency intervals (0.5-2Hz: *V* = 1112, *p* = 0.004; 0.5-4Hz: *V* = 1046, *p* = 0.021; 0.5-10Hz: *V* = 1059, *p* = 0.016), and trending (*V* = 943, *p* = 0.085) in the phase-phase comparison for the 0.5-10Hz interval.

**Fig. 3:**
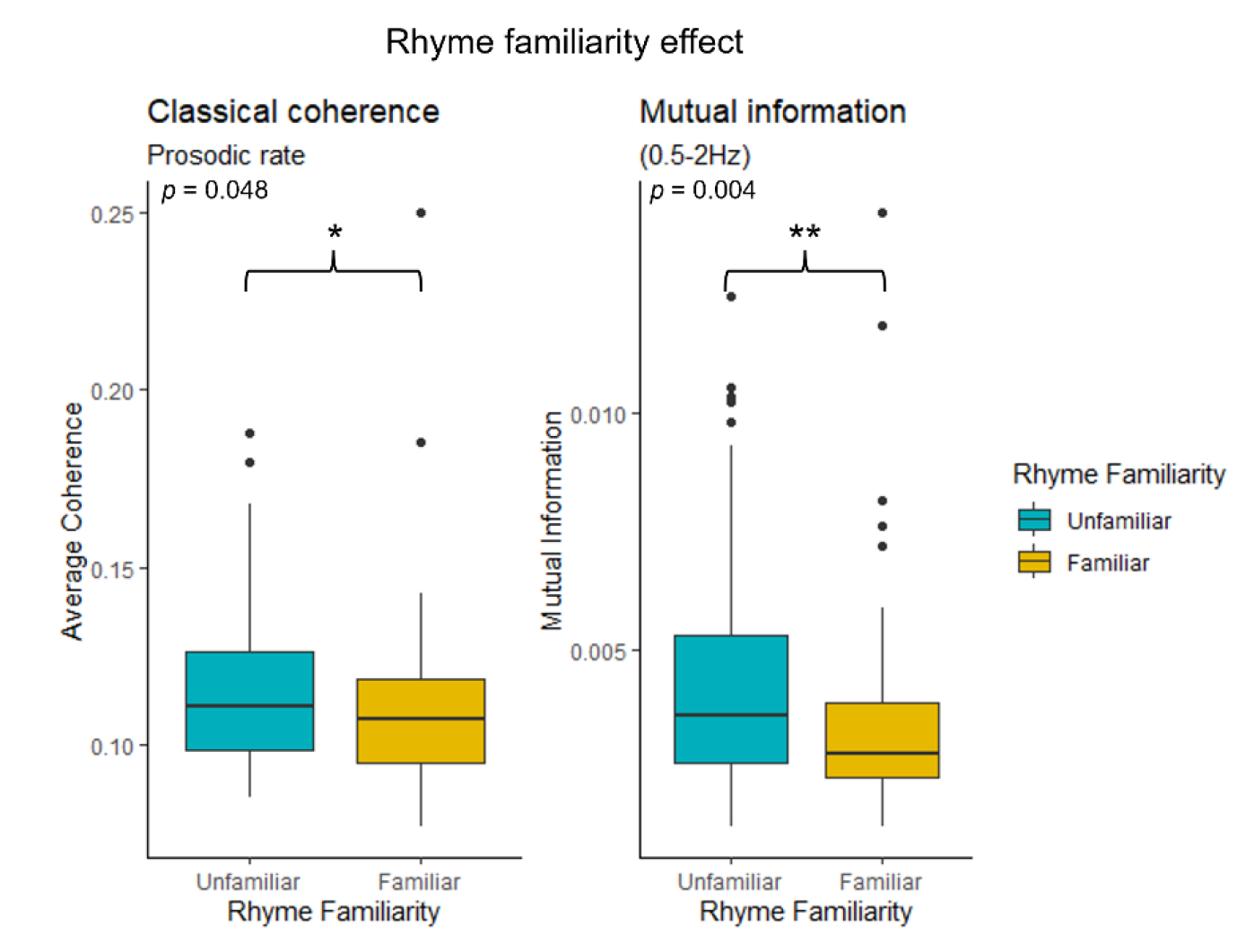
Rhyme familiarity. The rhyme familiarity effects in the Classical coherence (N = 50) and in the Mutual information (N = 55). Note the higher coherence/MI for the unfamiliar rhyme (blue) compared to the familiar rhyme (yellow). * *p* < 0.05, ** *p* < 0.01.

### 2. Unintelligible language discrimination

Across multiple methods we found a significant (*p* < 0.05) or at least trending (*p* < 0.1) language effect, predominantly in favor of the familiar language.

The classical coherence analysis showed a language trend in the prosodic rate when the maximal coherence (*V* = 826, *p* = 0.07) or the average of the two highest coherences (*V* = 813, *p* = 0.09) was used as a variable, but not when the average coherence was used. The trend was for the familiar rhyme to have higher coherence than the language-manipulated unintelligible rhyme. In the Hilbert coherence analysis, there was a significant language effect for the prosodic rate (*V* = 409, *p* = 0.04), but rather in favor of the unintelligible rhyme (Figure 4A).

**Fig. 4:**
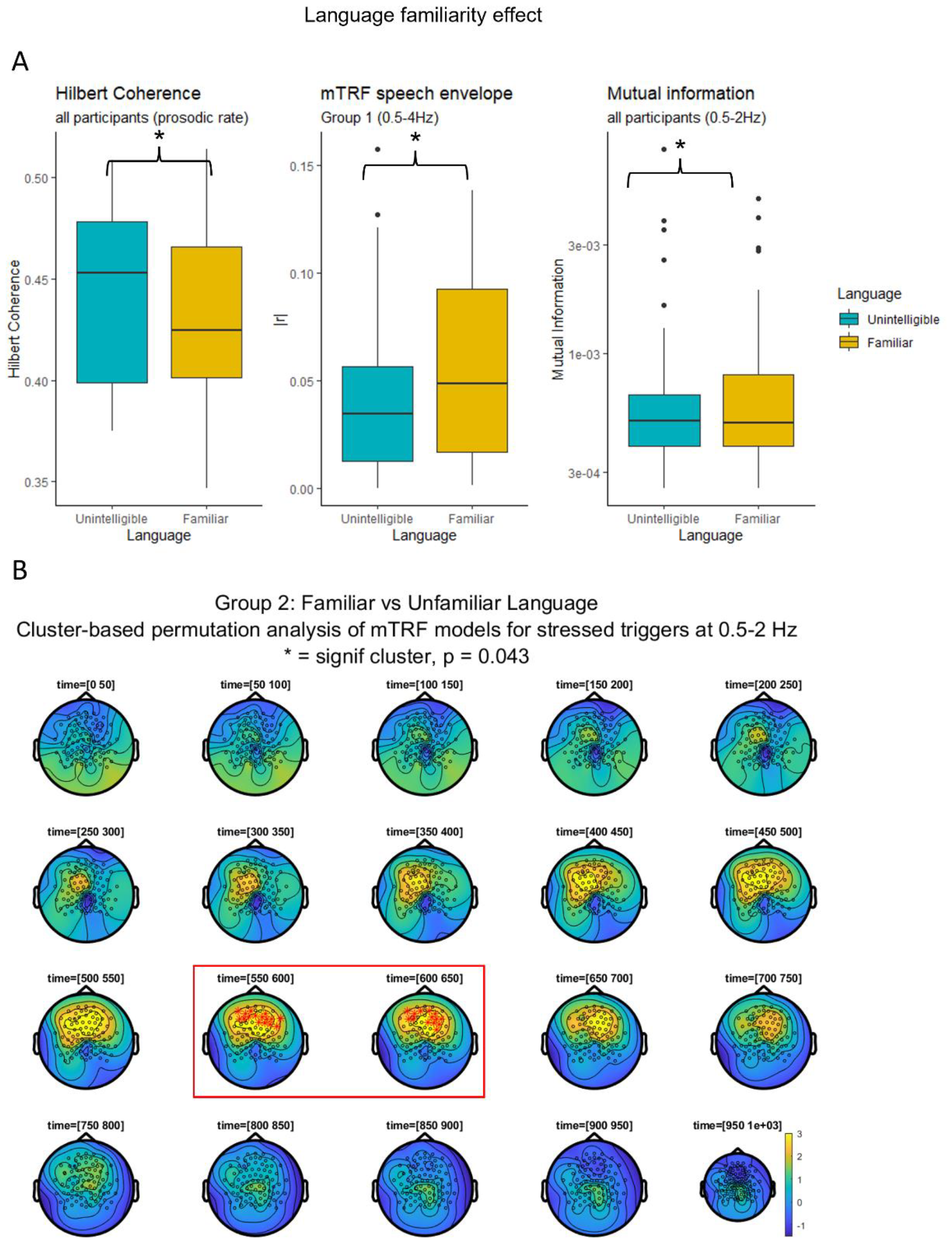
Unintelligible language discrimination. Panel A: The effect of language familiarity in Hilbert coherence, mTRF for the speech envelope, and Mutual Information. According to the TRF for speech envelope, the newborn showed higher coupling with the familiar language than the unfamiliar one. The MI indicates that the phase of the speech envelope of the familiar language entrained the EEG amplitude more for the familiar language than the unintelligible one. The Hilbert coherence depicts a reversed effect, possibly due to a variable oscillator and an attention-orienting effect induced by the unintelligible language. Panel B: Cluster-based permutation analysis of the TRF models for stressed syllables in Group 2. Note the positive cluster (higher predicted signal for the familiar language) between 550-650ms, marked with a red rectangle.

In the mTRF analysis of the speech envelope, there was also a trend for a language effect over all frequency intervals, but only in Group 1, with higher *r*^*2*^ and |*r*| values for the familiar rhyme than for the language manipulated rhyme, p < 0.1. The effect reached significance for |*r*| at 0.5-4Hz, *V* = 158, *p* = 0.048 (Figure 4A).

The cluster-based permutation analysis of the TRF for the speech envelope showed a trending cluster at 750-900ms at 0.5-4Hz, indicating higher activation for the language-manipulated rhyme, *p* = 0.095, but it did not reach significance (neither at other frequencies, nor when analyzed in each Group separately) (see Supplemental Material, Figure S10). However, in the TRF for the stressed syllable triggers, there was a significant cluster in Group 2, bilateral frontally, at 500-650ms, at 0.5-2Hz (*p* = 0.043) and 0.5-10Hz (*p* = 0.045) and trending at 0.5-4Hz (*p* = 0.081) (Figure 4B). This positive cluster indicated higher predicted signal for the familiar rhyme than for the language-manipulated rhyme. The later, trending cluster in favor of the unfamiliar language might suggest that the unintelligible language is tracked with a longer time lag and more spread over time and location, so that statistical significance for a cluster is not reached.

The mutual information analysis showed a language effect only in the amplitude-phase comparison, with higher mutual information for the familiar rhyme, significant for 0.5-2Hz (*V* = 1132, *p* = 0.033) (Figure 4A) and trending for 0.5-4Hz (*V* = 1103, *p* = 0.056).

Interestingly, the TRF correlation coefficient differed between language familiarity in the 0.5-4 Hz frequency band but not in the 0.5-2 Hz frequency band (*V* = 156, *p* = 0.058). This indicates that tracking of information in the frequency band related to syllabic information was more pronounced in the familiar compared to the unintelligible language. This added information in the larger bandwidth extended the difference in cortical tracking of the two conditions to reach statistical significance. A detailed interpretation of the opposing results of Hilbert coherence and TRF/MI is given in the *Discussion* section.

### 3. The rhythm effect

In two methods (classical coherence and mTRF analysis) there was a significant or trending difference between the original rhyme and the rhythm manipulated rhyme, with higher coherence and, respectively, higher mTRF correlation coefficient in the original rhyme (Figure 5). In the other methods there was no effect. The effect for the classical coherence was significant at the prosodic rate for the maximal (*V* = 907, *p* = 0.022) or average of the highest two (*V* = 881, *p* = 0.041) coherence values in the prosodic rate. The effect for the mTRF correlation values was trending in the mTRF for the speech envelope, for |*r*| at 0.5-2Hz (*V* = 126, *p* = 0.081) for Group 1 only, and in the mTRF for the stressed syllables for *r*^*2*^ at all frequency intervals, for Group 2 only (0.5-2Hz: *V* = 126, *p* = 0.081, 0.5-4Hz: *V* = 128, *p* = 0.067, 0.5-10Hz: *V* = 127, *p* = 0.074).

**Fig. 5:**
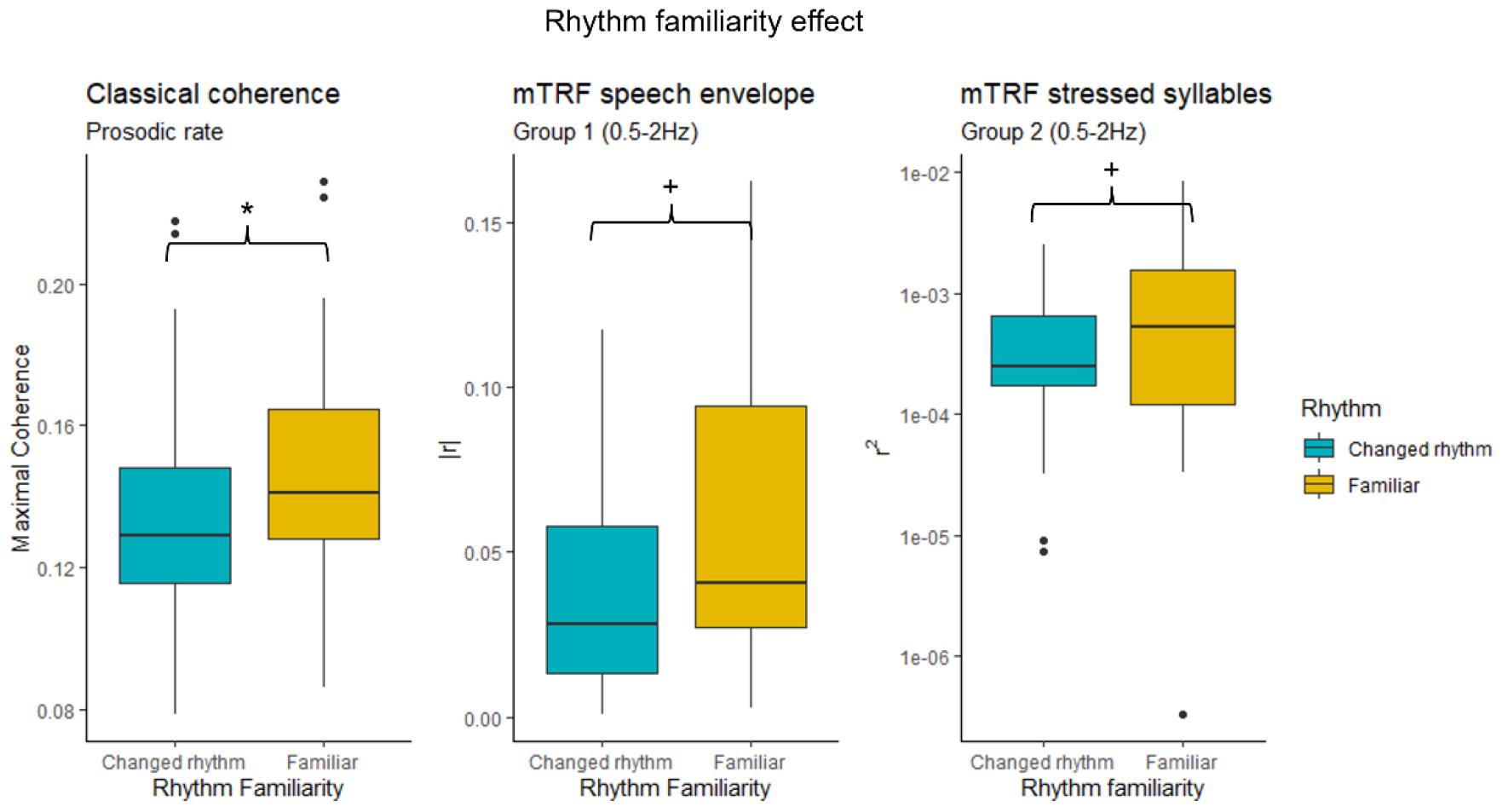
The rhythm effect. Note the higher coherence/correlation coefficients of mTRF models for the familiar rhyme rhythm (yellow) than for the rhythm-manipulated rhyme (blue). Significance notation: + = *p* < 0.1, * = *p* < 0.05

### 4. The phonological effect

Across two methods – the Hilbert coherence and the mTRF analysis – there was a significant or trending difference between the low-pass filtered manipulation and the original rhyme, always in favor of the original version. In the Hilbert coherence analysis, there was a trend (*V* = 799, *p* = 0.064) for a phonological effect in the prosodic rate. In the mTRF for the speech envelope, there was a trend for *r*^*2*^ at 0.5-4Hz in Group 1 (*V* = 101, *p* = 0.093) and in the mTRF for the stressed syllables there was an effect for *r* in Group 1 at 0.5-4Hz (*V* = 107, *p* = 0.044) (Figure 6), which became a trend at 0.5-2Hz (*V* = 106, *p* = 0.051) and at 0.5-10Hz (*V* = 104, *p* = 0.065).

**Fig. 6:**
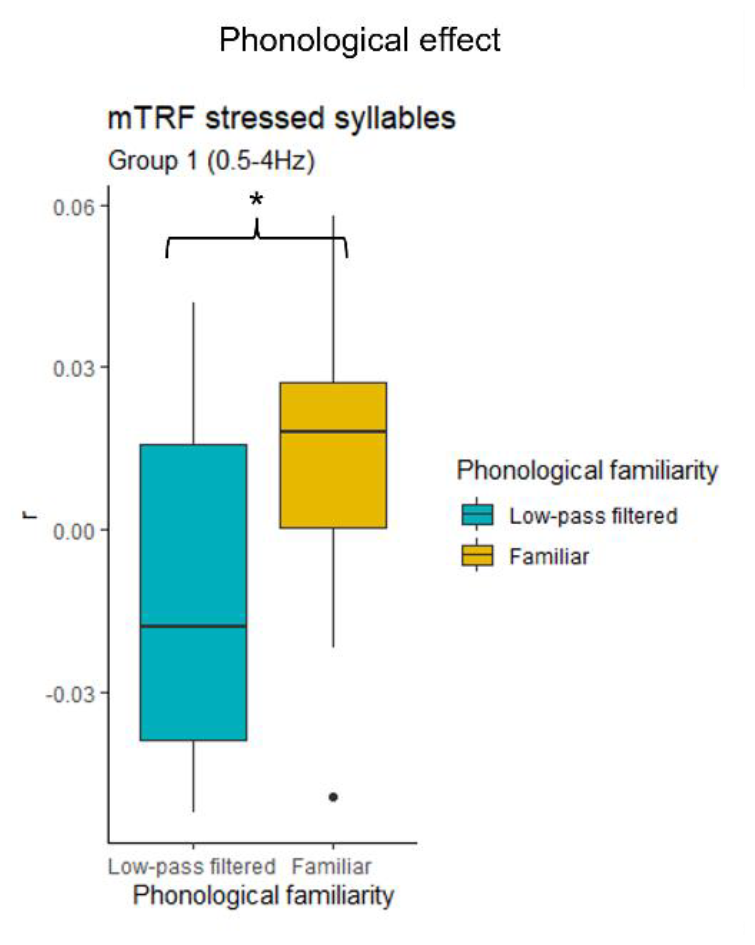
The phonological effect. Note the phonological effect in favor of the familiar unmanipulated rhyme (yellow). Significance notation: * = *p* < 0.05.

**Fig. 7:**
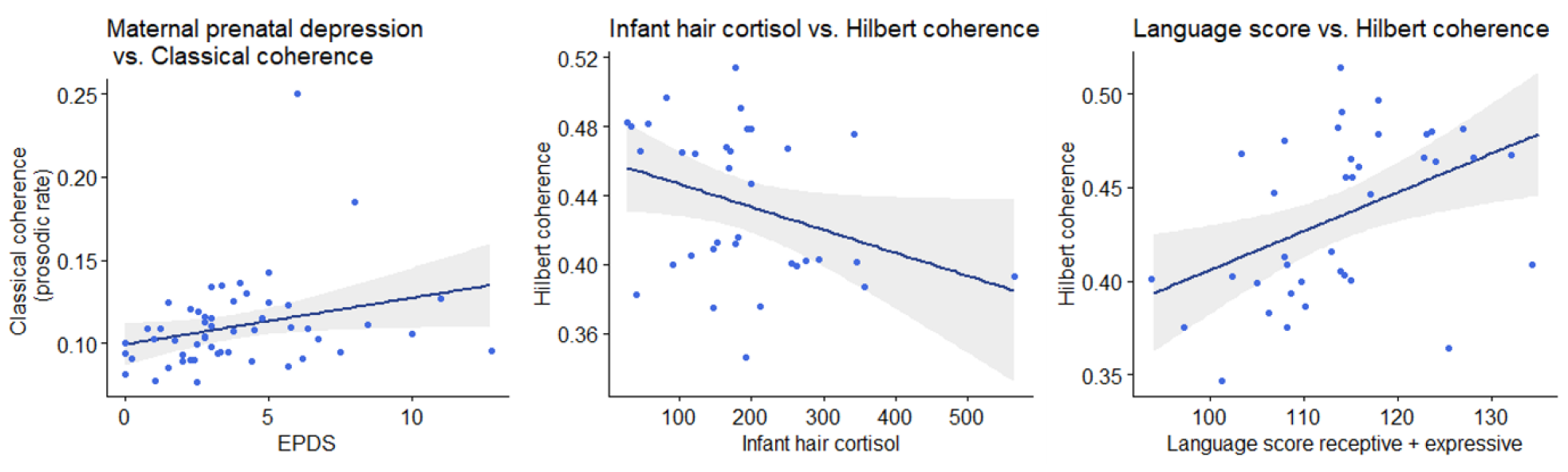
The correlations of cortical tracking with depression, stress, and language development. Note the strong positive correlation between the Hilbert coherence at two weeks and the language development at 6 months (right panel).

### 5. Comparison with surrogate and effects of group and rhyme

Compared to the coherence distribution on surrogate data, in the classical coherence 33% of the participant*condition files had significant coherence values for the prosodic and 38% for the syllable rates; in the Hilbert coherence, 58% had significant coherence values for the prosodic rate and 21% for the syllable rate. For a separate analysis of these files only, see Supplemental Material.

To control for group or rhyme differences, analyses comparing the two groups and the two rhymes were performed. Across all methods there were no group effects, indicating a well distributed sample across the two groups. In some methods there was a rhyme effect, but it alternated between favoring rhyme 1 and rhyme 2, which suggested no consistent bias (see Supplemental Material for detailed results).

### 6. Correlation of cortical tracking with maternal stress and depression and the predictive value for language development

There was a moderate positive correlation between the maternal depression score (EPDS) and the classical coherence for the prosodic rate, with ρ *(rho)* = 0.377, *p* = 0.005, indicating more tracking for the familiar rhyme in children of mothers who were more depressed during pregnancy. Furthermore, there was a negative correlation between the child hair cortisol levels at 2 weeks and the Hilbert coherence at the prosodic rate, indicating less tracking for the familiar rhyme in children with high hair cortisol, ρ = -0.35 (moderate correlation) and *p* = 0.045. The maternal hair cortisol did not correlate with the infant’s neural tracking (ρ = -0.006, *p* = 0.97) or with maternal EPDS (ρ = -0.084, *p* = 0.64). Further, there was a significant positive correlation between the Hilbert coherence for the prosodic rate for the familiar rhyme at 2 weeks and the language composite score (receptive and expressive) from the Bayley scales at 6 months, ρ = 0.50 (large correlation), *p* = 0.001, indicating that the ability to track the familiar rhyme was strongly correlated with later language development. There was no significant correlation between coherence and the Bayley cognitive scale (ρ = 0.21, *p* = 0.20).

## Discussion

Overall, the results indicate that the newborns tracked the unfamiliar rhyme more strongly than the familiar one, which likely is associated with enhanced attention to an unexpected stimulus input. Conversely, different manipulations of the rhymes (backward speech, changed rhythm, and low-pass filtered) were tracked less strongly than the familiar, original rhymes. This may be due to a degradation of different aspects of the signal in the manipulated versions, making it more difficult to track.

### Prenatal learning

There was a clear discrimination of the familiar and unfamiliar rhymes. The amplitude-amplitude MI and the classical coherence were both higher for the unfamiliar rhyme than for the familiar one, which shows an amplitude-driven tracking of the rhymes which is stronger (better correlation between amplitudes) for the unfamiliar rhyme than for the familiar one. The coherence is a function of frequency which takes into account the amplitudes of the signals, while the MI amp-amp is also a function of both signals’ amplitudes, which explains why the two measures are congruent in their effect. There are previous studies that show enhanced brain activity for unfamiliar or more complex stimuli (Henson et al., 2000), or for unfamiliar pitch contours (Homae et al., 2007) which could rely on an attention-orienting effect, ultimately leading to a better neural tracking of the acoustic stimulus. This explains the better tracking of the unfamiliar rhyme in our sample.

The current study is one of the first to show, with direct brain data, that a newborn already has memory traces of a rhyme they heard during the intrauterine life. Evidence of prenatal programing, with changes in speech processing occurring due to stimulation during fetal life, has been previously presented by Partanen (Partanen, Kujala, Näätänen, et al., 2013b). The prenatal learning paradigm in our study, however, is different since it compares two rhymes, both in the same (familiar) language. The result is, moreover, not rhyme-specific, since irrespective of which rhyme was familiar, the unfamiliar one was better tracked.

### Language paradigm

In the language-paradigm, the familiar language had better tracking (i.e. higher mutual information and higher mTRF correlation coefficients) than the unintelligible language. However, the analysis of the Hilbert coherence showed an opposite effect, with higher coherence for the unintelligible language. The analysis of the classical coherence showed no effect. Backwards language should have the same rhythm as the original rhyme, but if we consider that rhythm is syllable based, and that the syllables in the backward speech were phonetically very different, then also the infant’s interpretation of the rhythm would be different. Hence, we propose that the higher Hilbert coherence was due to a different interpretation of the rhythm by the infant, for whom the syllables of the backwards speech were phonetically unfamiliar. Therefore, the stressed syllables were perceived at variable intervals, which might have led to a variable frequency of cortical tracking, that was better measured with the Hilbert coherence. The presence of unexpected, unfamiliar phonemes might have stimulated the infant’s response to be stronger than for the familiar language, which could explain the higher Hilbert coherence for the unintelligible language compared to the familiar one.

However, as the forward linear model (TRF) predicted the EEG better in the familiar language (correlated better with the real EEG signal) than it could predict the EEG for the backward language, this suggests that the predictability of neural tracking was better in the familiar than in the unfamiliar, unintelligible language. Similarly, the phase of the speech envelope of the familiar language entrained the amplitude in the EEG (amp-phase MI) more than the phase in the backward language did, perhaps since it was easier to predict. The predictability of the familiar language phase might have provoked more neurons to fire simultaneously, giving rise to higher cortical EEG amplitudes that correlated with the phase changes in the speech envelope.

The fact that the TRF effect was present only in Group 1 suggests that in Group 2 the level of rhythm familiarity was not different enough between the familiar language and the backwards rendition of Rhyme 2. This might be due to the fact that Rhyme 2, in its original version, already has a rhythm rather unusual for the German language (dactyl, in a ¾ tact). Therefore, neither TRF (linear prediction) nor MI (phase-driven by the speech envelope) were sufficiently different between the two conditions of Rhyme 2.

The existing literature also reports seemingly different reactions to an unfamiliar language. On the one hand, infants orient faster to their native language (Dehaene-Lambertz & Houston, 1998; Kisilevsky et al., 2009) than to an unfamiliar one. Furthermore, in a NIRS study on newborns, May et al. (May et al., 2011) found increased Hb oxygenation to the familiar language and decrease to the unfamiliar language. On the other hand, a study by (Moon et al., 2013) showed that neonates increase the suckling frequency to a nonfamiliar language, which might indicate a general increase in alertness and, by extension, an intensification of cognitive processes. These findings suggest that the response to an unfamiliar language is complex and may result as a combination of increased attention, decreased capacity of prediction, and a more diffuse (i.e., spread over time, location, and oscillations frequency) processing of the unfamiliar phonemes and sentence structure.

Overall, however, our results confirm the newborns’ ability to discriminate the familiar language from an unfamiliar one, as reported by previous studies.

### Rhythm and phonological paradigms

With the rhythm and phonological paradigm, the current study found that newborns used both prosodic and phonologic cues to track the rhymes. In the manipulated rhymes, where these cues had been changed, the classical/Hilbert coherence values and the correlation coefficients of TRFs were lower than for the original rhyme.

The rhythm paradigm effect was always in favor of the familiar rhythm, with the classical coherence and the TRF correlation coefficients being higher for the familiar rhythm than for the manipulated one. It makes sense that the Hilbert coherence in this case was not very different between the two conditions, because the unfamiliar rhythm might have been followed at various frequencies, so that the Hilbert coherence was just as high in the rhythm-manipulated rhyme as it was in the familiar-rhythm rhyme. But the difference in the classical coherence and TRF correlation coefficient suggest that there is a linear relationship at a more constant frequency value between the EEG signal and the speech envelope. This relationship is stronger for the familiar rhythm than for the unfamiliar rhythm. This finding underlines the early sensibility of infants to prosody and expands present literature with evidence from the very young newborns, since previous reports are from older children: 9-month-olds (Martinez-Alvarez et al., 2023), 6-month-olds (Holzgrefe-Lang et al., 2018; Seidl, 2007), or 4-month-olds (Seidl & Cristià, 2008). Additionally, by comparing versions of the familiar rhyme, this result is also an indication that prenatal learning relies quite strongly on rhythmic cues. The low pass-filtered rhyme, missing the phonological cues, also had lower tracking than the original rhyme: lower TRF correlation coefficient and trending lower Hilbert coherence. This effect was significant only in Group 1, but it was also trending in the other group. The reason why the effect would appear only in Group 1 might have to do with the manipulations and rhyme characteristics. In Rhyme 1, the rhythmic cues alone, in the presence of degraded phonological cues, might have not been enough to retrieve the memory of the rhyme, so the low-pass filtered version was harder to follow than the original rhyme. However, in Rhyme 2, the removal of the phonological cues by low-pass filtering did not seem to have this effect, and the prosodic cues (the rhythm and melody of the familiar rhyme) were enough to retrieve the memory of the rhyme. This might be due to the ¾ tact of Rhyme 2, which is less usual for the German language, and might have made the rhyme easier to remember based only on prosodic cues. Overall, the phonological effects suggest that newborns already use phonological cues to track a rhyme. Indeed, a study using resynthesized speech (Ramus, 2002) found that newborns can discriminate between two languages based on rhythm but only if some phonotactic information is present. Our results support this theory of early phonotactic sensitivity, which builds up on earlier studies that found only 4.5 -month-olds and older infants to be able to process phonotactic cues (Friederici & Wessels, 1993; Jusczyk et al., 1994; Mattys et al., 1999).

### Surrogate data comparison

Additionally, the comparison of the coherence values with the surrogate data revealed that over a third of the participant*condition files had significant coherence values for the prosodic and syllable rates; in the Hilbert coherence, 58% had significant coherence values for the prosodic rate and 21% for the syllable rate. It seems the slower frequencies were easier to follow, which is in line with present literature, since it’s at these slower frequencies that one can find the most salient prosodic features of speech. At slower frequencies (i.e., 0.5-2Hz) is where the tact of the rhyme becomes apparent, i.e., the stress of the syllable. The prosodic features that give the stress (pitch, tempo, amplitude, and rhythm) help young infants orient towards and maintain attention for the speech stream (Martinez-Alvarez et al., 2023), and it has been previously shown that young infants have stronger cortical tracking for these low frequencies (delta, 0.5-4Hz) compared to theta or alpha (Attaheri et al., 2022).

### Correlations of cortical tracking with maternal distress and later infant language development

Higher maternal depression scores during pregnancy correlated with more tracking of the familiar rhyme, which seems counterintuitive considering that prenatal maternal depression is considered to negatively impact infant’s cognitive outcome (Barker et al., 2013; Deave et al., 2008; Field, 2011). However, the depression scores in our sample were strongly skewed to the right, with most mothers having low depression scores and just 6% of them having scores above 9, indicating symptoms of depression. This suggests that the slightly higher depression score might also indicate a moderate level of maternal distress, which has been found to associate with better infant cognitive outcome in healthy populations (such as our sample) (DiPietro et al., 2006). Other studies found non-toxic levels of maternal hair cortisol to positively correlate with the infant’s cognitive skills (Caparros-Gonzalez et al., 2019), also pointing towards a stimulative role of moderate maternal stress levels.

The fetal stress however was negatively associated with cortical tracking, with lower Hilbert coherence in the familiar rhymes for children with higher hair cortisol. Fetal stress is not just influenced by maternal mood, and therefore the direct assessment is important. The negative impact of fetal stress has been also described by Bergman and colleagues (Bergman et al., 2010), where the levels of cortisol in the amniotic fluid negatively predicted cognitive abilities at 17 months of age. In our case, however, hair cortisol is a marker of chronic fetal stress, reflecting long-term exposure, not the levels at a single timepoint.

Most importantly, the newborn’s ability to track the familiar rhyme (as measured with classical coherence) was positively correlated with the language development at 6 months of age, both with the receptive and expressive scores of the Bayley scales. Earlier studies indicated similar predictive value of cortical tracking (Menn, Ward, et al., 2022; Ní Choisdealbha et al., 2023). This shows how critical the development of non-invasive assessment methods for pre-verbal infants can be.

Measuring the neural speech tracking after birth can become a screening method for infants at risk, or a method to evaluate therapeutic interventions without having to wait until the child speaks to be able to measure the intervention’s effect.

### Methods comparison

All the four methods used in the current study measure the relationship between the brain signal and the auditory stimulus, but they each uncover specific details and measure slightly different aspects of this relationship. This is why results are sometimes different between methods, though most of the time, if there was an effect or a trend, it was present in more than one method, and with the same directionality. By applying multiple methods to the same dataset, the current study offers a fresh perspective on the challenges that infant data analysis can pose. Often studies only report specific methods and it remains unclear whether other methods and parameter settings, such as filters, frequency bands, or transformations, were also tested. Adapting a method is justified, because infants studies are usually heavily contaminated with noise (movement, crying, fussing, external factors, e.g., maternal pulse or heart rate artifacts) which can be different from one setting, sample, or age group to another. Furthermore, interindividual variability at this age is higher than in adults, which increases variance and decreases the study’s power. The expected effects in infants are also quite small, considering the lack of brain maturation, which makes brain activity between small regions less synchronized and more spread over time, leading to a wash-out of the possible differences. All these factors contribute to the weak effects, which might only become significant in a specific constellation of parameters, with a specific method. Therefore, we considered the current comparative report of methods to be useful for future reference and we think that trying (and reporting) more methods should be encouraged in the case of infants’ data.

## Limitations

The most important limitations in our study that lead to the weak effects are the artifacts contamination of the data and high inter-subject variability (large standard deviations), which are common for most studies on young infants. Another limitation could be the (too) fine difference between the original rhymes and their rhythm-manipulated versions. This, however, might be of use in later timepoints of analysis (6-months and 12-months after birth), where the same paradigm is used. At these older ages, the larger frame-study aims to investigate the infants’ ability to identify a correct or incorrect stress on words in their familiar language. Therefore, in the context of the larger study, the paradigm is useful for a dynamic comparison between different ages.

Another possible limitation is that the current study investigates only one-to-one frequency correspondences between EEG and speech envelope, as they are both filtered in the same frequency band and then the measure of interest is analyzed (coherence, MI, TRF etc.). This was done because, as different frequency bands of the EEG track different elements of the language stimulus (Cogan & Poeppel, 2011), we were interested in the neural tracking of different language structures: syllables (within the syllable rate) and words (within the prosodic rate). But this approach might be a limitation, ignoring higher-frequency properties of speech that may induce low-frequency changes in the brain signal. Therefore, further studies should also investigate the relationship between the brain signal and higher frequencies of the speech envelope.

## Conclusion

In the current study, we were able to contribute to the current knowledge on language-specific abilities in healthy 2-week-old newborns. We found that newborns can track a continuous stimulus, as shown by significant cortical tracking, especially at the prosodic (stressed syllables) rate. Importantly, we found that newborns can discriminate between a familiar and unfamiliar rhyme, indicating prenatal learning. Furthermore, violating the phonotactic rules of the familiar (native) language in the backward speech condition, as well as degrading the phonological cues in the low-pass filtered condition, made the cortical tracking more difficult than for the familiar, unmanipulated rhyme. This indicates some degree of phonotactic sensitivity at this young age, i.e., that there is already some knowledge on what syllables should sound like and where they are allowed to be placed in a sentence. Additionally, we found that the recognition of the familiar rhyme was negatively impacted by the manipulation of its rhythm, rendering it more difficult to track. Moreover, in analyzing the data with four common approaches, we highlight the variability of the outcomes and point to the difficulty when it comes to comparing such studies. The correlations of fetal stress, maternal mood, and later infant language development with the newborn’s ability to track the rhymes underline the importance of an early assessment method of language processing. More infant studies using multiple methods are needed in order to come up with a recommendation and agreement how to analyze reproducible speech-envelope data at this early stage.

## Notes

### Competing Interest Statement

The authors have declared no competing interest.

