## Supplemental Material for "Neural speech tracking in newborns: prenatal learning and contributing factors"

For the manuscript: Neural speech tracking in newborns: prenatal learning and contributing factors

### 1. Supplementary methods

#### a. Solution for the markers' synchronization

The audio file included markers for the start of each file as well as for each syllable, discerning between stressed and non-stressed syllables. The script sent the audio data on one of the audio channels, and a high-volume pulse for each marker on the second audio channel. These two audio channels entered a custom-made box that included a SunFounder Mega 2560 Rev3 control board (SunFounder, Shenzhen, China), programmed using the Arduino programming language. The box would to send a 5V pulse to a DIN generator (part of the EGI system) for the high volume pulse from the first channel, and let the audio data from the second channel to go to both channels of the speakers. This ensured high synchronization between the EEG markers and the true auditory stimulus. From previous pilot studies there was an evident delay in the markers that would accumulate over the 1-minute loop of the continuous rhyme and would reach significant lags of tens to hundreds of milliseconds. As this was relevant only for long, continuous auditory files, while most literature uses short stimuli, the synchronization problem had not been previously addressed, which led us to the custom-made solution.

#### b. APICE pipeline steps

1. Demeaning (centering the data around the mean, by substracting the mean of each channel from the respective channel).

2. Low-pass filtering at 40 Hz using a Hamming windowed sinc FIR filter, with a transition band of 10 Hz, passband edge of 40Hz, cutoff frequency of 45 Hz.

3. High-pass filtering at 0.1 Hz, using the same type of filter but with a transition band width of 0.1 Hz, passband edge of 0.1 Hz and cutoff frequency at 0.05 Hz.

4. Identification of artifacts and building of an "Artifacts" matrix of electrodes x sampling points x epochs with 1 indicating bad data and 0 indicating good data for each electrode-sample point. This process involved also multiple steps:

4.1. First step:

a) Rejection of data based on channel correlations

b) Rejection of data based on the power at different frequency bands

c) Rejection of short good segments of data between bad segments of data

4.2. Second step:

a) Rejection of data based on amplitude with a fixed threshold (500 mV)

4.3. Third step is a loop ran twice over the following processes:

a) Rejection of data based on amplitude with a relative threshold (3 interquartile ranges above the 75th percentile)

b) Rejection of data based on variance in a sliding time window, with a relative threshold per electrode

c) Rejection of data based on the running average

4.4. Fourth step is another loop ran twice over the following processes:

a) Rejection of bad data based on amplitude with a relative threshold and average referencing of the data before removing the high amplitude artifacts

b) Rejection of bad data based on variance in a sliding time, after referencing data to the mean of the clean data

c) Rejection of data based on the running average, on data referenced to the mean of the clean data

4.5. Fifth step:

a) Rejecting data based on the variance across electrodes

b) Include short bad data segments between two longer good segments

c) Reject short good data segments between two longer bad segments

5. Identification of artifacts again, this time based on the maximum change in a given time window is larger than a relative threshold. This is applied twice, the second time on data that has been reference to the average of the clean data. Then the short bad segments are included back (marked back as good) and the short good segments are marked as bad.

6. Defining the bad times and bad channels based on the "Artifacts" matrix. Bad times are defined based on the proportion of bad channels during a specific time, and the bad channels are then defined based on the proportion of time during which a channel is bad. The process is iterative, repeating three times, once with a threshold of 70%, once 50%, and lastly 30%. After that, the too short segments marked as bad are marked back as good, and the too short segments marked as good are marked as bad. Finally, a masking window of 0.5s is applied around bad times, marking as bad the half second before and after a bad time.

7. Spatial interpolation of channels that were marked as bad for short periods of time, using spherical spline.

8. Demeaning again

9. Highpass filtering again with 0.1 Hz

10. Independent component analysis (ICA): The data is copied, high-pass filtered at 2Hz, bad samples and bad channels are removes. A first PCA is applied, with 50 components kept in the PCA, and then an ICA. Then, wavelet-thresholding is used to estimate and remove transient artifacts. On the artifacts-free data, a second PCA and ICA are performed. The components to be removed are then identified using the iMARA algorithm [a script accompanying the paper Automatic classification of ICA components from infant EEG using MARA", by Haresign et al, 2021]. In contrast to the MARA algorithm, iMARA was specifically designed for babies and does not base on identification of the same frequency bands as in the adults to discriminate a component as brain activity vs. noise. This is advantageous, since the MARA algorithm uses alpha and beta power to make this discrimination, but infants EEGs are much slower, with more theta and delta power than alpha and beta. After the bad components were identified, the pipeline further estimates artifacts in the data using these components and removes the estimated artifacts from the original (not the copied) data.

11. Another spatial interpolation of the channels not working for a limited period, using spherical spline.

12. Another round of identification of artifacts, similar to step 4, including multiple steps:

12.1. First step: Rejection based on amplitude with a fixed threshold

12.2. Second step is a loop ran twice over the following processes:

a) Rejection of data based on a relative amplitude

b) Rejection of data based on variance over a sliding time window

c) Rejection of data based on the running average

12.3. Third step is also a loop ran twice over the following processes:

a) Rejection based on the relative amplitude, but on data that is referenced to the mean of clean data, and the threshold is applied over all the electrodes, not per electrode

b) Rejection based on variance over a sliding time window, but this time on data referenced to the mean of clean data

c) Rejection based on the running average, but this time on data referenced to the mean of clean data.

d) Rejection based on the variance across electrodes, with a relative threshold defined for all electrodes (three interquartile ranges from the median of all electrodes)

e) Mark short bad segments as good

f) Mark short good segments as bad.

13. Redefine the Bad Times and Bad Segments just as in step 6, based on the last changes to the "Artifacts" matrix.

##### c. Matlab code for the implementation of the Hilbert coherence

```
% data = bandpass-filtered, Hilbert-transformed data
[nc, ns, nt] = size(data); % data format: channels x samples x trials
phase = angle(data); % extract phase information
amp = abs(data); % extract amplitude information (absolute value)
signal_pow = amp.^2;
h_coh = zeros(nc, nt);
c2 = 83; % the audio channel in our data was the 83rd channel
for t=1:nt
    for c1 = 1:nc
         phase_diff = phase(c1,:,t) - phase(c2,:,t);
         amp_prod = amp(c1,:,t).*amp(c2,:,t);
         num = abs(sum(amp_prod.*exp(1i*phase_diff)));
        %num = abs(sum(data(c1,:,t).*conj(data(c2,:,t)))); % this is the
        %same as above, looks just closer as the formula. It gives the
        %exact same values, but it is a bit slower.
        denom = sqrt(sum(signal_pow(c1,:,t)).* sum(signal_pow(c2,:,t)));
        h_coh(c1, t) = num/denom;
    end
end
```

#### 2. Supplementary results

##### 2.1. Results of the classical coherence analysis

The comparison with surrogate coherence distributions showed that …% of coherence values were significant.

###### 2.1.1. Analysis of all coherence values (not just significant ones)

###### *2.1.1.1. Average coherence over the frequencies of interest as the dependent variable*

For the prosodic rate, the Wilcoxon signed rank test revealed no rhyme effect (no difference between conditions 1 and 5), no familiarity effect and no group effect. Similarly, for the prosodic rate there were also no effects of rhyme or group (see table below), but there was a familiarity effect, p = 0.048, with lower coherence values for the familiar rhyme (mean = 0.111) than for the unfamiliar rhyme (mean = 0.116). We were interested in localised effects in each group, so we also tested the familiarity and rhyme effects in each group separately, and found a familiarity effect in group 1, prosodic stress rate, with slightly higher coherence values for the unfamiliar rhyme (coh = 0.119) than for the familiar rhyme (coh = 0.108) (see table below and Fig. 1, panel A), but no familiarity effect for Group 2.

For the language, rhythm and phonological paradigms, the Wilcoxon signed rank tests comparing the familiar unmanipulated rhyme to the familiar manipulated rhymes over all participants showed no effects, and separate analyses for each group also showed no effects (see table below).

**Table S1: Results of classical coherence analysis – variable = average coherence over the frequency range of interest (prosodic or syllable rate)**

| **Data** | **Dependent variable** | **Compared conditions** | **Prosodic rate** | **Syllable rate** |
|---|---|---|---|---|
| All participants N = 50 | Average coherence | R1 vs R2 (rhyme effect) | V = 708, p-value = 0.4992 | V = 662, p-value = 0.8168 |
| | | Familiarity effect | V = 432, p-value = 0.04782 | V = 563, p-value = 0.475 |
| | | Gr 1 vs Gr 2 (group effect) | W = 1373, p-value = 0.3668 | W = 1395, p-value = 0.2916 |
| Group 1 (N = 23) | Average coherence | Familiarity effect | V = 72, p-value = 0.04488 | V = 111, p-value = 0.4274 |
| Group 2 (N = 27) | Average coherence | Familiarity effect | V = 154, p-value = 0.4133 | V = 181, p-value = 0.8593 |
| All participants N = 50 | Average coherence | Language effect | V = 603, p-value = 0.7428 | V = 777, p-value = 0.1797 |
| Group 1 (N = 22) | Average coherence | Language effect | V = 159, p-value = 0.3053 | V = 169, p-value = 0.1762 |
| Group 2 (N = 28) | Average coherence | Language effect | V = 152, p-value = 0.2545 | V = 227, p-value = 0.5979 |
| All participants N = 51 | Average coherence | Rhythm effect | V = 579, p-value = 0.4338 | V = 825, p-value = 0.1301 |
| Group 1 (N = 22) | Average coherence | Rhythm effect | V = 115, p-value = 0.7262 | V = 159, p-value = 0.3053 |
| Group 2 (N = 29) | Average coherence | Rhythm effect | V = 181, p-value = 0.442 | V = 272, p-value = 0.247 |
| All participants (N = 49) | Average coherence | Phonological effect | V = 516, p-value = 0.3427 | V = 550, p-value = 0.5407 |
| Group 1 (N = 22) | Average coherence | Phonological effect | V = 121, p-value = 0.8736 | V = 119, p-value = 0.8237 |
| Group 2 (N = 27) | Average coherence | Phonological effect | V = 141, p-value = 0.2584 | V = 163, p-value = 0.546 |

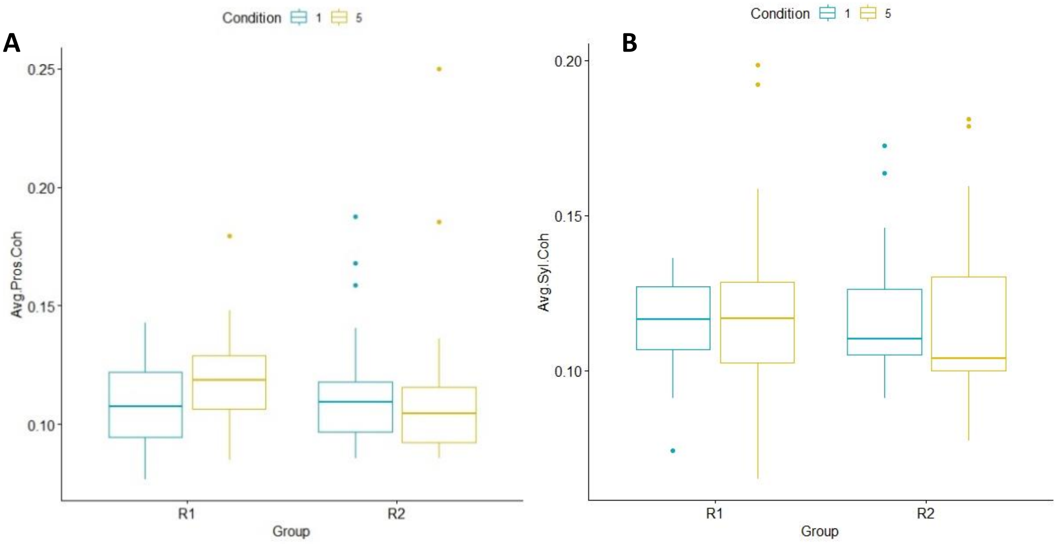

**Fig. S1: Classical coherence values (average values)** in conditions 1 (Butzemann unmanipulated) and 5 (Es war eine Mutter unmanipulated), in Groups 1 (familiar with Rhyme 1) and Group 2 (familiar with Rhyme 2), for prosodic (panel A) and syllable rate (panel B). Note the trend for a higher coherence for the unfamiliar rhyme in Group 1 in the prosodic frequency rate (panel A, left).

##### *2.1.1.2. Maximal coherence over the frequencies of interest as the dependent variable*

We took the maximal coherence value over the frequency interval of interest (prosodic or syllable rate) for each participant and condition and used it as a dependent variable. The Wilcoxon signed rank tests found no group effect, no rhyme effect, and no familiarity effect. In the language paradigm, there was a trend for a higher coherence in the unmanipulated rhyme (coh = 0.147) than in the language manipulated rhyme (coh = 0.139) in the prosodic rate, p = 0.070. The trend was driven by the first group, where the trend was similar, with p = 0.068 (see table below), while in the second group there was no effect. In the rhythm paradigm there was also a significant effect (also driven by group 1), in the prosodic rate, with higher coherence for the unmanipulated rhyme (coh = 0.147) than for the rhythm-manipulated rhyme (coh = 0.135).

**Table S2: Results of classical coherence analysis – variable = maximal coherence over the frequency range of interest (prosodic or syllable rate)**

| **Data** | **Dependent variable** | **Compared conditions** | **Prosodic rate** | **Syllable rate** |
|---|---|---|---|---|
| All participants N = 50 | Maximal coherence | R1 vs R2 (rhyme effect) | V = 641, p-value = 0.9769 | V = 732, p-value = 0.3642 |
| | | Familiarity effect | V = 601, p-value = 0.7282 | V = 546, p-value = 0.3797 |
| | | Gr 1 vs Gr 2 (group effect) | W = 1253, p-value = 0.9421 | W = 1269, p-value = 0.8546 |
| Group 1 (N = 23) | Maximal coherence | Familiarity effect | V = 128, p-value = 0.7768 | V = 89, p-value = 0.1424 |
| Group 2 (N = 27) | Maximal coherence | Familiarity effect | V = 180, p-value = 0.8408 | V = 187, p-value = 0.9717 |
| All participants N = 50 | Maximal coherence | Language effect | V = 826, p-value = 0.06955 | V = 677, p-value = 0.7066 |
| Group 1 (N = 22) | Maximal coherence | Language effect | V = 183, p-value = 0.06844 | V = 150, p-value = 0.4628 |

| | | | | |
|---|---|---|---|---|
| Group 2 (N = 28) | Maximal coherence | Language effect | V = 239, p-value = 0.4248 | V = 200, p-value = 0.9553 |
| All participants N = 51 | Maximal coherence | Rhythm effect | V = 907, p-value = 0.02246 | V = 653, p-value = 0.929 |
| Group 1 (N = 22) | Maximal coherence | Rhythm effect | V = 180, p-value = 0.08541 | V = 117, p-value = 0.7745 |
| Group 2 (N = 29) | Maximal coherence | Rhythm effect | V = 290, p-value = 0.1207 | V = 222, p-value = 0.9321 |
| All participants (N = 49) | Maximal coherence | Phonological effect | V = 565, p-value = 0.643 | V = 605, p-value = 0.945 |
| Group 1 (N = 22) | Maximal coherence | Phonological effect | V = 140, p-value = 0.6789 | V = 113, p-value = 0.6789 |
| Group 2 (N = 27) | Maximal coherence | Phonological effect | V = 157, p-value = 0.4553 | V = 195, p-value = 0.8966 |

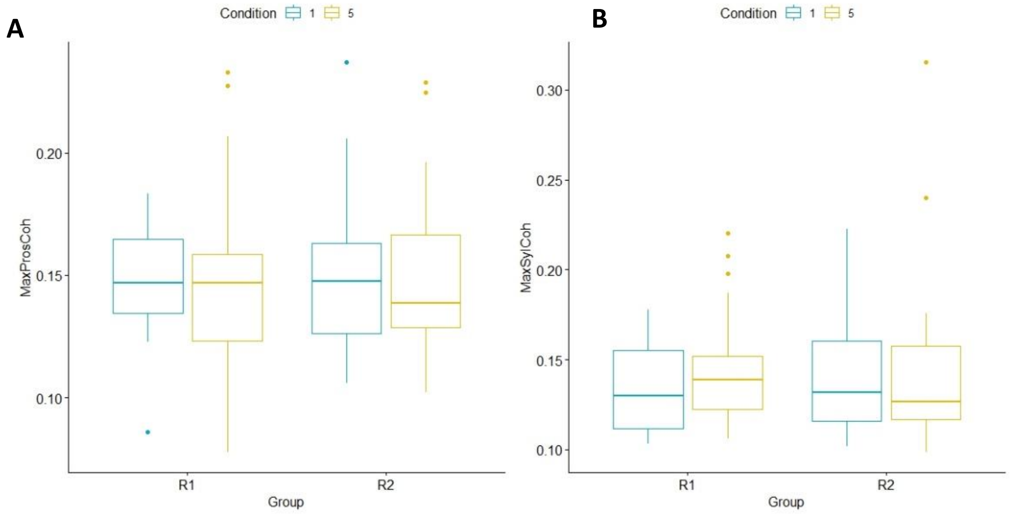

**Fig. S2: : Classical coherence values (maximum values)** in conditions 1 (Butzemann unmanipulated) and 5 (Es war eine Mutter unmanipulated), in groups R1 (familiar with Rhyme 1) and R2 (familiar with Rhyme 2), for prosodic (panel A) and syllable rate (panel B). Note that for the prosodic rate, there is no significant difference between the two rhymes in group 1 anymore.

###### *2.1.1.3. Average of highest 2 coherences over the frequencies of interest as the dependent variable*

We took the highest two coherence values over each frequency interval of interest (prosodic or syllable rate) and used their average as a dependent variable. Wilcoxon signed rank tests found no group or rhyme effects, but there was a trend for a familiarity effect in the syllable rate, with lower coherence in the familiar rhyme (mean = 0.126) than in the unfamiliar rhyme (mean = 0.132), p = 0.065. There was also a trend (p = 0.074) for a language effect in Group 1 with slightly higher coherence values for the unmanipulated condition (coh = 0.134) than for the language-manipulated condition (coh = 0.124). In the rhythm paradigm, there was an effect (mainly driven by Group 1), with higher coherence values for the unmanipulated rhyme (coh = 0.134) than for the rhythm-manipulated rhyme (coh = 0.126), p = 0.041 (see table below)

**Table S3: Results of classical coherence analysis – variable = average of the two highest coherence values in the frequency range of interest (prosodic or syllable rate)**

| Data | Dependent variable | Compared conditions | Prosodic rate | Syllable rate |
|---|---|---|---|---|
| All participants N = 50 | Average of two max. coherences | R1 vs R2 (rhyme effect) | V = 688, p-value = 0.6293 | V = 596, p-value = 0.6923 |

| | | | | |
|---|---|---|---|---|
| | | Familiarity effect | V = 560, p-value = 0.4573 | V = 446, p-value = 0.06522 |
| | | Gr 1 vs Gr 2 (group effect) | W = 1354, p-value = 0.4406 | W = 1335, p-value = 0.5223 |
| Group 1 (N = 23) | Average of two max. coherences | Familiarity effect | V = 109, p-value = 0.3931 | V = 97, p-value = 0.2226 |
| Group 2 (N = 27) | Average of two max. coherences | Familiarity effect | V = 184, p-value = 0.9153 | V = 130, p-value = 0.1624 |
| All participants N = 50 | Average of two max. coherences | Language effect | V = 813, p-value = 0.09116 | V = 630, p-value = 0.9461 |
| Group 1 (N = 22) | Average of two max. coherences | Language effect | V = 182, p-value = 0.07378 | V = 150, p-value = 0.4628 |
| Group 2 (N = 28) | Average of two max. coherences | Language effect | V = 229, p-value = 0.567 | V = 173, p-value = 0.5076 |
| All participants N = 51 | Average of two max. coherences | Rhythm effect | V = 881, p-value = 0.04148 | V = 614, p-value = 0.6494 |
| Group 1 (N = 22) | Average of two max. coherences | Rhythm effect | V = 163, p-value = 0.2479 | V = 112, p-value = 0.6556 |
| Group 2 (N = 29) | Average of two max. coherences | Rhythm effect | V = 288, p-value = 0.1316 | V = 205, p-value = 0.7983 |
| All participants (N = 49) | Average of two max. coherences | Phonological effect | V = 561, p-value = 0.6149 | V = 561, p-value = 0.6149 |
| Group 1 (N = 22) | Average of two max. coherences | Phonological effect | V = 121, p-value = 0.8736 | V = 122, p-value = 0.8987 |
| Group 2 (N = 27) | Average of two max. coherences | Phonological effect | V = 159, p-value = 0.4846 | V = 170, p-value = 0.6617 |

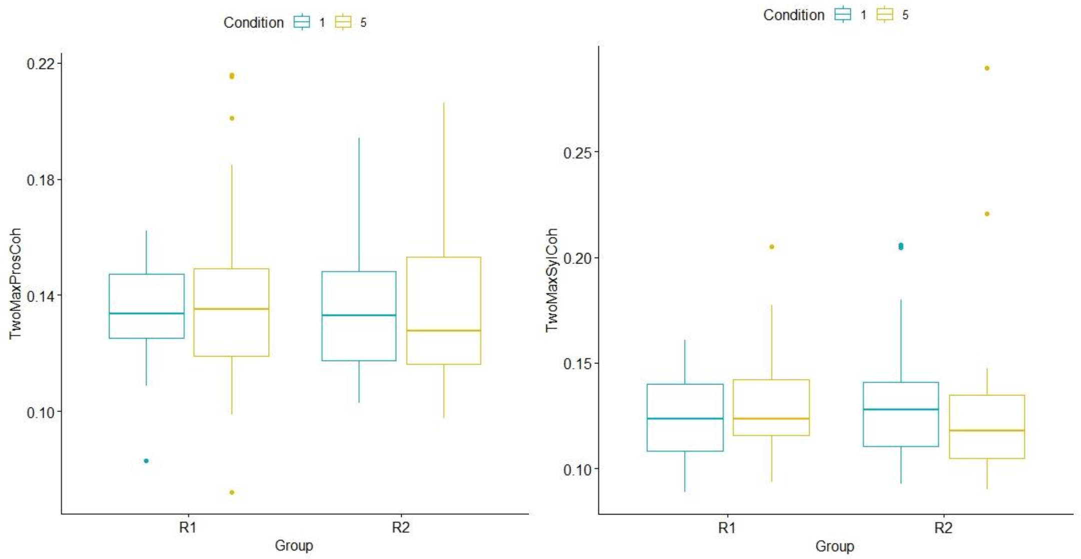

**Fig. S3: Classical coherence (average of two maximums)** in conditions 1 (*Butzemann* unmanipulated) and 5 (*Es war eine Mutter* unmanipulated) in groups R1 (familiar with Rhyme 1) and R2 (familiar with Rhyme 2), for prosodic (panel A) and syllable rate (panel B). Note a trending lower coherence at the syllable rate for the familiar rhyme (condition 1 in Group R1 and condition 5 in Group R2) than for the unfamiliar rhyme.

##### 2.1.2. Analysis of only the significant coherence values

Out of 274 files (participant*condition), 91 (33%) had at least one significant coherence value in the prosodic rate and 104 (38%) had at least one significant coherence value in the syllable rate. The not significant values were discarded and the significant values for each frequency interval were averaged. This average (or, in many cases, single) significant value for each rate (syllable or prosodic), participant, and condition was further used in the analysis.

Rhyme and familiarity effects: for the prosodic rate, 6 participants had significant coherence in both the familiar and the unfamiliar rhyme conditions. For the syllable rate, 12 participants had significant coherence in both conditions. Wilcoxon signed rank tests showed no rhyme effects for either frequency interval, but there was a familiarity effect in the prosodic rate (V = 0, p = 0.031), with a lower coherence for the familiar rhyme (mean = 0.092) than for the unfamiliar rhyme (coh = 0.130). The effect was mainly due to Group 2, because there was just one participant from Group 1 in this sample.

Group effects: for the prosodic rate this could not be tested, because of the small sample size (Wilcoxon rank sum tests for unequal groups require group sizes of at least 5 and 3 and a total sample size of 8 [Dwivedi et al., 2017]). For the syllable rate, the Wilcoxon rank sum test (independent variables) was not significant. There were no language, rhythm, or familiarity effects, in many cases the sample sizes being too small for analysis (see table below).

**Table S4: Results of classical coherence analysis on just the significant coherence values – variable = average coherence over the frequency range of interest (prosodic or syllable rate)**

| Data | Dependent variable | Compared conditions | Prosodic rate | Syllable rate |
|---|---|---|---|---|
| All participants n = 6 (pros), n = 12 (syl) | Average significant coherence | R1 vs R2 (rhyme effect) | V = 5, p-value = 0.3125 | V = 44, p-value = 0.7334 |
| | | Familiarity effect | V = 0, p-value = 0.03125 | V = 26, p-value = 0.3394 |
| | | Gr 1 vs Gr 2 (group effect) | Sample too small | W = 73, p-value = 0.8859 |
| Group 1, n = 1 (pros), n = 7 (syl) | Average sig. coherence | Familiarity effect | Sample too small | V = 8, p-value = 0.375 |
| Group 2, n = 5 (pros), n = 5 (syl) | Average sig. coherence | Familiarity effect | V = 0, p-value = 0.0625 | V = 6, p-value = 0.8125 |
| All participants n = 8 (pros), n = 6 (syl) | Average sig. coherence | Language effect | V = 18, p-value = 1 | V = 11, p-value = 1 |
| Group 1, n = 3 (pros), n = 5 (syl) | Average sig. coherence | Language effect | V = 3, p-value = 1 | V = 11, p-value = 0.4375 |
| Group 2, n = 5 (pros), n = 1 (syl) | Average sig. coherence | Language effect | V = 9, p-value = 0.8125 | Sample too small |
| All participants, n = 4 (pros), n = 4 (syl) | Average sig. coherence | Rhythm effect | V = 2, p-value = 0.375 | V = 9, p-value = 0.25 |
| Group 1, n = 2 (pros), n = 3 (syl) | Average sig. coherence | Rhythm effect | Sample too small | V = 5, p-value = 0.5 |
| Group 2, n = 2 (pros), n = 1 (syl) | Average sig. coherence | Rhythm effect | Sample too small | Sample too small |
| All participants, n = 2 (pros), n = 6 (syl) | Average sig. coherence | Phonological effect | Sample too small | V = 11, p-value = 1 |
| Group 1, n = 1 (pros), n = 4 (syl) | Average sig. coherence | Phonological effect | Sample too small | V = 7, p-value = 0.625 |

| | | | | |
|---|---|---|---|---|
| Group 2, n = 1 (pros), n = 2 (syl) | Average sig. coherence | Phonological effect | Sample too small | Sample too small |

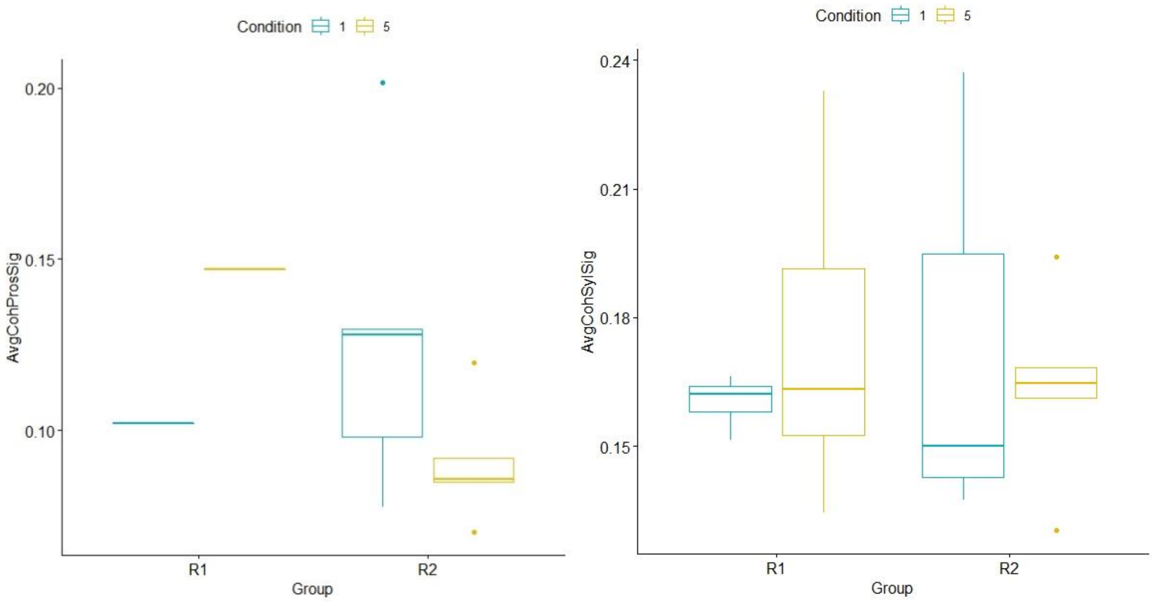

**Fig. S4: Classical coherence (average coherence, analysis of just the significant coherence values)** in conditions 1 (*Butzemann* unmanipulated) and 5 (*Es war eine Mutter* unmanipulated) in groups R1 (familiar with Rhyme 1) and R2 (familiar with Rhyme 2), for prosodic (panel A) and syllable rate (panel B). Note the lower coherence at the prosodic rate for the familiar rhyme (condition 1 in Group R1 and condition 5 in Group R2) than for the unfamiliar rhyme.

#### 2.2. Hilbert coherence analysis

##### 2.2.1. Analysis of all values

###### *2.2.1.1. Analysis of all Hilbert coherence values for prosodic rate*

Wilcoxon signed rank tests found a significant rhyme effect, $p < 0.001$, with higher coherence for the R2 rhyme (Es war eine Mutter) than for the R1 rhyme (Butzemann). There was no effect of familiarity or group. In both groups, there was a significant rhyme effect with higher coherence for the R2 rhyme. Over all participants, also the language effect was significant, $p = 0.043$ (mostly due to Group 2), with higher coherence for the language-manipulated rhyme (mean = 0.439) than for the original familiar rhyme (mean = 0.432). There were no effects in the rhythm paradigm, but there was a trend ($p = 0.064$) in the phonological paradigm, with higher coherence values for the original rhyme (mean = 0.431) than for the phonologically manipulated rhyme (mean = 0.424).

###### *2.2.1.2. Analysis of all Hilbert coherence values for syllable rate*

Wilcoxon signed rank tests showed no rhyme or familiarity effect, and Wilcoxon rank sum test showed no group effect (see Fig. S5 and Table S5). There were no significant effects for any of the other paradigms.

**Table S5: Results of Hilbert coherence analysis (on all values, not just the significant ones)**

| **Data** | **Dependent variable** | **Compared conditions** | **Prosodic rate** | **Syllable rate** |
|---|---|---|---|---|

| | | | | |
|---|---|---|---|---|
| All participants n = 50 | Average coherence | R1 vs R2 (rhyme effect) | V = 1275, p-value = 7.79e-10 | V = 773, p-value = 0.1925 |
| | | Familiarity effect | V = 714, p-value = 0.4632 | V = 705, p-value = 0.5178 |
| | | Gr 1 vs Gr 2 (group effect) | W = 1258, p-value = 0.9146 | W = 1220, p-value = 0.8818 |
| Group 1, n = 23 | Average coherence | Familiarity effect | V = 0, p-value = 2.384e-07 | V = 121, p-value = 0.6221 |
| Group 2, n = 27 | Average coherence | Familiarity effect | V = 0, p-value = 1.49e-08 | V = 132, p-value = 0.1775 |
| All participants n = 50 | Average coherence | Language effect | V = 409, p-value = 0.04274 | V = 560, p-value = 0.608 |
| Group 1, n = 22 | Average coherence | Language effect | V = 84, p-value = 0.2877 | V = 118, p-value = 0.9457 |
| Group 2, n = 28 | Average coherence | Language effect | V = 126, p-value = 0.08145 | V = 168, p-value = 0.438 |
| All participants, n = 51 | Average coherence | Rhythm effect | V = 624, p-value = 0.7182 | V = 654, p-value = 0.9365 |
| Group 1, n = 22 | Average coherence | Rhythm effect | V = 140, p-value = 0.6789 | V = 159, p-value = 0.3053 |
| Group 2, n = 29 | Average coherence | Rhythm effect | V = 176, p-value = 0.3808 | V = 170, p-value = 0.3145 |
| All participants, n = 49 | Average coherence | Phonological effect | V = 799, p-value = 0.06393 | V = 600, p-value = 0.9058 |
| Group 1, n = 22 | Average coherence | Phonological effect | V = 169, p-value = 0.1762 | V = 106, p-value = 0.5235 |
| Group 2, n = 27 | Average coherence | Phonological effect | V = 253, p-value = 0.1286 | V = 211, p-value = 0.6109 |

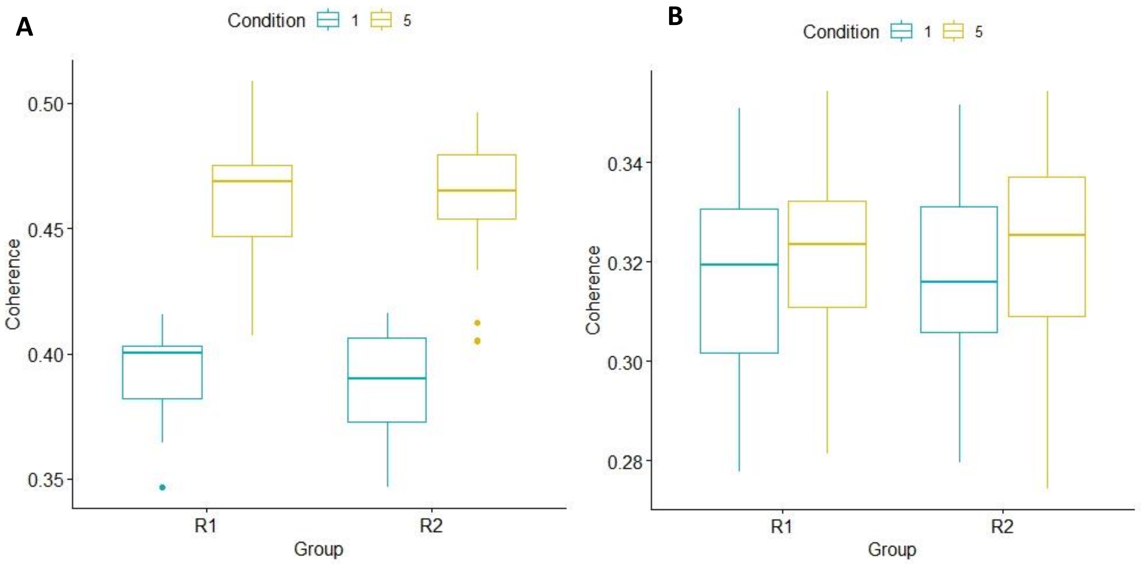

**Fig. S5: Hilbert cohernce for prosodic rate (panel A) and syllable rate (panel B) over all files**, n = 50 (not just the significant ones).

##### 2.2.2. Analysis of only significant values

###### *2.2.2.1. Analysis of significant Hilbert coherence for prosodic rate*

Out of 274 files, 160 (58.4%) had significant Hilbert coherence values in the prosodic rate. Wilcoxon signed rank tests showed a significant effect of rhyme (p = 0.004, V = 45), with higher coherence values for Rhyme 2, but no effect for familiarity or group. For the language, rhythm, and phonological paradigm there were no significant effects (see table below).

###### *2.2.2.2. Analysis of significant Hilbert coherence for syllable rate*

Out of 274 files (participant*condition), 59 (21.5%) had significant Hilbert coherence values in the syllable rate after comparing to the distribution made with surrogate data.

Wilcoxon signed rank tests on only these significant values found no effect of rhyme or familiarity, and the Wilcoxon rank sum test found no group effect (Fig. S6). For the phonological paradigm there was also no significant result, and other comparisons were not possible due to the small or inexistent sample size (see Table S6).

**Table S6: Results of Hilbert coherence analysis (on just the significant values)**

| **Data** | **Dependent variable** | **Compared conditions** | **Prosodic rate** | **Syllable rate** |
|---|---|---|---|---|
| All participants n = 9 (pros), n = 4 (syl) | Average significant coherence | R1 vs R2 (rhyme effect) | V = 45, p-value = 0.003906 | V = 3, p-value = 0.625 |
| | | Familiarity effect | V = 27, p-value = 0.6523 | V = 6, p-value = 0.875 |
| | | Gr 1 vs Gr 2 (group effect) | W = 32, p-value = 0.5148 | W = 6, p-value = 0.6857 |
| Group 1, n = 4 (pros), n = 2 (syl) | Average sig. coherence | Familiarity effect | V = 0, p-value = 0.125 | Sample too small |
| Group 2, n = 5 (pros), n = 2 (syl) | Average sig. coherence | Familiarity effect | V = 0, p-value = 0.0625 | Sample too small |
| All participants n = 22 (pros), n = 1 (syl) | Average sig. coherence | Language effect | V = 98, p-value = 0.3705 | Sample too small |
| Group 1, n = 1 (pros), n = 0 (syl) | Average sig. coherence | Language effect | Sample too small | Sample too small |
| Group 2, n = 21 (pros), n = 1 (syl) | Average sig. coherence | Language effect | V = 92, p-value = 0.4319 | Sample too small |
| All participants, n = 24 (pros), n = 2 (syl) | Average sig. coherence | Rhythm effect | V = 140, p-value = 0.7898 | Sample too small |
| Group 1, n = 2 (pros), n = 0 (syl) | Average sig. coherence | Rhythm effect | Sample too small | Sample too small |
| Group 2, n = 22 (pros), n = 2 (syl) | Average sig. coherence | Rhythm effect | V = 123, p-value = 0.924 | Sample too small |
| All participants, n = 15 (pros), n = 4 (syl) | Average sig. coherence | Phonological effect | V = 82, p-value = 0.2293 | V = 7, p-value = 0.625 |
| Group 1, n = 0 (pros), n = 1 (syl) | Average sig. coherence | Phonological effect | Sample too small | Sample too small |
| Group 2, n = 15 (pros), n = 3 (syl) | Average sig. coherence | Phonological effect | V = 82, p-value = 0.2293 | V = 4, p-value = 0.75 |

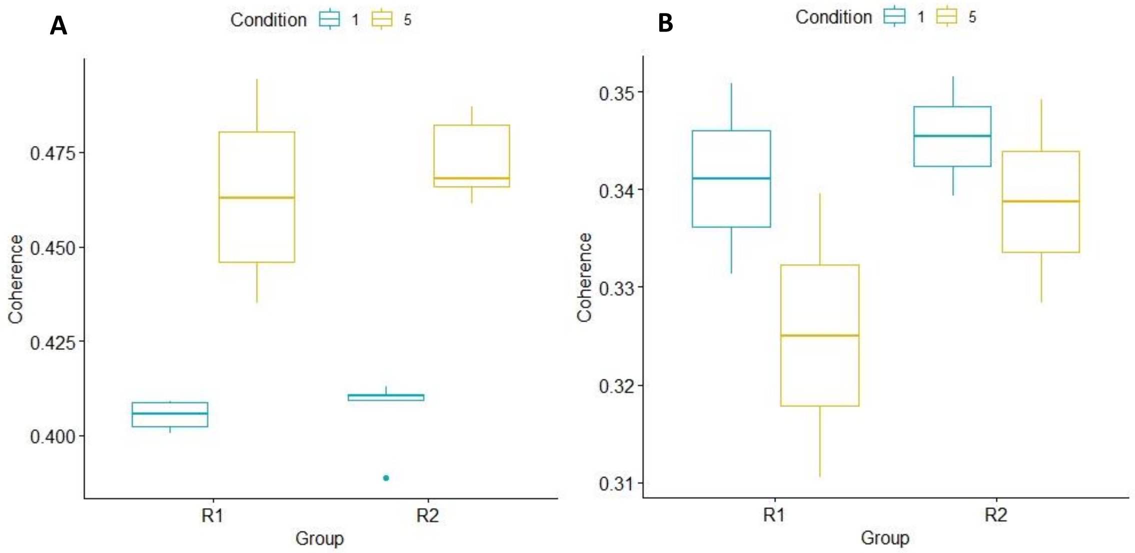

**Fig. S6: Hilbert coherence with only the significant values for the prosodic (panel A, n = 9) and syllable rate (panel B, n = 4)**.

#### 2.3. mTRF analysis

##### 2.3.1. mTRF models between EEG and the speech envelope

###### *2.3.1.1. Bandpass filtered over 0.5 – 2 Hz*

Cluster based permutation (CBP) analysis showed no significant clusters for this frequency band.

The analysis of the r correlation values between the predicted and the real signal (using Spearman correlation) was performed in R Studio. Wilcoxon signed rank tests showed a rhyme effect, p = 0.007, based mainly on Group 1, with a higher r for R1 than for R2, but no other effects (see Table S7 and Fig. S7).

When analyzing the squared r (mean r of each participant raised to the power of 2), there remains only a trend (p = 0.097) for a language effect in group 1, with stronger correlation for the familiar language (mean r_sqrd = 0.0072) than for the language-manipulated rhyme (mean r_sqrd = 0.0028).

When analyzing the absolute r (absolute value of the mean r), there are two trends in Group 1, one for a language effect, towards higher |r| values for the familiar language (mean|r| =0.069) than for the language manipulated rhyme (mean |r| =0.40), and one for rhythm effect, towards higher |r| values for the original unmanipulated rhyme (mean|r| =0.061) than the rhythm manipulated rhyme (mean|r| = 0.040).

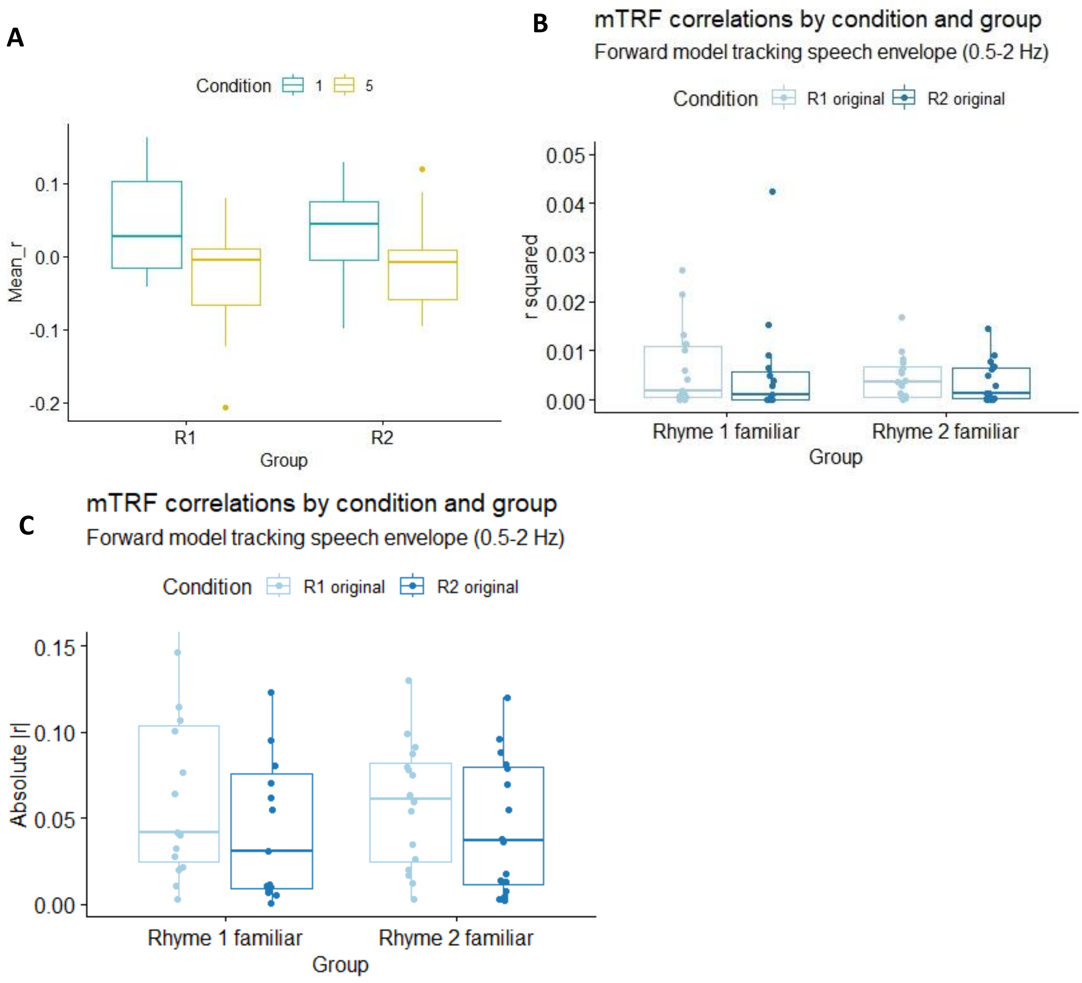

**Fig. S7: The correlation coefficients of the mTRF models for speech envelope, 0.5-2Hz**: comparison of the two groups and the two rhymes. Note a higher *r* for rhyme 1 in both groups in panel A. Panel A = with r, Panel B with r squared, Panel C with absolute value of r.

##### *2.3.1.2. Bandpass filtered over 0.5 – 4 Hz*

For the mTRF models computed on data bandpass filtered over 0.5-4 Hz there was a significant rhyme effect in all participants (but mainly based on Group 1), shown by a middle frontal and left central cluster, with stronger activation for rhyme 2, between 450 and 750ms (Fig. S8). In Group 1, the same contrast (rhyme 1, familiar, vs rhyme 2, unfamiliar) led to a trending cluster left central-parietal with a stronger activation in the unfamiliar rhyme between 450 and 800ms (Fig. S9). There was also a trend for a language effect, shown by a cluster middle and left central-parietal, between 750 and 900ms, indicating higher activation for the unfamiliar language (Fig. S10).

The analysis of the r correlation values between the predicted and the real signal (using Spearman correlation) was performed in R Studio. Wilcoxon signed rank tests showed similar results to the data filtered 0.5-2Hz: an effect of rhyme in all participants and in group 1, with the first rhyme having higher r values (mean r = 0.036) than the second rhyme (mean r = -0.018).

When analysing r squared, there are two trends showing up in Group 1: one for a language effect, with a higher r for the original (r squared = 0.006) than for the manipulated rhyme (squared r = 0.002) and one for a phonological effect, with higher r for the original (squared r = 0.006) than for the manipulated rhyme (squared r = 0.002).

When analysing |r|, there is a language effect in group 1, with higher r values for the original rhyme (|r| = 0.062) than for the manipulated rhyme (|r| = 0.036).

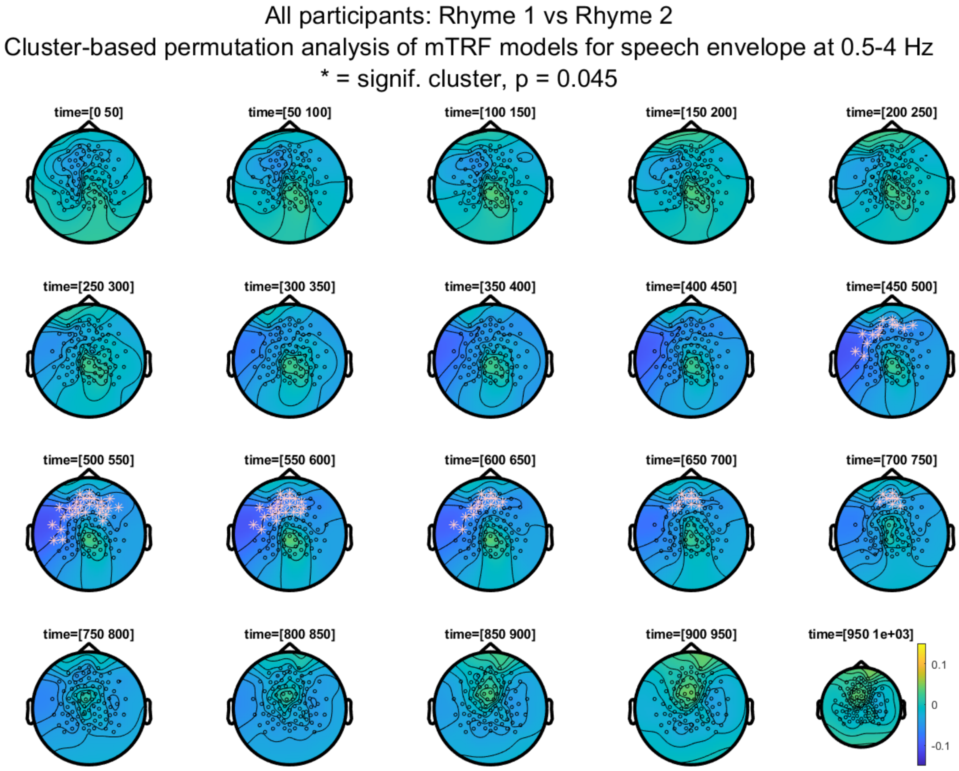

**Fig. S8: CBP analysis of mTRF models for speech envelope at 0.5-4Hz, in all participants, showing the rhyme effect (contrast between Rhyme 1 and Rhyme 2)**. Note a stronger activation in Rhyme 2 midle frontal and left central between 450-750ms.

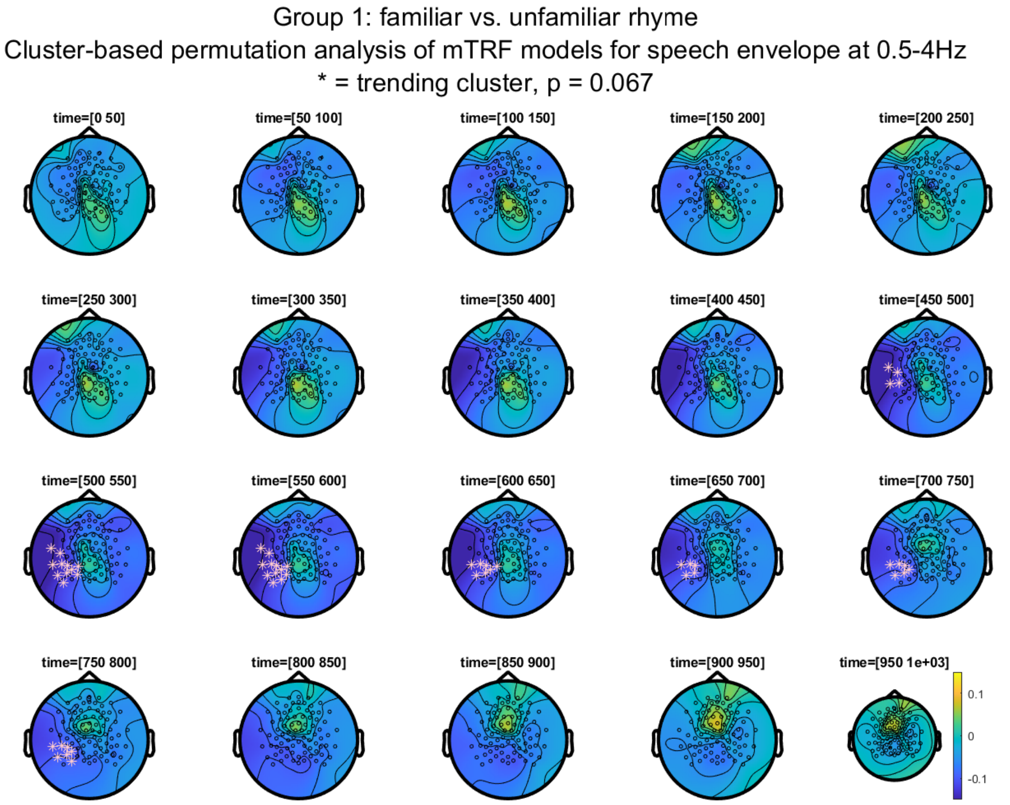

**Fig. S9: CBP analysis of mTRF models for speech envelope at 0.5-4Hz, in Group 1, contrast between unfamiliar and familiar rhyme**. Note a stronger activation in the unfamiliar rhyme between 450 and 800ms left central-parietal.

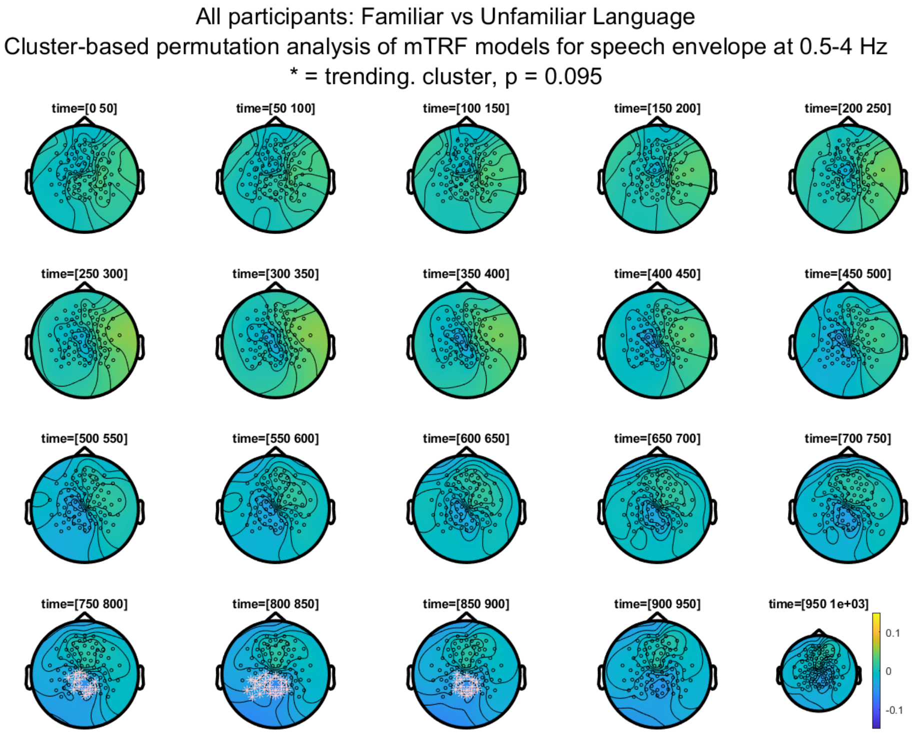

**Fig. S10: CBP analysis (0.5 – 4Hz), in all participants, contrast between familiar and unfamiliar language.** Note a trending cluster middle and left central-parietal, between 750 and 900ms, indicating higher activation for the unfamiliar language.

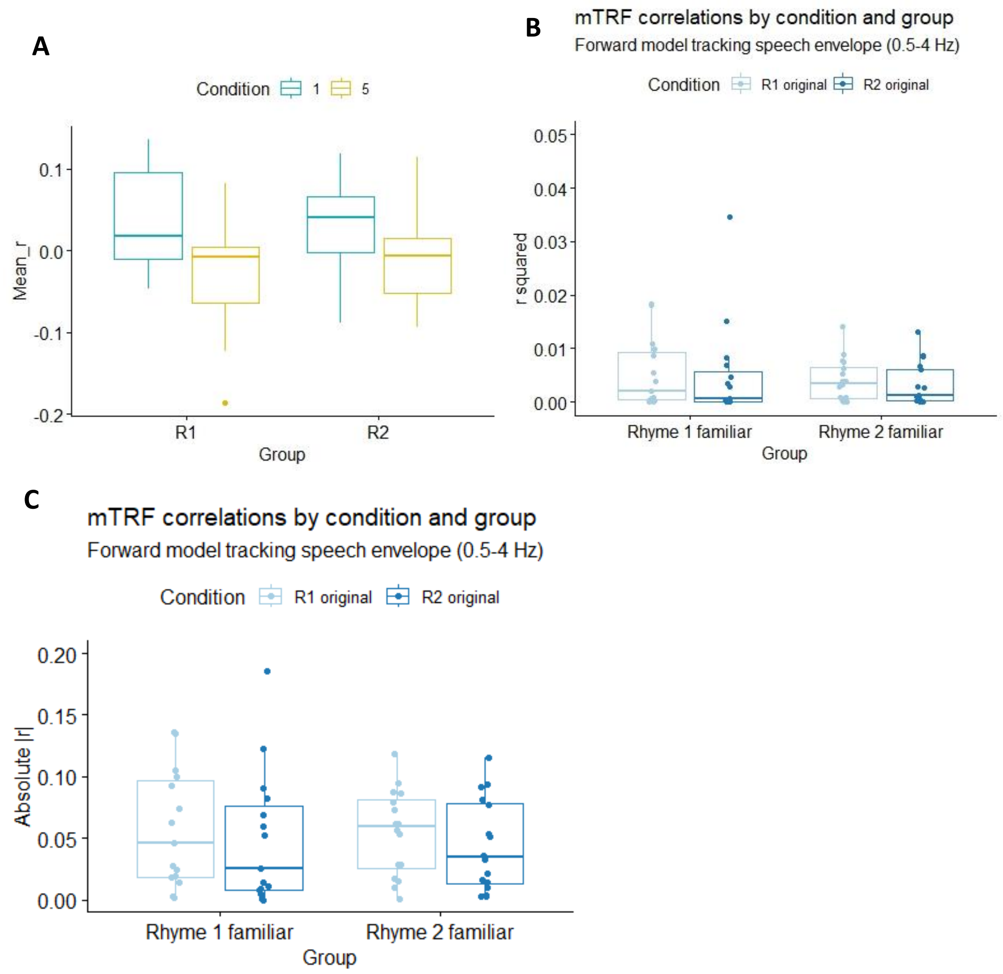

**Figure S11: The correlation coefficients of the mTRF models for speech envelope, 0.5-4Hz**: comparison of the two groups and the two rhymes. Note a higher *r* for rhyme 1 in both groups in panel A. Panel A = with r, Panel B with r squared, Panel C with absolute value of r.

##### *2.3.1.3. Bandpass filtered over 0.5 – 10 Hz*

For the mTRF models computed on data bandpass filtered over 0.5-10 Hz, there was a rhyme effect with a middle frontal and left central cluster between 450ms and 750ms after stimulus onset, p = 0.046 (Fig. S13). This effect was based mainly on Group 1, where there was a trending cluster in the left centro-parietal region (Fig. S12) between 500ms and 700ms from stimulus onset. As the difference is made by subtracting the familiar from the unfamiliar rhyme, and blue indicates negative values, this figure shows a higher activation in the cluster region for the familiar rhyme than for the unfamiliar rhyme (trending, p = 0.083).

The analysis of the r correlation values between the predicted and the real signal (using Spearman correlation) was performed with Wilcoxon signed rank tests and showed similar results to the data in the other frequency bands. There was an effect of rhyme in all participants and in group 1, with the first rhyme having higher r values (mean r = 0.035) than the second rhyme (mean r = -0.017).

When analysing r squared, similar to the other frequency bands, the rhyme effect disappears and a trend for a language effect in group 1 appears, with higher r squared values for the original rhyme (r sqrd = 0.0058) than for the language manipulated rhyme (r sqrd = 0.0024).

The analysis of |r| shows, like in the other frequency bands, a trend for a language effect in group 1, with a higher |rZ for the original rhyme (|r| = 0.061) than for the rhythm-manipulated rhyme (|r| = 0.036).

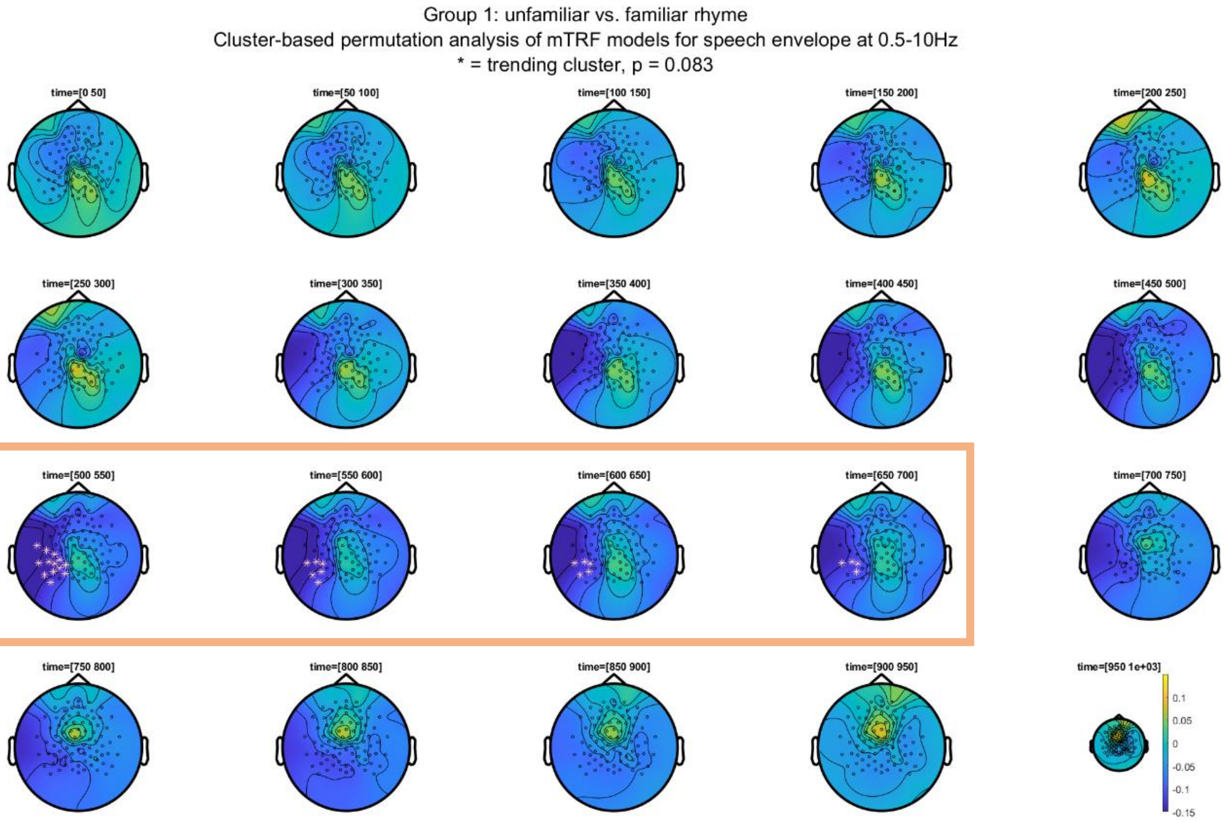

**Fig. S12: CBP analysis of TRF (speech envelope, 0.5-10Hz, Group 1, rhyme familiarity effect: unfamiliar – familiar rhyme)**. Note a trending cluster in the left centro-parietal region (pink asterix) between 500ms and 700ms from stimulus onset (orange rectangle), showing a higher activation for the familiar than the unfamiliar rhyme.

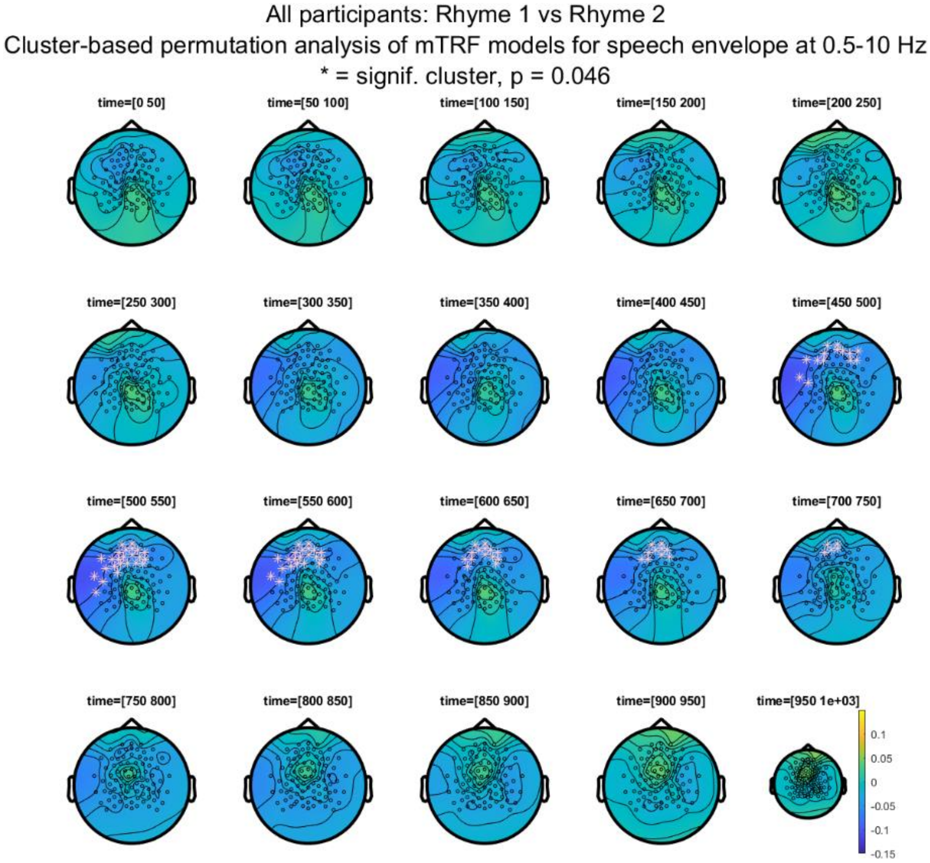

**Fig. S13: CBP analysis of mTRF models for speech envelope at 0.5-10Hz, rhyme effect (Rhyme 1 vs. rhyme 2 in all participants).** Note the stronger activation in Rhyme 2 than in Rhyme 1. Pink asterix = the signif. middle frontal and left central cluster.

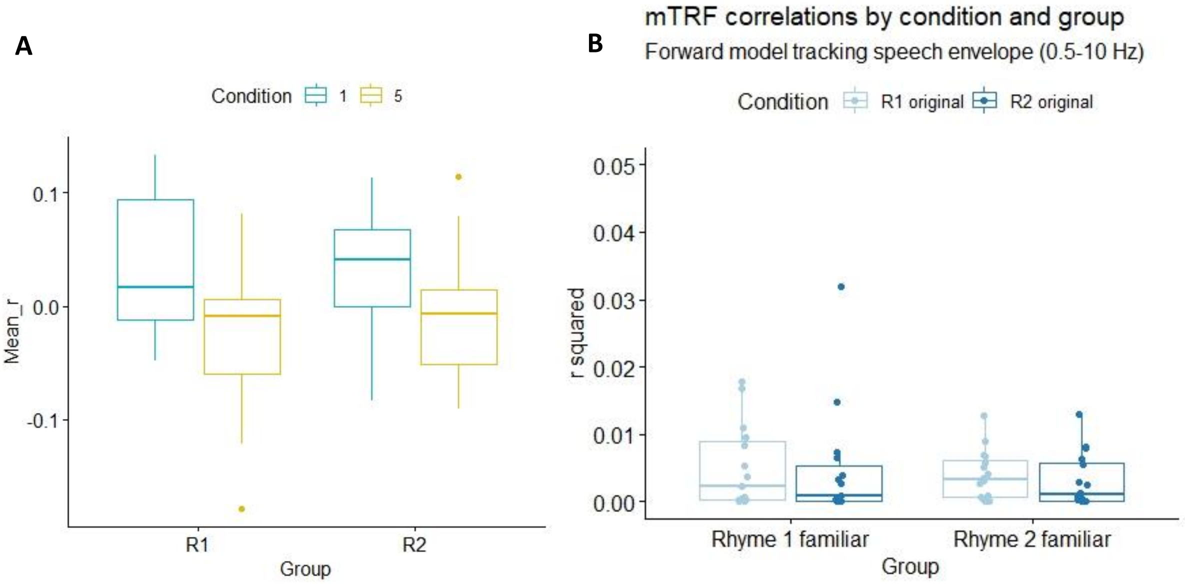

C

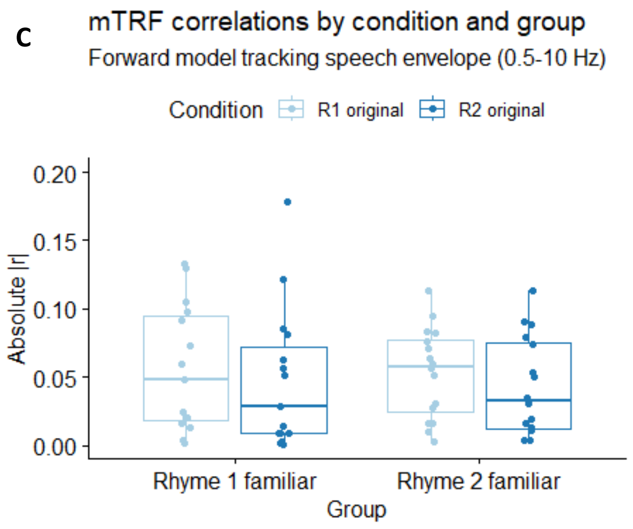

**Figure S14: The correlation coefficients of the mTRF models for speech envelope, 0.5-10Hz**: comparison of the two groups and the two rhymes. Note a higher *r* for rhyme 1 in both groups in panel A. Panel A = with r, Panel B with r squared, Panel C with absolute value of r.

**Table S7: Results of mTRF analysis for the speech envelope: cluster-based permutation analysis**

| **Data** | **Dependent variable** | **Compared conditions** | **0.5 – 2 Hz** | **0.5 – 4 Hz** | **0.5 – 10 Hz** |
|---|---|---|---|---|---|
| All participants n = 31 | mTRF model | R1 vs R2 (rhyme effect) | No signif cluster | 1 neg cluster, p = 0.045 | 1 neg cluster, p = 0.046 |
| | | Familiarity effect | No signif cluster | No signif clusters | No signif clusters |
| n = 23 (Gr1), n = 21 (Gr2) | | Gr 1 vs Gr 2 (group effect) | No signif cluster | No signif clusters | No signif clusters |
| Group 1, n = 15 | mTRF model | Familiarity effect | No signif cluster | 1 neg cluster, p = 0.067 | 1 neg cluster, p = 0.083 |
| Group 2, n = 16 | mTRF model | Familiarity effect | No signif cluster | No signif clusters | No signif clusters |
| All participants n = 38 | mTRF model | Language effect | No signif cluster | 1 neg cluster p = 0.095 | No signif clusters |
| Group 1, n = 20 | mTRF model | Language effect | No signif cluster | No signif clusters | No signif clusters |
| Group 2, n = 18 | mTRF model | Language effect | No signif cluster | No signif clusters | No signif clusters |
| All participants, n = 36 | mTRF model | Rhythm effect | No signif cluster | No signif clusters | No signif clusters |
| Group 1, n = 18 | mTRF model | Rhythm effect | No signif cluster | No signif clusters | No signif clusters |
| Group 2, n = 18 | mTRF model | Rhythm effect | No signif cluster | No signif clusters | No signif clusters |

| | | | | | |
|---|---|---|---|---|---|
| All participants, n = 30 | mTRF model | Phonological effect | No signif cluster | No signif clusters | No signif clusters |
| Group 1, n = 16 | mTRF model | Phonological effect | No signif cluster | No signif clusters | No signif clusters |
| Group 2, n = 14 | mTRF model | Phonological effect | No signif cluster | No signif clusters | No signif clusters |

**Table S8: Results of mTRF analysis for the speech envelope: correlation coefficient *r***

| **Data** | **Dependent variable** | **Compared conditions** | **0.5 – 2 Hz** | **0.5 – 4 Hz** | **0.5 – 10 Hz** |
|---|---|---|---|---|---|
| All participants n = 31 | *r* | R1 vs R2 (rhyme effect) | V = 112, p-value = 0.00664 | V = 110, p-value = 0.005818 | V = 111, p-value = 0.006217 |
| | | Familiarity effect | V = 268, p-value = 0.7063 | V = 271, p-value = 0.6636 | V = 271, p-value = 0.6636 |
| | | Gr 1 vs Gr 2 (group effect) | W = 483, p-value = 0.9721 | W = 468, p-value = 0.8722 | W = 473, p-value = 0.9276 |
| Group 1, n = 15 | *r* | Familiarity effect | V = 99, p-value = 0.02557 | V = 99, p-value = 0.02557 | V = 98, p-value = 0.03015 |
| Group 2, n = 16 | *r* | Familiarity effect | V = 97, p-value = 0.1439 | V = 98, p-value = 0.1297 | V = 97, p-value = 0.1439 |
| All participants n = 38 | *r* | Language effect | V = 439, p-value = 0.328 | V = 441, p-value = 0.3138 | V = 441, p-value = 0.3138 |
| Group 1, n = 20 | *r* | Language effect | V = 143, p-value = 0.165 | V = 142, p-value = 0.1769 | V = 143, p-value = 0.165 |
| Group 2, n = 18 | *r* | Language effect | V = 85, p-value = 1 | V = 88, p-value = 0.9323 | V = 87, p-value = 0.9661 |
| All participants, n = 36 | *r* | Rhythm effect | V = 368, p-value = 0.592 | V = 370, p-value = 0.5707 | V = 366, p-value = 0.6137 |
| Group 1, n = 18 | *r* | Rhythm effect | V = 108, p-value = 0.3465 | V = 108, p-value = 0.3465 | V = 107, p-value = 0.3692 |
| Group 2, n = 18 | *r* | Rhythm effect | V = 84, p-value = 0.9661 | V = 84, p-value = 0.9661 | V = 84, p-value = 0.9661 |
| All participants, n = 30 | *r* | Phonological effect | V = 249, p-value = 0.7457 | V = 239, p-value = 0.9032 | V = 239, p-value = 0.9032 |
| Group 1, n = 16 | *r* | Phonological effect | V = 88, p-value = 0.3225 | V = 82, p-value = 0.4954 | V = 82, p-value = 0.4954 |

| | | | | | |
|---|---|---|---|---|---|
| Group 2, n = 14 | *r* | Phonological effect | V = 48, p-value = 0.8077 | V = 47, p-value = 0.7609 | V = 48, p-value = 0.8077 |

**Table S9: Results of mTRF analysis for the speech envelope: squared correlation coefficient $r^2$**

| **Data** | **Dependent variable** | **Compared conditions** | **0.5 – 2 Hz** | **0.5 – 4 Hz** | **0.5 – 10 Hz** |
|---|---|---|---|---|---|
| All participants n = 31 | $r^2$ | R1 vs R2 (rhyme effect) | V = 207, p-value = 0.4327 | V = 207, p-value = 0.4327 | V = 209, p-value = 0.4559 |
| | | Familiarity effect | V = 254, p-value = 0.9154 | V = 248, p-value = 1 | V = 241, p-value = 0.9001 |
| | | Gr 1 vs Gr 2 (group effect) | W = 485, p-value = 0.9498 | W = 467, p-value = 0.8612 | W = 463, p-value = 0.8175 |
| Group 1, n = 15 | $r^2$ | Familiarity effect | V = 69, p-value = 0.6387 | V = 69, p-value = 0.6387 | V = 67, p-value = 0.7197 |
| Group 2, n = 16 | $r^2$ | Familiarity effect | V = 78, p-value = 0.6322 | V = 80, p-value = 0.5619 | V = 81, p-value = 0.5282 |
| All participants n = 38 | $r^2$ | Language effect | V = 462, p-value = 0.1892 | V = 478, p-value = 0.1216 | V = 474, p-value = 0.1365 |
| Group 1, n = 20 | $r^2$ | Language effect | V = 150, p-value = 0.09731 | V = 155, p-value = 0.06372 | V = 153, p-value = 0.07585 |
| Group 2, n = 18 | $r^2$ | Language effect | V = 83, p-value = 0.9323 | V = 90, p-value = 0.865 | V = 90, p-value = 0.865 |
| All participants, n = 36 | $r^2$ | Rhythm effect | V = 399, p-value = 0.3074 | V = 87, p-value = 0.9661 | V = 398, p-value = 0.3149 |
| Group 1, n = 18 | $r^2$ | Rhythm effect | V = 121, p-value = 0.1297 | V = 121, p-value = 0.1297 | V = 120, p-value = 0.1415 |
| Group 2, n = 18 | $r^2$ | Rhythm effect | V = 82, p-value = 0.8986 | V = 87, p-value = 0.9661 | V = 86, p-value = 1 |
| All participants, n = 30 | $r^2$ | Phonological effect | V = 300, p-value = 0.1706 | V = 309, p-value = 0.1191 | V = 299, p-value = 0.1772 |
| Group 1, n = 16 | $r^2$ | Phonological effect | V = 97, p-value = 0.1439 | V = 101, p-value = 0.09344 | V = 99, p-value = 0.1167 |
| Group 2, n = 14 | $r^2$ | Phonological effect | V = 56, p-value = 0.8552 | V = 60, p-value = 0.6698 | V = 58, p-value = 0.7609 |

**Table S10: Results of mTRF analysis for the speech envelope: squared correlation coefficient |r|**

| Data | Dependent variable | Compared conditions | 0.5 – 2 Hz | 0.5 – 4 Hz | 0.5 – 10 Hz |
|---|---|---|---|---|---|
| All participants n = 31 | \|r\| | R1 vs R2 (rhyme effect) | V = 187, p-value = 0.2395 | V = 195, p-value = 0.3082 | V = 198, p-value = 0.3369 |
| | | Familiarity effect | V = 248, p-value = 1 | V = 244, p-value = 0.9461 | V = 239, p-value = 0.8696 |
| | | Gr 1 vs Gr 2 (group effect) | W = 485, p-value = 0.9498 | W = 467, p-value = 0.8612 | W = 463, p-value = 0.8175 |
| Group 1, n = 15 | \|r\| | Familiarity effect | V = 73, p-value = 0.4887 | V = 70, p-value = 0.5995 | V = 69, p-value = 0.6387 |
| Group 2, n = 16 | \|r\| | Familiarity effect | V = 83, p-value = 0.4637 | V = 85, p-value = 0.4037 | V = 86, p-value = 0.3755 |
| All participants n = 38 | \|r\| | Language effect | V = 466, p-value = 0.1702 | V = 476, p-value = 0.1289 | V = 470, p-value = 0.1526 |
| Group 1, n = 20 | \|r\| | Language effect | V = 156, p-value = 0.05826 | V = 158, p-value = 0.04844 | V = 157, p-value = 0.05317 |
| Group 2, n = 18 | \|r\| | Language effect | V = 81, p-value = 0.865 | V = 86, p-value = 1 | V = 84, p-value = 0.9661 |
| All participants, n = 36 | \|r\| | Rhythm effect | V = 400, p-value = 0.3 | V = 394, p-value = 0.3461 | V = 397, p-value = 0.3226 |
| Group 1, n = 18 | \|r\| | Rhythm effect | V = 126, p-value = 0.08143 | V = 119, p-value = 0.154 | V = 121, p-value = 0.1297 |
| Group 2, n = 18 | \|r\| | Rhythm effect | V = 79, p-value = 0.7987 | V = 83, p-value = 0.9323 | V = 83, p-value = 0.9323 |
| All participants, n = 30 | \|r\| | Phonological effect | V = 285, p-value = 0.2894 | V = 291, p-value = 0.2367 | V = 285, p-value = 0.2894 |
| Group 1, n = 16 | \|r\| | Phonological effect | V = 93, p-value = 0.2114 | V = 95, p-value = 0.1754 | V = 93, p-value = 0.2114 |
| Group 2, n = 14 | \|r\| | Phonological effect | V = 53, p-value = 1 | V = 56, p-value = 0.8552 | V = 54, p-value = 0.9515 |

#### 2.3.2. mTRF models between EEG and the triggers for stressed syllables

##### *2.3.2.1. Bandpass filtered over 0.5 – 2 Hz*

For the mTRF computed on data filtered between 0.5 and 2 Hz, there were two significant clusters, one for a rhyme effect, with a positive middle-right frontal and central cluster from 100 to 450ms, indicating more activation for the first rhyme than the second (Fig. S15), and one for a language effect in group 2, with a positive cluster bilateral and middle frontal from 550 to 650ms, indicating more activation for the original rhyme than the language-manipulated one (Fig. S16).

The analysis of the r coefficients of the mTRFs (the correlation between the predicted and real signal), the Wilcoxon signed rank tests showed a trend for a phonological effect in Group 1, p = 0.051, with higher r values for the original rhyme (mean r = 0.01) than for the phonological manipulation (mean r = -0.017).

When analysing the r squared values, there was just a trend for a rhythm effect in group 2, p = 0.081, with higher r squared values for the original rhyme (r squared  = 0.0015) than for rhythm manipulated rhyme (r squared = 0.00053).

The analysis of the absolute values of r showed no significant or trending results.

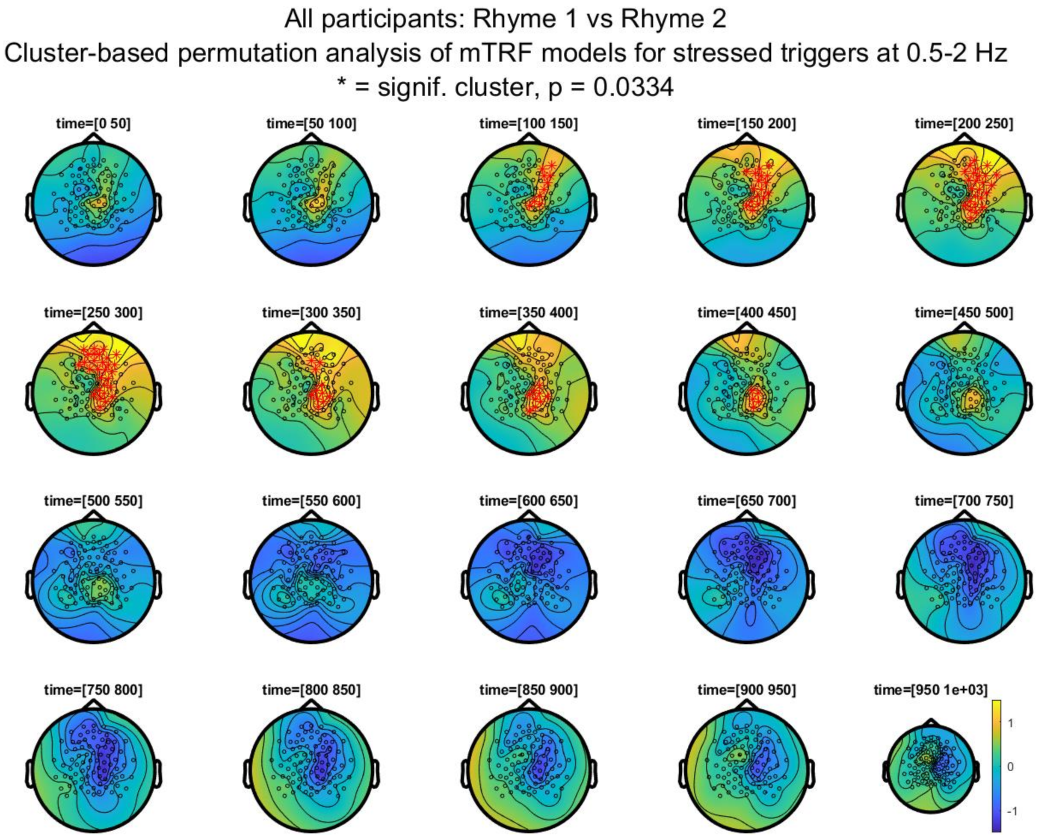

**Figure S15: Cluster based permutation analysis of the mTRF models for stressed syllables, 0.5-2Hz**: comparison of the two rhymes. Note a fronto-central early positive cluster with more activation for Rhyme 1 (*Butzemann*).

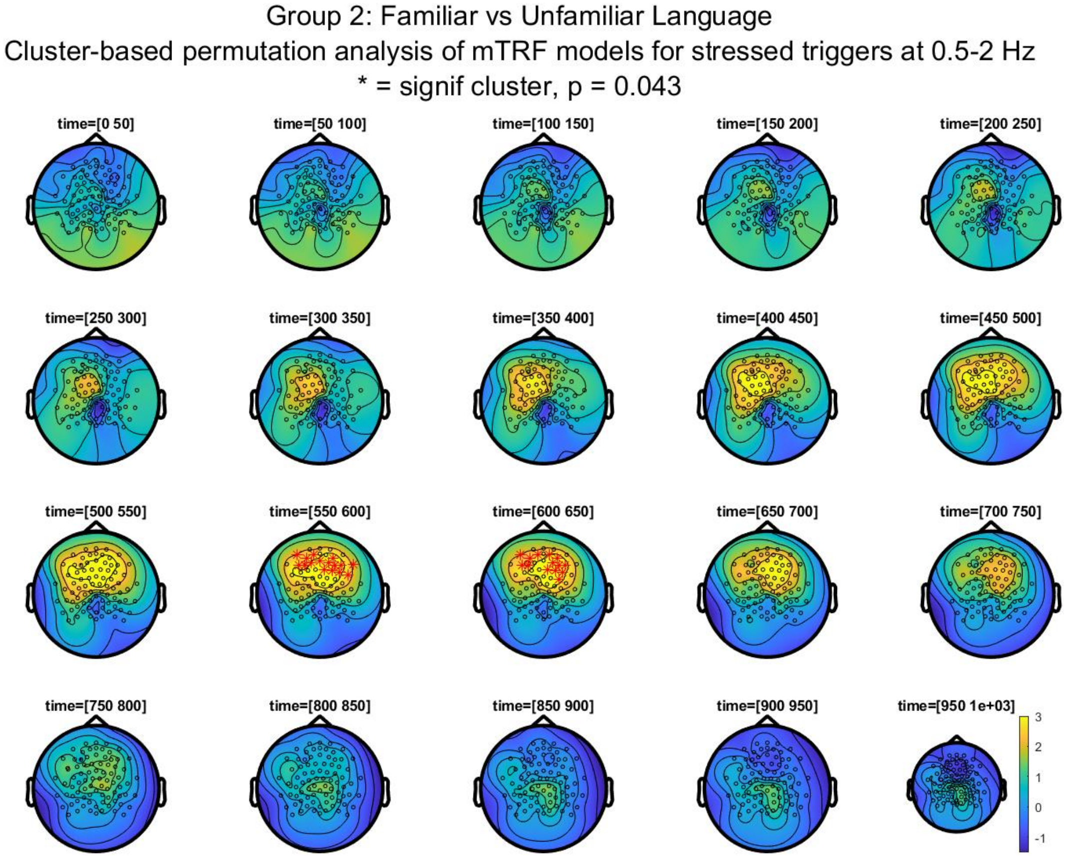

**Figure S16: Cluster based permutation analysis of the mTRF models for stressed syllables, 0.5-2Hz**: comparison of the familiar language and the backward speech (here noted as unfamiliar language). Note a fronta positive cluster with more activation for the familiar language than for the unfamiliar, unintelligible one (backward speech).

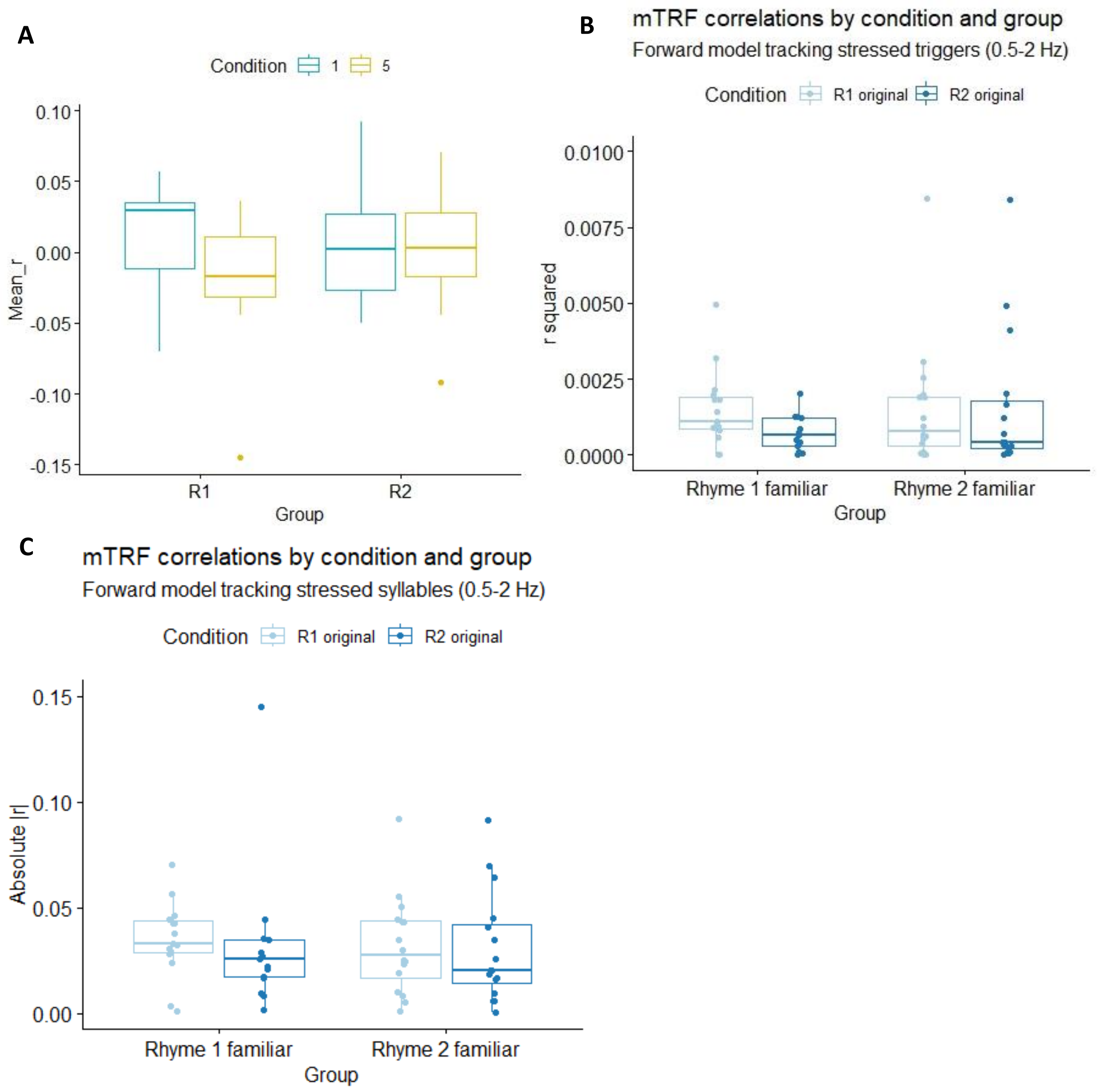

**Figure S17: Correlation coefficient of the mTRF models for stressed syllables, 0.5-2Hz**: comparison condition 1 (*Butzemann*) and condition 5 (*Es war eine Mutter*) in Group 1 (familiar rhyme R1 = condition 1) and Group 2 (familiar rhyme R2 = condition 5). Panel A = with r, Panel B with r squared, Panel C with absolute value of r.

###### *2.3.2.2. Bandpass filtered over 0.5 – 4 Hz*

For the mTRF computed on data filtered between 0.5 and 4 Hz, there were two trending clusters, one for a rhyme effect, with a positive middle-frontal and central cluster, indicating higher activity for the first rhyme than the second (Fig. S18), and one for a language effect in group 2 with a right frontal positive cluster (indicating higher activity in the original rhyme than in the language manipulated rhyme) (Fig. S19).

When analysing the r correlation values between the predicted and real signal (using Spearman correlation), the Wilcoxon signed rank tests showed a significant phonologic effect in group 1, with the original rhyme having higher r values (mean r = 0.011) than the phonological manipulated one (mean r = -0.014).

The analysis of r squared values showed a trend for a rhythm effect in group 2, towards higher r squared values in the original rhyme (r sqrd = 0.0016) than in the rhythm manipulated rhyme (r sqrd = 0.00037).

The analysis of the absolute value or r revealed no effects and no trends.

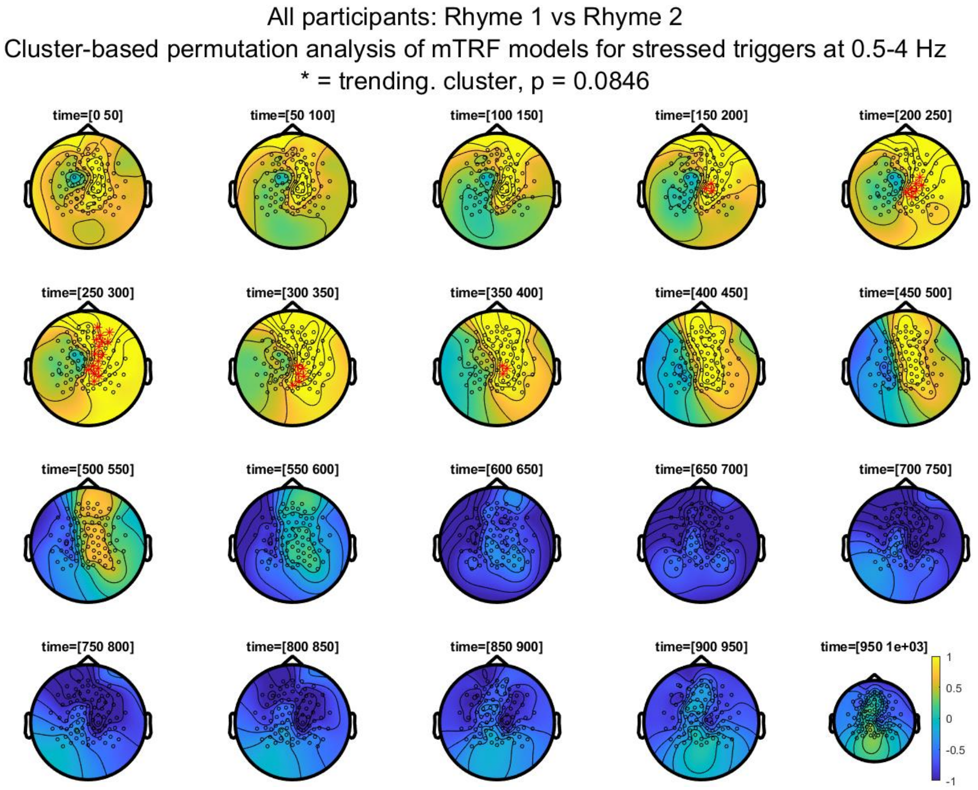

**Figure S18: Cluster based permutation analysis of the mTRF models for stressed syllables, 0.5-4Hz**: comparison of the two rhymes. Note a fronto-central early positive cluster with more activation for Rhyme 1 (*Butzemann*).

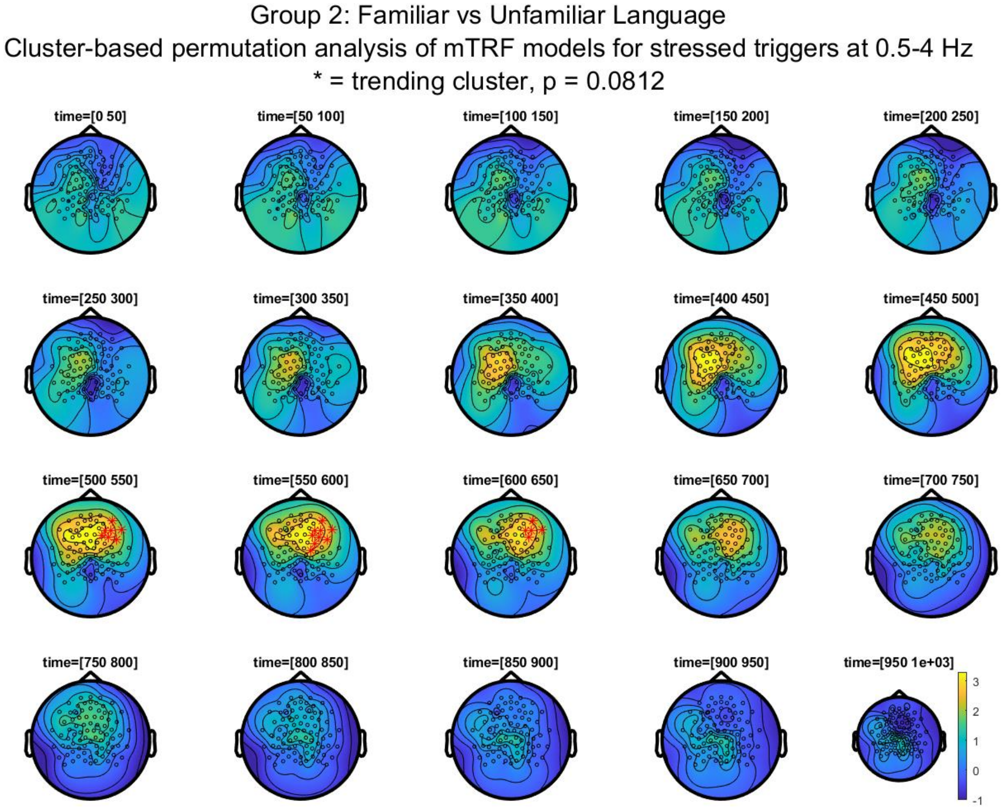

**Figure S19: Cluster based permutation analysis of the mTRF models for stressed syllables, 0.5-4Hz, Group 2**: comparison of the language conditions (familiar language vs. backward speech). Note a right-frontal positive cluster with more activation for the familiar language.

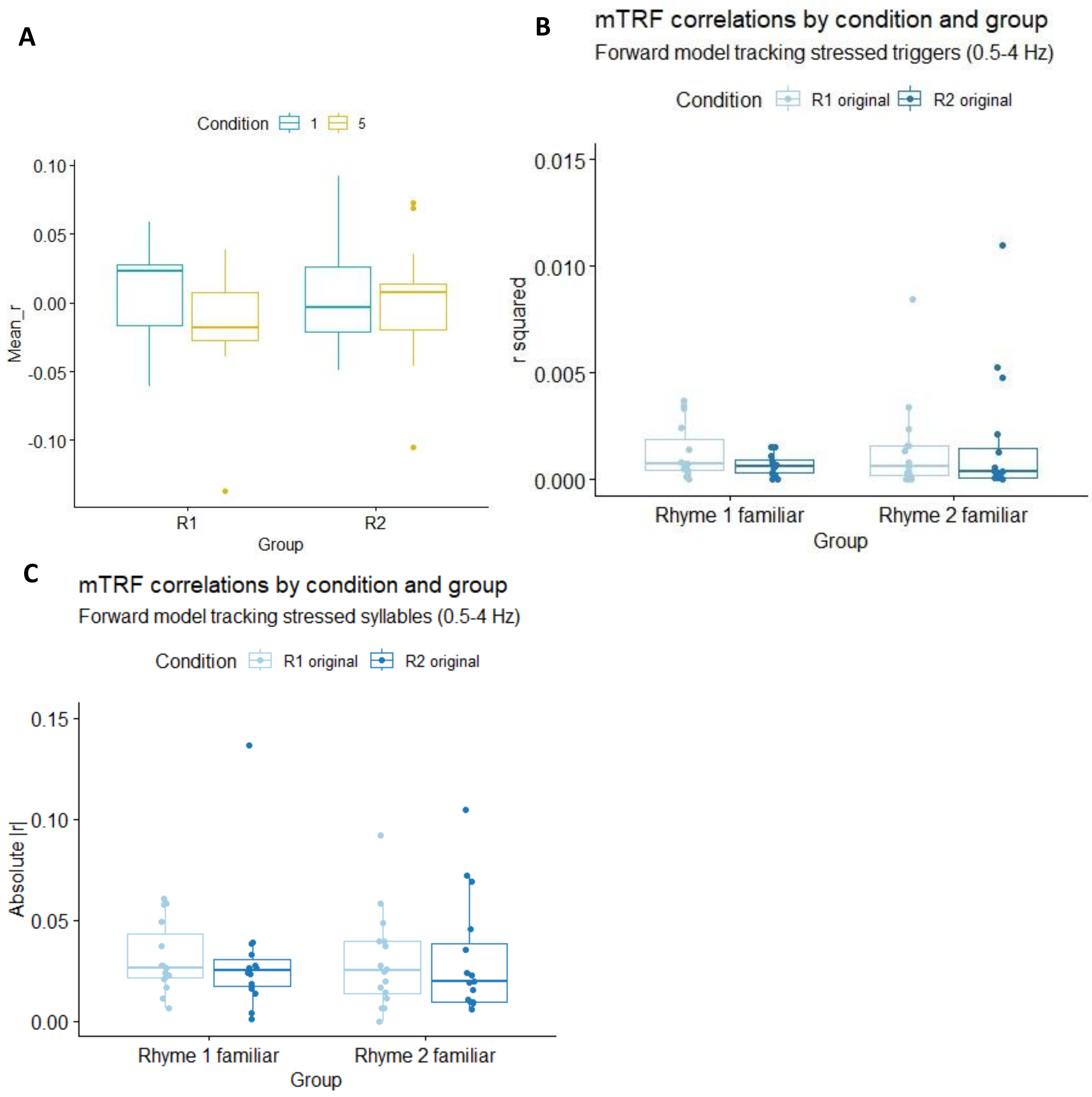

**Figure S20: Correlation coefficient of the mTRF models for stressed syllables, 0.5-4Hz**: comparison condition 1 (*Butzemann*) and condition 5 (*Es war eine Mutter*) in Group 1 (familiar rhyme R1 = condition 1) and Group 2 (familiar rhyme R2 = condition 5). Panel A = with r, Panel B with r squared, Panel C with absolute value of r.

###### *2.3.2.3. Bandpass filtered over 0.5 – 10 Hz*

For the mTRF models computed on data bandpass filtered over 0.5-10 Hz, there was a rhyme effect with a significant cluster bilateral frontal, between 600 and 850ms, indicating higher activity in the second rhyme than the first (dark blue = negative values) (Fig. S21). There was also a language effect in group 2, with a significant cluster in the frontal (mostly right) region between 500 and 650ms, indicating higher activity for the familiar language (unmanipulated rhyme) (Fig. S22).

The analysis of the r correlation values between the predicted and the real signal (using Spearman correlation) was performed with Wilcoxon signed rank tests and showed no significant effects, just a trend for the phonetic paradigm in group 1, with the original rhyme having higher r values (mean r = 0.0086) than the phonetic manipulation (mean r = - 0.012).

When analysing r squared, there was just a trend for a rhythm effect in group 2, towards higher r squared values for the original rhyme (r sqrd = 0.0014) than for the rhythm manipulated rhyme (r sqrd = 0.00032).

When analysing the absolute value of r (|r|) there were no effects at all.

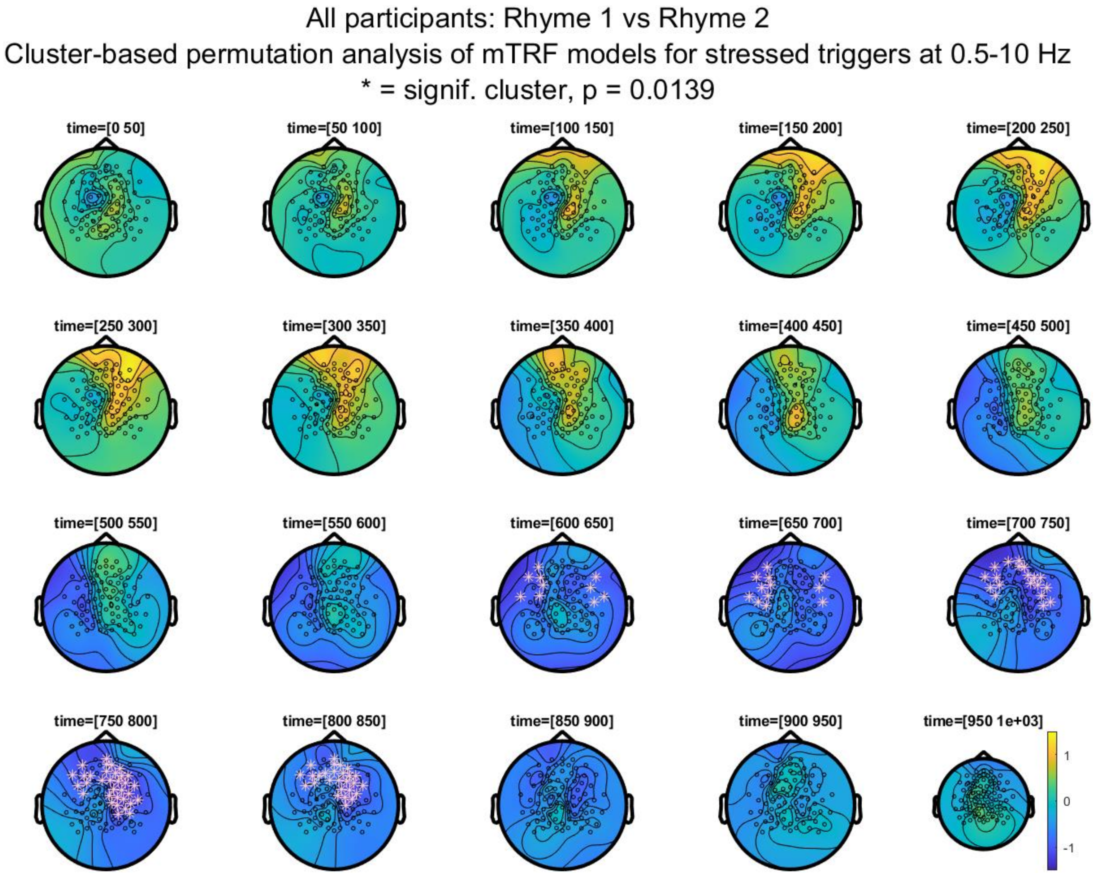

**Figure S21: Cluster based permutation analysis of the mTRF models for stressed syllables, 0.5-10Hz**: comparison of the two rhymes. Note a fronto-central late negative cluster with more activation for Rhyme 2 (*Es war eine Mutter*).

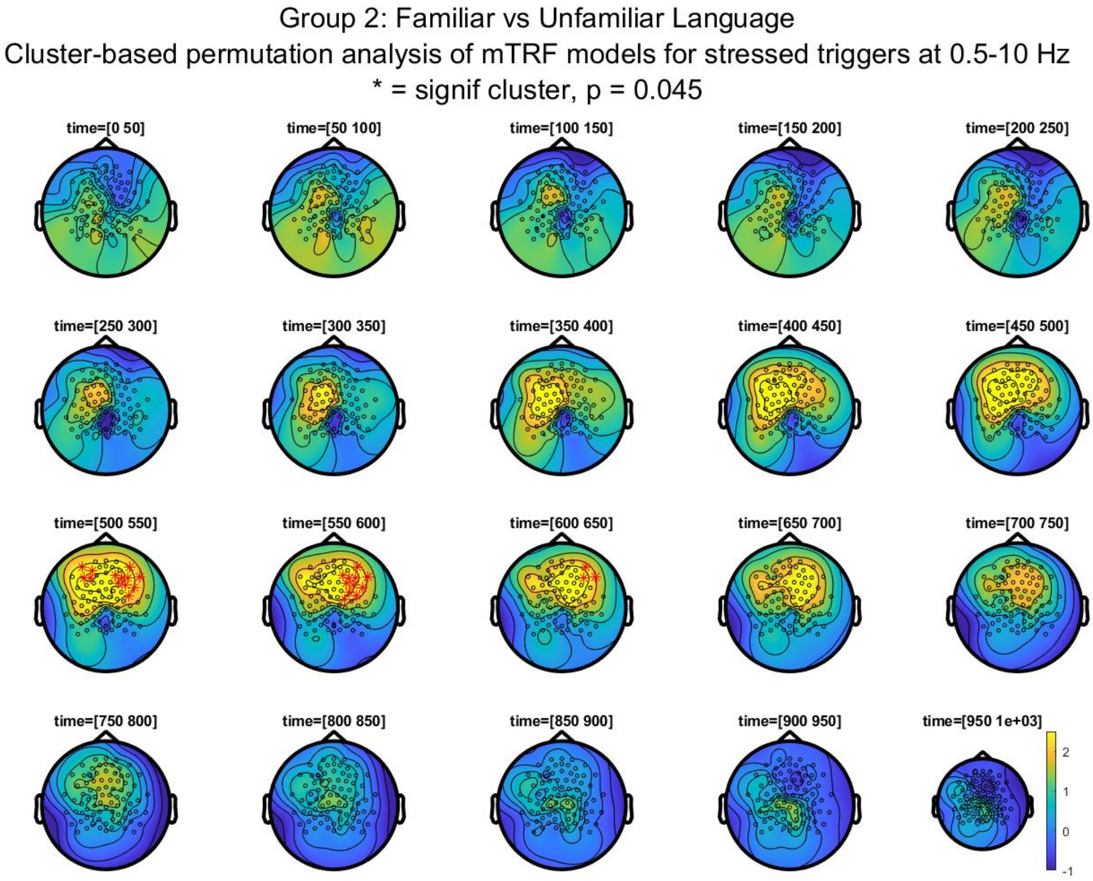

**Figure S22: Cluster based permutation analysis of the mTRF models for stressed syllables, 0.5-10Hz**: comparison of the two language conditions (familiar vs. backward speech). Note a fronto-central positive cluster with more activation for the familiar language.

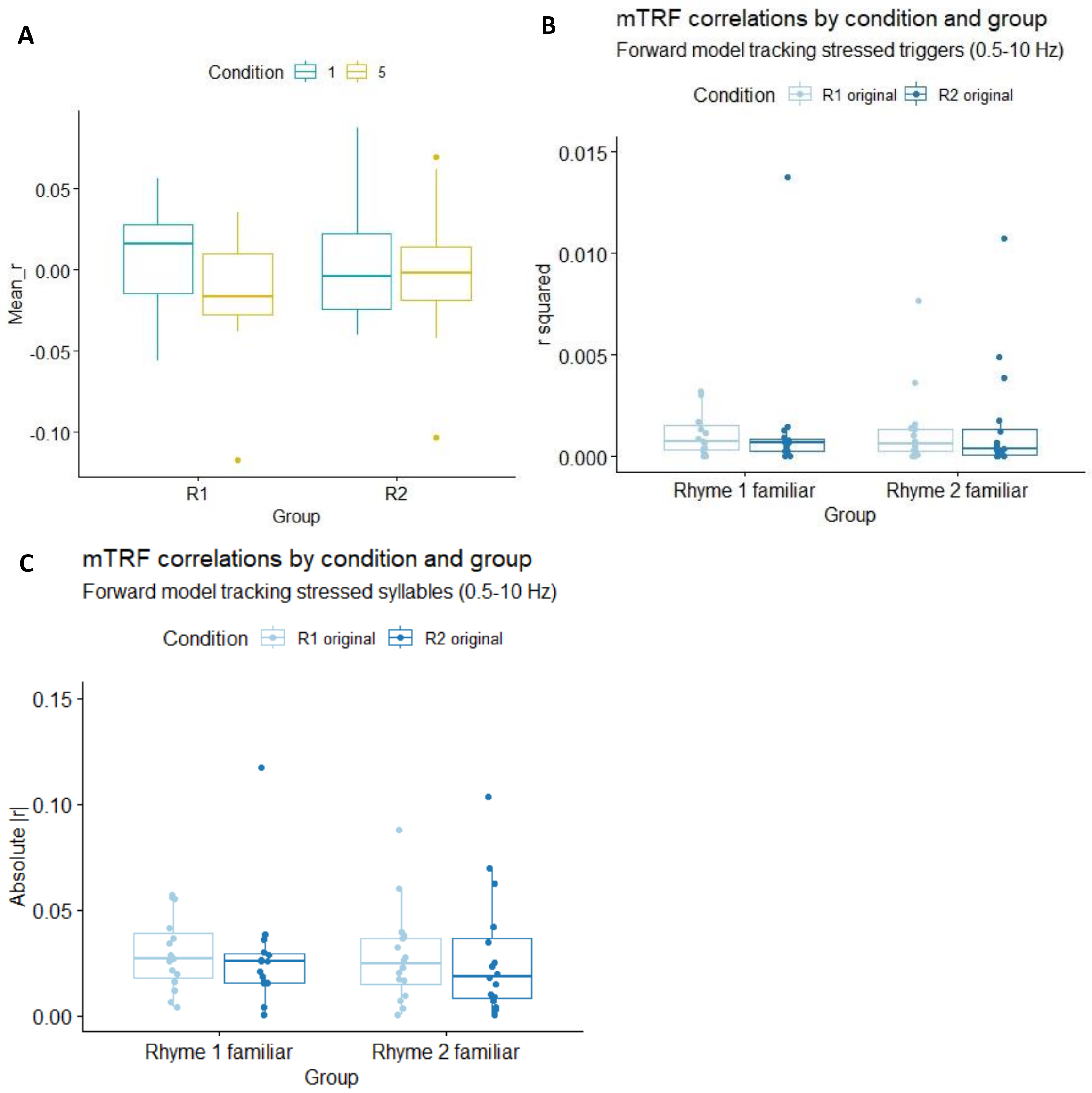

**Fig. S23: Correlation coefficient of the mTRF models for stressed syllables, 0.5-10Hz**: comparison condition 1 (*Butzemann*) and condition 5 (*Es war eine Mutter*) in Group 1 (familiar rhyme R1 = condition 1) and Group 2 (familiar rhyme R2 = condition 5). Panel A = with r, Panel B with r squared, Panel C with absolute value of r.

**Table S11: Results of mTRF analysis for the stressed syllables: cluster-based analysis of the mTRF models**

| **Data** | **Dependent variable** | **Compared conditions** | **0.5 – 2 Hz** | **0.5 – 4 Hz** | **0.5 – 10 Hz** |
|---|---|---|---|---|---|
| All participants n = 31 | mTRF model | R1 vs R2 (rhyme effect) | 1 pos cluster p = 0.0334 | 1 pos cluster p = 0.0846 | 1 neg cluster p = 0.0139 |
| | | Familiarity effect | No signif clusters | No signif clusters | No signif clusters |
| n = 23 (Gr1), n = 21 (Gr2) | | Gr 1 vs Gr 2 (group effect) | No signif clusters | No signif clusters | No signif clusters |
| Group 1, n = 15 | mTRF model | Familiarity effect | No signif clusters | No signif clusters | No signif clusters |

| | | | | | |
|---|---|---|---|---|---|
| Group 2, n = 16 | mTRF model | Familiarity effect | No signif clusters | No signif clusters | No signif clusters |
| All participants n = 38 | mTRF model | Language effect | No signif clusters | No signif clusters | No signif clusters |
| Group 1, n = 20 | mTRF model | Language effect | No signif clusters | No signif clusters | No signif clusters |
| Group 2, n = 18 | mTRF model | Language effect | 1 pos cluster p = 0.0434 | 1 pos cluster p = 0.0812 | 1 pos cluster p = 0.0450 |
| All participants, n = 36 | mTRF model | Rhythm effect | No signif clusters | No signif clusters | No signif clusters |
| Group 1, n = 18 | mTRF model | Rhythm effect | No signif clusters | No signif clusters | No signif clusters |
| Group 2, n = 18 | mTRF model | Rhythm effect | No signif clusters | No signif clusters | No signif clusters |
| All participants, n = 30 | mTRF model | Phonological effect | No signif clusters | No signif clusters | No signif clusters |
| Group 1, n = 16 | mTRF model | Phonological effect | No signif clusters | No signif clusters | No signif clusters |
| Group 2, n = 14 | mTRF model | Phonological effect | No signif clusters | No signif clusters | No signif clusters |

**Table S12: Results of mTRF analysis for the stressed triggers: correlation coefficient *r***

| **Data** | **Dependent variable** | **Compared conditions** | **0.5 – 2 Hz** | **0.5 – 4 Hz** | **0.5 – 10 Hz** |
|---|---|---|---|---|---|
| All participants n = 31 | *r* | R1 vs R2 (rhyme effect) | V = 191, p-value = 0.2724 | V = 181, p-value = 0.1954 | V = 182, p-value = 0.2023 |
| | | Familiarity effect | V = 299, p-value = 0.3271 | V = 282, p-value = 0.5168 | V = 277, p-value = 0.5814 |
| | | Gr 1 vs Gr 2 (group effect) | W = 464, p-value = 0.8284 | W = 455, p-value = 0.7318 | W = 461, p-value = 0.7958 |
| Group 1, n = 15 | *r* | Familiarity effect | V = 85, p-value = 0.1688 | V = 83, p-value = 0.2078 | V = 82, p-value = 0.2293 |
| Group 2, n = 16 | *r* | Familiarity effect | V = 69, p-value = 0.9799 | V = 79, p-value = 0.5966 | V = 79, p-value = 0.5966 |
| All participants n = 38 | *r* | Language effect | V = 424, p-value = 0.4465 | V = 407, p-value = 0.6055 | V = 366, p-value = 0.9543 |
| Group 1, n = 20 | *r* | Language effect | V = 135, p-value = 0.2774 | V = 136, p-value = 0.2611 | V = 125, p-value = 0.4749 |
| Group 2, n = 18 | *r* | Language effect | V = 83, p-value = 0.9323 | V = 73, p-value = 0.6095 | V = 66, p-value = 0.4171 |

| | | | | | |
|---|---|---|---|---|---|
| All participants, n = 36 | *r* | Rhythm effect | V = 399, p-value = 0.3074 | V = 377, p-value = 0.4989 | V = 356, p-value = 0.7269 |
| Group 1, n = 18 | *r* | Rhythm effect | V = 117, p-value = 0.1815 | V = 113, p-value = 0.2462 | V = 108, p-value = 0.3465 |
| Group 2, n = 18 | *r* | Rhythm effect | V = 88, p-value = 0.9323 | V = 82, p-value = 0.8986 | V = 75, p-value = 0.6705 |
| All participants, n = 30 | *r* | Phonological effect | V = 292, p-value = 0.2286 | V = 298, p-value = 0.184 | V = 283, p-value = 0.3085 |
| Group 1, n = 16 | *r* | Phonological effect | V = 106, p-value = 0.05066 | V = 107, p-value = 0.04431 | V = 104, p-value = 0.0654 |
| Group 2, n = 14 | *r* | Phonological effect | V = 47, p-value = 0.7609 | V = 54, p-value = 0.9515 | V = 50, p-value = 0.9032 |

**Table S13: Results of mTRF analysis for the stressed triggers: squared correlation coefficient $r^2$**

| Data | Dependent variable | Compared conditions | 0.5 – 2 Hz | 0.5 – 4 Hz | 0.5 – 10 Hz |
|---|---|---|---|---|---|
| All participants n = 31 | $r^2$ | R1 vs R2 (rhyme effect) | V = 174, p-value = 0.1517 | V = 215, p-value = 0.5294 | V = 218, p-value = 0.5682 |
| | | Familiarity effect | V = 276, p-value = 0.5948 | V = 255, p-value = 0.9001 | V = 256, p-value = 0.8848 |
| | | Gr 1 vs Gr 2 (group effect) | W = 518, p-value = 0.5998 | W = 534, p-value = 0.4538 | W = 522, p-value = 0.5614 |
| Group 1, n = 15 | $r^2$ | Familiarity effect | V = 87, p-value = 0.1354 | V = 72, p-value = 0.5245 | V = 71, p-value = 0.5614 |
| Group 2, n = 16 | $r^2$ | Familiarity effect | V = 82, p-value = 0.4954 | V = 74, p-value = 0.782 | V = 75, p-value = 0.7436 |
| All participants n = 38 | $r^2$ | Language effect | V = 417, p-value = 0.509 | V = 402, p-value = 0.6566 | V = 408, p-value = 0.5955 |
| Group 1, n = 20 | $r^2$ | Language effect | V = 120, p-value = 0.5958 | V = 119, p-value = 0.6215 | V = 118, p-value = 0.6477 |
| Group 2, n = 18 | $r^2$ | Language effect | V = 95, p-value = 0.7019 | V = 91, p-value = 0.8317 | V = 93, p-value = 0.766 |
| All participants, n = 36 | $r^2$ | Rhythm effect | V = 392, p-value = 0.3624 | V = 378, p-value = 0.4891 | V = 388, p-value = 0.3964 |

| | | | | | |
|---|---|---|---|---|---|
| Group 1, n = 18 | $r^2$ | Rhythm effect | V = 78, p-value = 0.766 | V = 70, p-value = 0.5226 | V = 75, p-value = 0.6705 |
| Group 2, n = 18 | $r^2$ | Rhythm effect | V = 126, p-value = 0.08143 | V = 128, p-value = 0.06654 | V = 127, p-value = 0.07368 |
| All participants, n = 30 | $r^2$ | Phonological effect | V = 169, p-value = 0.1981 | V = 177, p-value = 0.2621 | V = 175, p-value = 0.2449 |
| Group 1, n = 16 | $r^2$ | Phonological effect | V = 47, p-value = 0.2979 | V = 48, p-value = 0.3225 | V = 48, p-value = 0.3225 |
| Group 2, n = 14 | $r^2$ | Phonological effect | V = 43, p-value = 0.583 | V = 47, p-value = 0.7609 | V = 44, p-value = 0.6257 |

**Table S14: Results of mTRF analysis for the stressed triggers: absolute correlation coefficient *|r|***

| **Data** | **Dependent variable** | **Compared conditions** | **0.5 – 2 Hz** | **0.5 – 4 Hz** | **0.5 – 10 Hz** |
|---|---|---|---|---|---|
| All participants n = 31 | $\|r\|$ | R1 vs R2 (rhyme effect) | V = 188, p-value = 0.2474 | V = 222, p-value = 0.6219 | V = 223, p-value = 0.6357 |
| | | Familiarity effect | V = 269, p-value = 0.6919 | V = 262, p-value = 0.7942 | V = 254, p-value = 0.9154 |
| | | Gr 1 vs Gr 2 (group effect) | W = 518, p-value = 0.5998 | W = 534, p-value = 0.4538 | W = 522, p-value = 0.5614 |
| Group 1, n = 15 | $\|r\|$ | Familiarity effect | V = 80, p-value = 0.2769 | V = 72, p-value = 0.5245 | V = 67, p-value = 0.7197 |
| Group 2, n = 16 | $\|r\|$ | Familiarity effect | V = 80, p-value = 0.5619 | V = 71, p-value = 0.8999 | V = 74, p-value = 0.782 |
| All participants n = 38 | $\|r\|$ | Language effect | V = 405, p-value = 0.6257 | V = 409, p-value = 0.5856 | V = 419, p-value = 0.4907 |
| Group 1, n = 20 | $\|r\|$ | Language effect | V = 110, p-value = 0.8695 | V = 117, p-value = 0.6742 | V = 118, p-value = 0.6477 |
| Group 2, n = 18 | $\|r\|$ | Language effect | V = 95, p-value = 0.7019 | V = 96, p-value = 0.6705 | V = 99, p-value = 0.5798 |
| All participants, n = 36 | $\|r\|$ | Rhythm effect | V = 391, p-value = 0.3707 | V = 376, p-value = 0.5089 | V = 382, p-value = 0.4507 |
| Group 1, n = 18 | $\|r\|$ | Rhythm effect | V = 81, p-value = 0.865 | V = 79, p-value = 0.7987 | V = 81, p-value = 0.865 |

| | | | | | |
|---|---|---|---|---|---|
| Group 2, n = 18 | /r/ | Rhythm effect | V = 121, p-value = 0.1297 | V = 122, p-value = 0.1187 | V = 118, p-value = 0.1674 |
| All participants, n = 30 | /r/ | Phonological effect | V = 179, p-value = 0.2801 | V = 178, p-value = 0.271 | V = 179, p-value = 0.2801 |
| Group 1, n = 16 | /r/ | Phonological effect | V = 48, p-value = 0.3225 | V = 49, p-value = 0.3484 | V = 51, p-value = 0.4037 |
| Group 2, n = 14 | /r/ | Phonological effect | V = 44, p-value = 0.6257 | V = 47, p-value = 0.7609 | V = 44, p-value = 0.6257 |

#### 2.4. Mutual information analysis

##### 2.4.1. Amp-amp

Due to two extreme outliers in condition 5 (original rhyme 2), we first removed the outliers 3SD above the mean when comparing these two conditions (rhyme effect, familiarity effect and group effect).

The analysis of MI values averaged over 0.5 – 2 Hz, 0.5-4Hz and 0.5-10 Hz showed a significant effect for rhyme familiarity over all participants, mostly driven by Group 1, with higher MI for the unfamiliar rhyme than for the familiar rhyme (Fig. S24).

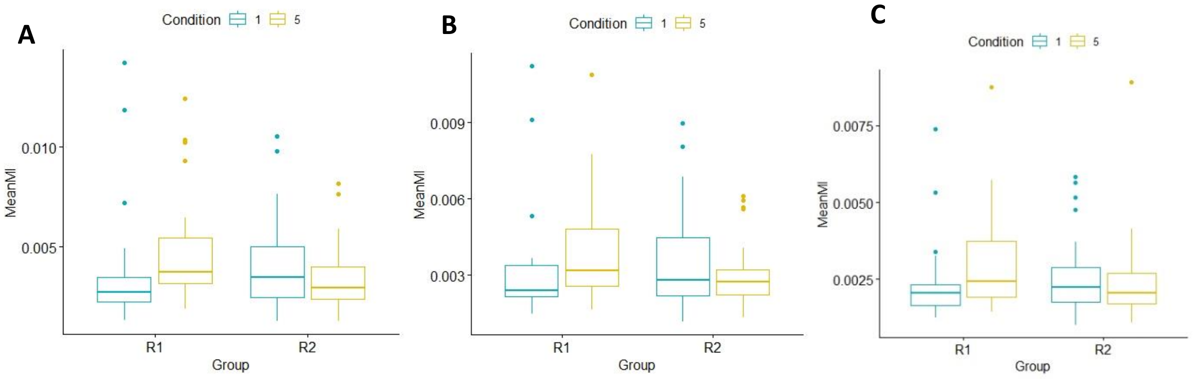

**Fig S24: Mutual Information Amp-Amp**, averaged over 0.5-2Hz (A), 0.5-4Hz (B), 0.5-10Hz (C).

**Table S15: Results of MI amplitude-amplitude analysis**

| Data | Dependent variable | Compared conditions | 0.5 – 2 Hz | 0.5 – 4 Hz | 0.5 – 10 Hz |
|---|---|---|---|---|---|
| All participants n = 55 | MI amp-amp | R1 vs R2 (rhyme effect) | V = 638, p-value = 0.2706 | V = 659, p-value = 0.3545 | V = 600, p-value = 0.1556 |
| | | Familiarity effect | V = 1112, p-value = 0.004219 | V = 1046, p-value = 0.02098 | V = 1059, p-value = 0.01564 |
| | | Gr 1 vs Gr 2 (group effect) | W = 1538, p-value = 0.8598 | W = 1542, p-value = 0.841 | W = 1577, p-value = 0.6817 |

| | | | | | |
|---|---|---|---|---|---|
| Group 1, n = 27 | MI amp-amp | Familiarity effect | V = 60, p-value = 0.001253 | V = 76, p-value = 0.005465 | V = 51, p-value = 0.0004787 |
| Group 2, n = 30 | MI amp-amp | Familiarity effect | V = 274, p-value = 0.4045 | V = 262, p-value = 0.5561 | V = 243, p-value = 0.8394 |
| All participants n = 58 | MI amp-amp | Language effect | V = 797, p-value = 0.6534 | V = 852, p-value = 0.9815 | V = 849, p-value = 0.9629 |
| Group 1, n = 27 | MI amp-amp | Language effect | V = 171, p-value = 0.679 | V = 200, p-value = 0.804 | V = 171, p-value = 0.679 |
| Group 2, n = 31 | MI amp-amp | Language effect | V = 236, p-value = 0.8242 | V = 247, p-value = 0.9923 | V = 263, p-value = 0.7793 |
| All participants, n = 56 | MI amp-amp | Rhythm effect | V = 739, p-value = 0.6332 | V = 684, p-value = 0.3545 | V = 645, p-value = 0.2135 |
| Group 1, n = 26 | MI amp-amp | Rhythm effect | V = 155, p-value = 0.6171 | V = 154, p-value = 0.5995 | V = 169, p-value = 0.8809 |
| Group 2, n = 30 | MI amp-amp | Rhythm effect | V = 220, p-value = 0.8078 | V = 194, p-value = 0.44 | V = 175, p-value = 0.2449 |
| All participants, n = 30 | MI amp-amp | Phonological effect | V = 699, p-value = 0.5547 | V = 665, p-value = 0.3813 | V = 667, p-value = 0.3905 |
| Group 1, n = 25 | MI amp-amp | Phonological effect | V = 154, p-value = 0.8325 | V = 143, p-value = 0.615 | V = 123, p-value = 0.2996 |
| Group 2, n = 14 | MI amp-amp | Phonological effect | V = 200, p-value = 0.5158 | V = 200, p-value = 0.5158 | V = 218, p-value = 0.7766 |

##### 2.4.2. Phase-amp

Due to two extreme outliers in condition 5 (original rhyme 2), we first removed the outliers 3SD above the mean when comparing these two conditions (rhyme effect, familiarity effect and group effect). The analysis of MI values averaged over 0.5 – 2 Hz, 0.5-4Hz and 0.5-10 Hz showed no effects and no trends.

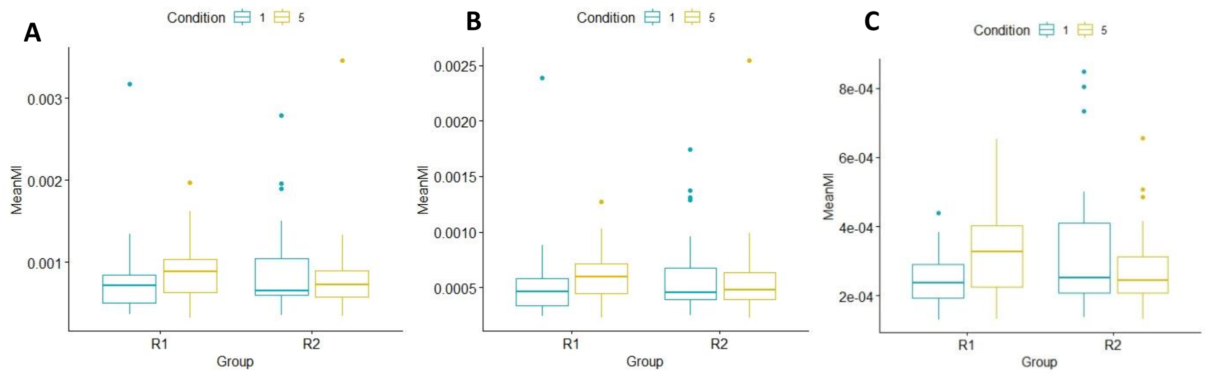

**Fig. S25: Mutual information phase-amplitude** averaged over 0.5-2Hz (A), 0.5-4Hz (B) and 0.5-10Hz (C).

**Table S16: Results of MI phase-amplitude analysis**

| **Data** | **Dependent variable** | **Compared conditions** | **0.5 – 2 Hz** | **0.5 – 4 Hz** | **0.5 – 10 Hz** |
|---|---|---|---|---|---|
| All participants n = 55 | MI phase-amp | R1 vs R2 (rhyme effect) | V = 618, p-value = 0.2857 | V = 602, p-value = 0.228 | V = 540, p-value = 0.1763 |
| | | Familiarity effect | V = 807, p-value = 0.5816 | V = 822, p-value = 0.4964 | V = 841, p-value = 0.1677 |
| | | Gr 1 vs Gr 2 (group effect) | W = 1470, p-value = 0.8553 | W = 1476, p-value = 0.8263 | W = 1349, p-value = 0.9244 |
| Group 1, n = 27 | MI phase-amp | Familiarity effect | V = 133, p-value = 0.1855 | V = 127, p-value = 0.1414 | V = 121, p-value = 0.1056 |
| Group 2, n = 30 | MI phase-amp | Familiarity effect | V = 219, p-value = 0.7922 | V = 217, p-value = 0.7611 | V = 218, p-value = 0.7766 |
| All participants n = 58 | MI phase-amp | Language effect | V = 864, p-value = 0.9506 | V = 868, p-value = 0.926 | V = 874, p-value = 0.8892 |
| Group 1, n = 27 | MI phase-amp | Language effect | V = 185, p-value = 0.9341 | V = 184, p-value = 0.9153 | V = 186, p-value = 0.9529 |
| Group 2, n = 31 | MI phase-amp | Language effect | V = 261, p-value = 0.8092 | V = 264, p-value = 0.7646 | V = 262, p-value = 0.7942 |
| All participants, n = 56 | MI phase-amp | Rhythm effect | V = 791, p-value = 0.9577 | V = 791, p-value = 0.9577 | V = 734, p-value = 0.6045 |
| Group 1, n = 26 | MI phase-amp | Rhythm effect | V = 188, p-value = 0.7644 | V = 185, p-value = 0.8222 | V = 177, p-value = 0.9801 |
| Group 2, n = 30 | MI phase-amp | Rhythm effect | V = 213, p-value = 0.7 | V = 216, p-value = 0.7457 | V = 195, p-value = 0.4522 |

| | | | | | |
|---|---|---|---|---|---|
| All participants, n = 30 | MI phase-amp | Phonological effect | V = 762, p-value = 0.9499 | V = 768, p-value = 0.99 | V = 745, p-value = 0.8374 |
| Group 1, n = 25 | MI phase-amp | Phonological effect | V = 170, p-value = 0.8532 | V = 170, p-value = 0.8532 | V = 163, p-value = 1 |
| Group 2, n = 14 | MI phase-amp | Phonological effect | V = 227, p-value = 0.9193 | V = 228, p-value = 0.9354 | V = 222, p-value = 0.8394 |

##### 2.4.3. Phase-phase

Due to two extreme outliers in condition 5 (original rhyme 2), we first removed the outliers 3SD above the mean when comparing these two conditions (rhyme effect, familiarity effect and group effect). The analysis of MI values averaged over 0.5 – 2 Hz, 0.5-4Hz and 0.5-10 Hz showed a trend for a rhyme effect in group 1, with higher MI values for the unfamiliar rhyme. This trend spread to the whole sample of participants when averaging over the 0.5-10Hz band, forming a trend of familiarity effect, with higher MI for the unfamiliar rhyme (Fig S26, Table S17).

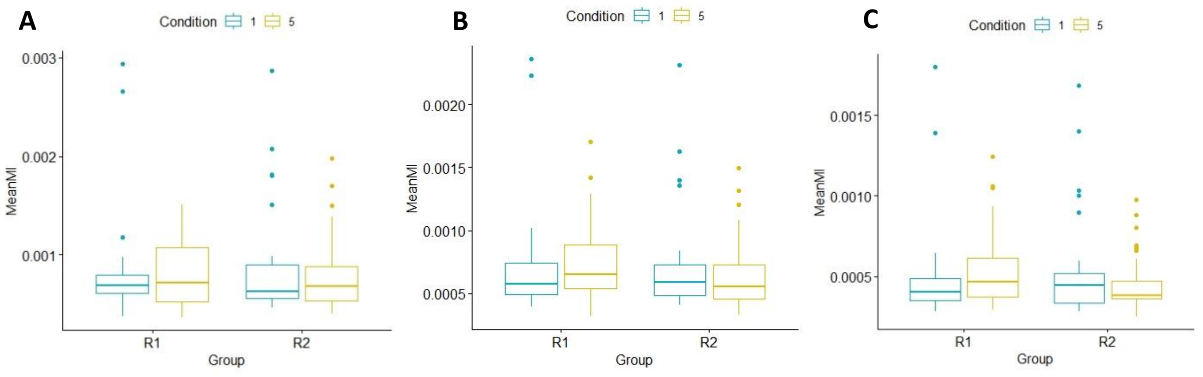

**Fig. S26: Mutual information phase-phase** averaged over 0.5-2Hz (A), 0.5-4Hz (B) and 0.5-10Hz (C).

**Table S17: Results of MI phase-phase analysis**

| **Data** | **Dependent variable** | **Compared conditions** | **0.5 – 2 Hz** | **0.5 – 4 Hz** | **0.5 – 10 Hz** |
|---|---|---|---|---|---|
| All participants n = 55 | MI phase-phase | R1 vs R2 (rhyme effect) | V = 685, p-value = 0.6236 | V = 714, p-value = 0.8095 | V = 710, p-value = 0.7829 |
| | | Familiarity effect | V = 882, p-value = 0.2314 | V = 925, p-value = 0.1171 | V = 943, p-value = 0.08506 |
| | | Gr 1 vs Gr 2 (group effect) | W = 1514, p-value = 0.6956 | W = 1540, p-value = 0.5813 | W = 1599, p-value = 0.3602 |
| Group 1, n = 27 | MI phase-phase | Familiarity effect | V = 115, p-value = 0.07723 | V = 112, p-value = 0.06548 | V = 110, p-value = 0.05847 |
| Group 2, n = 30 | MI phase-phase | Familiarity effect | V = 238, p-value = 0.9193 | V = 259, p-value = 0.5978 | V = 264, p-value = 0.5291 |

| | | | | | |
|---|---|---|---|---|---|
| All participants n = 58 | MI phase-phase | Language effect | V = 929, p-value = 0.5719 | V = 888, p-value = 0.8043 | V = 858, p-value = 0.9876 |
| Group 1, n = 27 | MI phase-phase | Language effect | V = 175, p-value = 0.7496 | V = 167, p-value = 0.6109 | V = 172, p-value = 0.6964 |
| Group 2, n = 31 | MI phase-phase | Language effect | V = 305, p-value = 0.2724 | V = 290, p-value = 0.4214 | V = 266, p-value = 0.7352 |
| All participants, n = 56 | MI phase-phase | Rhythm effect | V = 810, p-value = 0.9253 | V = 851, p-value = 0.6685 | V = 856, p-value = 0.639 |
| Group 1, n = 26 | MI phase-phase | Rhythm effect | V = 189, p-value = 0.7454 | V = 203, p-value = 0.4992 | V = 209, p-value = 0.4082 |
| Group 2, n = 30 | MI phase-phase | Rhythm effect | V = 224, p-value = 0.8712 | V = 228, p-value = 0.9354 | V = 226, p-value = 0.9032 |
| All participants, n = 30 | MI phase-phase | Phonological effect | V = 710, p-value = 0.6181 | V = 645, p-value = 0.2969 | V = 619, p-value = 0.2073 |
| Group 1, n = 25 | MI phase-phase | Phonological effect | V = 171, p-value = 0.8325 | V = 144, p-value = 0.6338 | V = 143, p-value = 0.615 |
| Group 2, n = 14 | MI phase-phase | Phonological effect | V = 205, p-value = 0.5838 | V = 192, p-value = 0.4161 | V = 182, p-value = 0.3085 |

##### 2.4.4. Amp-phase

Due to two extreme outliers in condition 5 (original rhyme 2), we first removed the outliers 3SD above the mean when comparing these two conditions (rhyme effect, familiarity effect and group effect). The analysis of MI values averaged over 0.5 – 2 Hz, 0.5-4Hz and 0.5-10 Hz showed a trend for a rhyme effect with higher MI values for the second rhyme, and in Group 1 this effect reached significance (rhyme 2 being there unfamiliar). There was also a language effect in 0.5-2Hz, with higher MI values for the original rhyme than the language manipulated rhyme. This language effect became just a trend in 0.5-4Hz and was not significant over higher frequencies (0.5-10Hz).

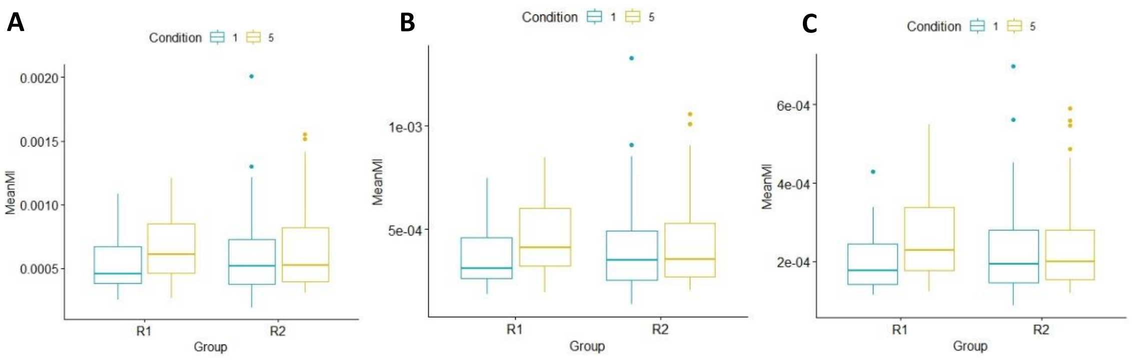

**Fig. 27: Mutual information amplitude-phase**, averaged over 0.5-2Hz (A), 0.5-4Hz (B), 0.5-10Hz (C).

**Table S18: Results of MI amplitude-phase analysis**

| Data | Dependent variable | Compared conditions | 0.5 – 2 Hz | 0.5 – 4 Hz | 0.5 – 10 Hz |
|---|---|---|---|---|---|
| All participants n = 55 | MI amp-phase | R1 vs R2 (rhyme effect) | V = 457, p-value = 0.05407 | V = 466, p-value = 0.06549 | V = 476, p-value = 0.08044 |
| | | Familiarity effect | V = 715, p-value = 0.6293 | V = 743, p-value = 0.4562 | V = 766, p-value = 0.3367 |
| | | Gr 1 vs Gr 2 (group effect) | W = 1280, p-value = 0.9598 | W = 1288, p-value = 1 | W = 1284, p-value = 0.9812 |
| Group 1, n = 27 | MI amp-phase | Familiarity effect | V = 104, p-value = 0.04098 | V = 101, p-value = 0.03399 | V = 107, p-value = 0.0491 |
| Group 2, n = 30 | MI amp-phase | Familiarity effect | V = 189, p-value = 0.3818 | V = 195, p-value = 0.4522 | V = 209, p-value = 0.6408 |
| All participants n = 58 | MI amp-phase | Language effect | V = 1132, p-value = 0.03261 | V = 1103, p-value = 0.05583 | V = 1065, p-value = 0.1056 |
| Group 1, n = 27 | MI amp-phase | Language effect | V = 248, p-value = 0.1624 | V = 244, p-value = 0.1937 | V = 221, p-value = 0.4553 |
| Group 2, n = 31 | MI amp-phase | Language effect | V = 329, p-value = 0.1156 | V = 324, p-value = 0.1406 | V = 322, p-value = 0.1517 |
| All participants, n = 56 | MI amp-phase | Rhythm effect | V = 826, p-value = 0.8225 | V = 800, p-value = 0.9902 | V = 761, p-value = 0.7659 |
| Group 1, n = 26 | MI amp-phase | Rhythm effect | V = 189, p-value = 0.7454 | V = 194, p-value = 0.6528 | V = 176, p-value = 1 |
| Group 2, n = 30 | MI amp-phase | Rhythm effect | V = 229, p-value = 0.9515 | V = 213, p-value = 0.7 | V = 208, p-value = 0.6263 |
| All participants, n = 30 | MI amp-phase | Phonological effect | V = 784, p-value = 0.9099 | V = 780, p-value = 0.9366 | V = 798, p-value = 0.8178 |
| Group 1, n = 25 | MI amp-phase | Phonological effect | V = 160, p-value = 0.9578 | V = 164, p-value = 0.9789 | V = 178, p-value = 0.6915 |
| Group 2, n = 14 | MI amp-phase | Phonological effect | V = 242, p-value = 0.8553 | V = 234, p-value = 0.9838 | V = 235, p-value = 0.9677 |
